# Galectin-3 is a Nanotherapeutic Target in Graft-versus-Host Disease Mediated Kidney Injury

**DOI:** 10.1101/2025.11.19.688972

**Authors:** Magdalini Panagiotakopoulou, Ashley N. Sullivan, Kimon V. Argyropoulos, Anastasia I. Kousa, Logan R. Hillger, Kaitlyn Pierpont, Jorge Nunez, Eric Rosiek, Francesca Mazzoni, Wenfei Kang, Emma Grabarnik, Anastasiya Egorova, Brianna Gipson, Romina Ghale, Stephen Ruiz, Surya V. Seshan, Thangamani Muthukumar, Miguel-Angel Perales, Alan M. Hanash, Marcel R. M. Van Den Brink, Edgar A. Jaimes, Daniel A. Heller

## Abstract

Kidney injury is a frequent and serious complication of allogeneic hematopoietic cell transplantation (HCT), yet its pathophysiology remains poorly understood and effective treatments are lacking. Through analysis of kidney tissue from HCT recipients, we identified substantial acute tubular injury and T cell infiltration that correlated with extra-renal graft-versus-host disease (GVHD) severity, linking systemic alloreactivity and post-transplant renal pathology. In murine models, GVHD was similarly associated with acute kidney injury, renal Th1-type T cell infiltration, and hyperactivation of NF-κB and JAK-STAT signalling. Notably, galectin-3, a damage-associated lectin, was upregulated in both patient biopsies and experimental GVHD target organs. Leveraging this pathological feature, we engineered galectin-3-targeted lipid nanoparticles for tissue-specific delivery of ruxolitinib, an approved GVHD therapy. Galectin-3 upregulation was also identified in canonical acute GVHD target organs including the liver and intestinal tract, and nanoparticle-delivered ruxolitinib substantially enhanced renal function, reduced systemic GVHD, and minimized hematologic toxicity compared to conventional drug administration. Our findings demonstrate renal involvement in acute GVHD and establish a nanoparticle-based strategy for precision delivery of immunomodulatory therapies to affected tissues.

## Introduction

Allogeneic hematopoietic cell transplantation (allo-HCT) is a highly effective treatment for a variety of benign and malignant hematologic disorders. However, it is also associated with graft-versus host disease (GVHD), a condition where T cells in the allograft attack the recipient’s tissues -such as the gastrointestinal tract, skin and liver- and which is associated with poor prognosis^1, 2^. Nearly 50,000 allo-HCT procedures are currently performed annually worldwide, and the number continues to increase by 10% each year. GVHD occurs in up to 50% of patients receiving HCT from an (HLA)-matched sibling and up to 80% in unmatched donors^3^ and carries an approximate 40% mortality rate for chronic GVHD^4^ and up to 70% for acute GVHD^5^.

Acute kidney injury (AKI) is a common complication after HCT, with a reported incidence rate of up to 80%^6, 7, 8, 9, 10, 11, 12, 13, 14, 15, 16, 17^and is linked to substantial non-relapse morbidity and mortality in HCT recipients. Early onset of AKI in HCT patients is linked to higher mortality, highlighting the need for early intervention^18^. In a recent clinical study, we found that the incidence of acute kidney injury (AKI) was 64%, chronic kidney disease (CKD) developed in 21% of these patients, and AKI was linked to a 2.77 higher hazard ratio of non-relapse mortality^19^. Among patients with kidney injury who require dialysis, mortality reaches 100%^15, 20, 21^. Thus, a better understanding of kidney injury post-HCT that may lead to new approaches is urgently needed to improve outcomes for HCT patients.

The etiology of kidney injury after HCT is diverse, including volume depletion, infections, drug toxicities, thrombotic microangiopathy, glomerular diseases (i.e. membranous nephropathy), and tubulointerstitial injury^22^. Although the etiology of AKI in HCT is multifactorial, and the kidney is not considered a traditional organ-target of GVHD, as are the gastrointestinal tract, skin and liver, growing evidence suggests that alloreactive immune responses may themselves directly contribute to renal injury after allo-HCT. Retrospective clinical studies have shown that aGVHD is a risk factor for AKI^11, 23, 24, 25, 26^, and there are some case studies suggesting the existence of kidney-GVHD^27, 28^. However, kidney injury in patients after HCT remains poorly characterized due in part to the paucity of renal biopsies in these patients given the higher risk of complications, in particular bleeding. As such, kidney biopsies are rarely performed in patients with acute GVHD, and clinical or pathologic criteria for renal involvement in acute GVHD have not yet been established^29^. This creates a gap in existing knowledge around the diagnostic features of this type of injury after HCT. Preclinical models of HCT show features of immune mediated kidney injury; however, the progression of kidney injury, the nature of immune cell subpopulations, and changes in gene expression are not heretofore well-characterized^30, 31, 32^.

Clinical management of GVHD can be challenging; immuno-suppression with corticosteroids constitutes the first-line therapy in both acute (aGVHD) and chronic GVHD (cGVHD), but steroid treatment can also result in severe toxicities. Moreover, responses to steroids are maintained in less than 50% of patients with aGVHD and only 40–50% of patients with cGVHD in some series^33, 34, 35^. The JAK inhibitor ruxolitinib is the sole FDA-approved therapy for steroid refractory aGVHD, and its use is limited by substantial hematologic toxicities, including thrombocytopenia and anemia^36, 37^ as well as an increased risk of serious bacterial, fungal, mycobacterial, and viral infections, including tuberculosis, herpes zoster reactivation, and progressive multifocal leukoencephalopathy^38^. Broader concerns about the JAK inhibitor class, reflected in FDA boxed warnings for tofacitinib, baricitinib, and upadacitinib regarding major adverse cardiovascular events, malignancy, thrombosis, and mortality^39^, further motivate strategies to restrict systemic exposure while preserving on-target activity at the site of disease. Salvage therapies beyond the second-line remain suboptimal^40^. Regarding the kidneys, drug delivery strategies to target therapies to the kidneys constitute a clear unmet need^41, 42^. There are currently no specific therapeutics available for the treatment of acute kidney injury (AKI) of any etiology and most of the care is supportive. Few drugs are available for the treatment of chronic kidney disease, largely due to their poor pharmacokinetic properties^43, 44^. Nanotherapeutic drug delivery strategies to improve pharmacokinetics of drugs in the kidneys remain under-investigated, and few have succeeded in preclinical stages^45, 46, 47^. Given the current management of AKI post allo-HCT remains supportive for the large majority of patients, nanotherapeutic delivery of drugs with an established efficacy in GVHD may provide a novel therapeutic approach^48^.

Here, we provide evidence linking human AKI to aGVHD and identify a novel therapeutic strategy for improving renal function and reducing systemic aGVHD after HCT. Comprehensive analysis of kidney biopsies from post-HCT patients, collected within 14 months after transplantation revealed substantial acute tubular injury and extensive T cell infiltration, particularly within the epithelial compartment, that showed a striking positive correlation with the severity of extra-renal GVHD. Building on these observations, we investigated the pathobiology and molecular signatures of AKI post-transplant in an MHC-disparate mouse model of GVHD. Mice displayed elevated serum and urine markers of kidney injury as well as kidney function impairment proportionate to the severity of GVHD. We uncovered extensive upregulation of inflammatory signaling pathways that is consistent with the identification of tubulitis and glomerulitis. We also performed a comprehensive immune cell repertoire analysis in the infiltrated T cells and found substantial CD8^+^ and Tbet^+^ enrichment. We also observed overexpression of pro-inflammatory regulators galectin-3 and P-selectin in GVHD-AKI mouse kidneys. Remarkably, these findings were validated in our patient biopsies, which showed increased endothelial P-selectin expression and elevated galectin-3 across multiple kidney compartments, particularly in patients with acute GVHD. Upregulation of these targets led to increased accumulation of targeted nanoparticles in the kidneys of GVHD mice. Using a lipid nanoparticle drug delivery system, we tested various small molecule inhibitors of overregulated pathways identified by transcriptome analysis and discovered and validated JAK/STAT as a therapeutic target for early treatment of GVHD-AKI. In mice with severe GVHD, nanoparticle-mediated ruxolitinib delivery was therapeutically superior to unencapsulated drug controls. Additionally, targeted delivery of ruxolitinib successfully reversed the manifestations of GVHD-AKI and mitigated hematologic toxicity of ruxolitinib. Our findings suggest that a substantial inflammatory component in GVHD-AKI kidneys results in JAK/STAT pathway activation leading to kidney injury which can be addressed effectively by galectin-3-localized JAK/STAT pathway inhibition.

## Results

### Patients with acute extra-renal GVHD exhibit GVHD-consistent kidney pathology

We studied the pathobiology of kidney injury in renal biopsies collected from post-HCT patients. We collected and analyzed early-timepoint kidney tissues from 29 BMT recipients at MSKCC who underwent a kidney biopsy within the first 14 months after allo-HCT (**Fig. 1a**). Needle-core biopsies were collected between 2010 and 2024, with biopsy indications including acute kidney injury and/or nephrotic or non-nephrotic range proteinuria. Eight patients with BK virus (BKV) nephropathy were excluded to prevent misattribution of BKV-caused acute interstitial nephritis to GVHD, and one biopsy obtained in a perimortem setting (8 days prior to death) met exclusion criteria due to terminal bacterial infection.

**Fig. 1.**
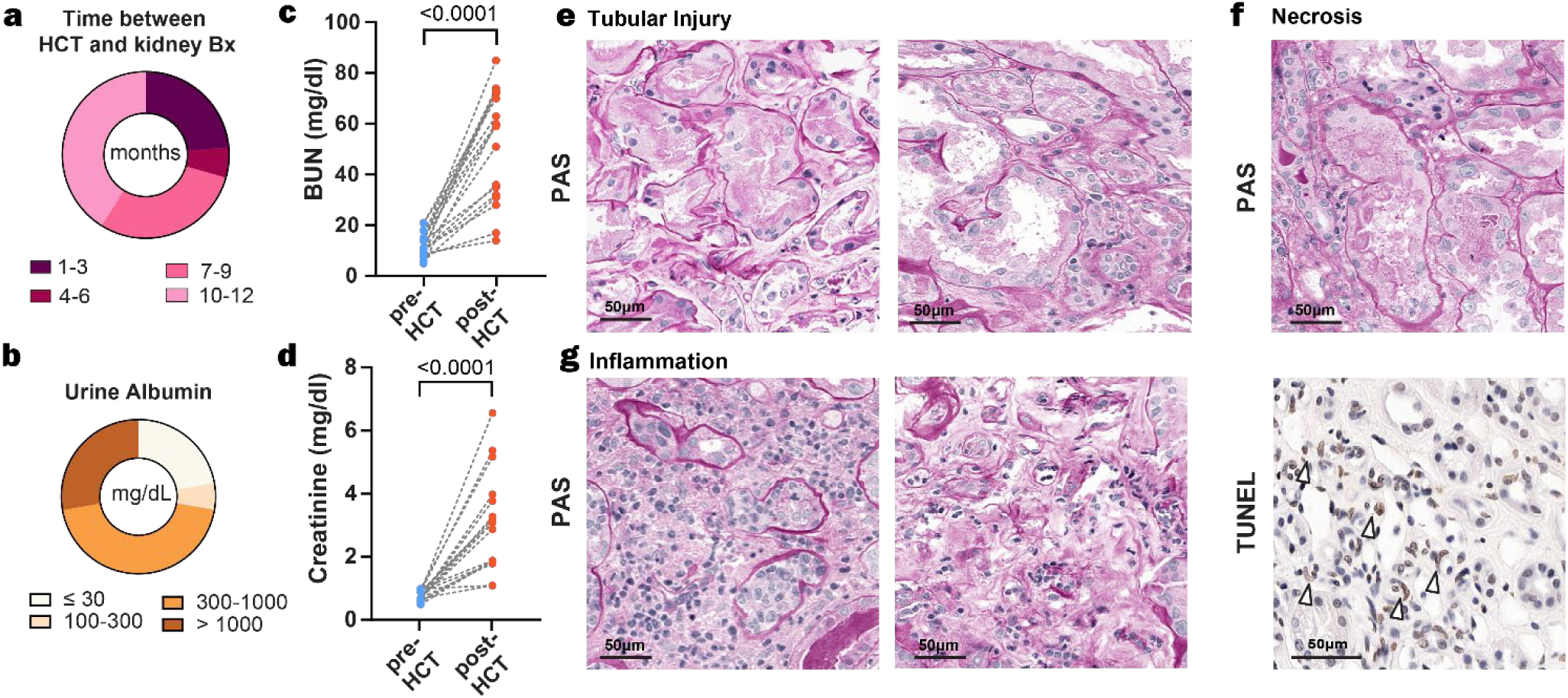
Renal function, T cell localization, and tissue injury in BMT patients. **a**, Time (months) between HCT and kidney biopsy. **b**, Urinary albumin excretion levels post-HCT. **c**, BUN levels pre- and post- HCT, Wilcoxon paired test. **d**, Serum creatinine levels pre- and post-HCT, Wilcoxon paired test. **e-f**, Representative PAS staining of tubular injury and inflammation in post- HCT kidneys. **g**, PAS and TUNEL staining of post-HCT kidney tissues exhibiting necrosis. For **a-d**: n=20 (pre- and post-HCT). For **c,d,** Wilcoxon matched-pairs signed rank test. Bx = biopsy

Demographic and disease/treatment information on the remaining 20 patients is summarized in **Supplementary Table 1**. The most frequent indications for allo-HCT in these patients were Acute Myeloid Leukemia (AML) and Acute Lymphoblastic Leukemia (ALL). 47% of the patients were conditioned with total body irradiation (TBI) and chemotherapy, the other 53% with chemotherapy only. Because renal biopsies are performed based on clinical indication and are not temporally synchronized with peak systemic GVHD activity, we annotated both GVHD status at the time of biopsy and prior history of GVHD. Of the 20 patients studied, 17 were diagnosed clinically with GVHD at some point prior to the biopsy. At the time of the kidney biopsy, 6 of these patients had active acute GVHD, 4 had chronic GVHD (all with prior acute GVHD diagnoses), and one patient had overlap syndrome as classified by NIH consensus criteria that assess disease features beyond temporal onset^49^. Three additional patients presented GVHD-like manifestations but failed to meet criteria for formal clinical diagnosis (hereafter referred to as “indeterminate”). Five patients had received T cell depleted transplants which is known to reduce severe acute GVHD incidence by up to 70%^50, 51^. Of the 11 confirmed GVHD cases, 4 were on calcineurin inhibitors for GVHD prophylaxis at the time of renal biopsy (**Supplementary Table 1**). A full list of medications for all patients is presented in **Supplementary Table 2** and the pathological findings are summarized in **Supplementary Table 3**. All patients had normal kidney function at baseline before allo-HCT (**Fig. 1c-d**). When stratified by extra-renal GVHD status, patients with acute GVHD showed significant increases in both BUN and serum creatinine after HCT, whereas patients without GVHD showed a modest increase in creatinine but not BUN. Chronic GVHD cases showed numerical increases in both parameters that did not reach statistical significance, likely reflecting the small sample size in this subgroup (n = 4) **(Supplementary Fig. 1**).

With regard to histologic findings of the kidney parenchyma, following allo-HCT, 14 of 20 patients exhibited acute tubular injury (**Fig. 1e**), with 7 cases showing frank acute tubular necrosis (**Fig. 1f**). Moreover, 13 of 20 showed global glomerulosclerosis, which may be associated with aging or with other underlying kidney pathologies (**Supplementary Fig. 2**). In terms of pathologic findings in the vasculature, thrombotic microangiopathy (TMA) was observed in 5 of 20 patients (25%) - comprising 3 chronic GVHD, 1 indeterminate, and 1 non-GVHD patient. Separately, chronic endothelial injury was noted in 1 patient with chronic GVHD (**Supplementary Table 3**). Arteriosclerosis and arteriolosclerosis were consistently observed in 90% of patients, likely reflecting age-related changes and possibly underlying comorbidities such as hypertension. Most importantly, with regard to renal interstitial tissue, 85% of the kidney biopsies had either mild or moderate inflammatory infiltrates (**Fig. 1g)**, which prompted us to investigate the nature and localization of these infiltrates. Three patients had findings consistent with membranous nephropathy while five patients had minimal change disease.

Since T cells are the major mediators driving GVHD pathology we sought to quantify CD3^+^ T cell infiltration in patient renal biopsies to determine if alloimmune inflammation directly contributes to post-HCT acute kidney injury. As controls, we used: 1) grossly and microscopically normal kidney sections that had been routinely sampled in the context of pathologic work-up performed in nephrectomy specimen for renal tumors, 2) kidney sections from biopsies of 6 patients who did not undergo HCT and presented with kidney injury of unrelated etiologies (hereafter denoted “Other etiologies”). The diagnosis of those 6 patients is described in **Supplementary Table 4**. Our analysis revealed that post-HCT kidneys exhibited significantly increased CD3^+^ T cell infiltration compared to both normal and non-HCT kidney injury controls (p<0.0001) (**Fig. 2a**), which indicates that the degree of renal T cell infiltration observed in GVHD is not a nonspecific feature of kidney injury or malignancy, but is enriched in the alloimmune GVHD setting.

**Fig. 2.**
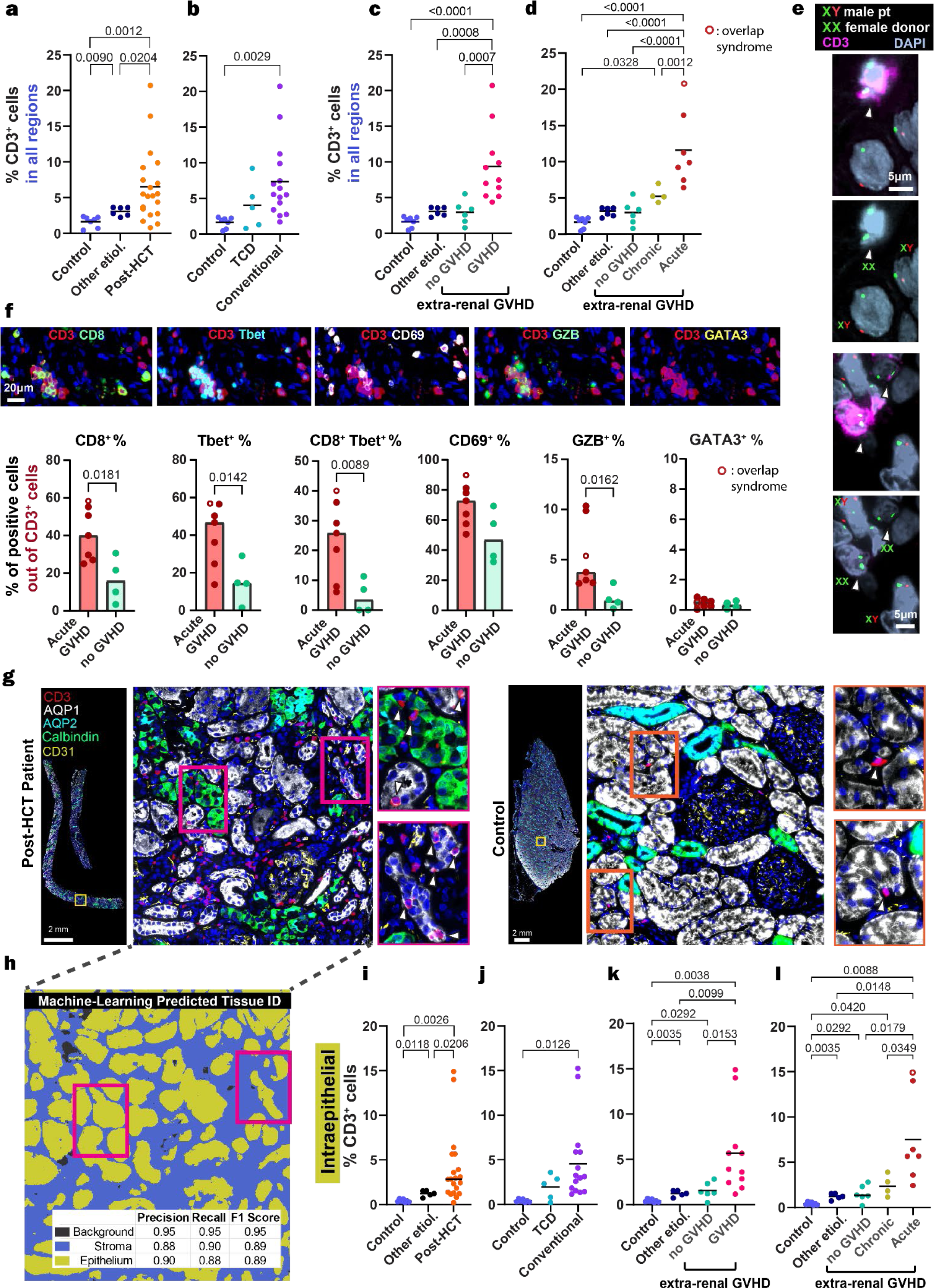
a-d, Spatial and functional characterization of alloimmune T cells in post-HCT human kidneys. **a-d**,Percentage of CD3^+^ cells in all regions comparing control groups against post-HCT patients, T cell depletion (TCD), and conventional transplant recipients, and various extra-renal GVHD manifestations **e**, DNA X/Y fish in sex-mismatch patients with acute GVHD, showing examples of XX (female) CD3^+^ cells infiltrating the tissue of the XY (male) recipient. **f,** Multiplexed immunofluorescence and quantification of T cells positive for various activation and polarization markers **g**, Multiplexed immunofluorescence illustrating tissue localization of CD3^+^ T cells. Immunofluorescence markers for kidney compartments: calbindin (distal tubules), CD31 (endothelium), Aquaporin 1 (proximal tubules), and Aquaporin 2 (collecting ducts), DAPI (nuclei) **h**, Machine learning-based calculation of tissue identity from panel g, with precision, recall and F1 score: background, stroma, and epithelium classes in post-HCT patients. **i-l**, Percentage of CD3^+^ cells in epithelium comparing control groups against post-HCT patients, TCD and regular transplant recipients, and various extra-renal GVHD manifestations. For **a-d, i-l**: n_contro_l=6, n_other etiologies_ = 6, n_post-HCT_=20. **f**: Welch’s t-test. **a-d, i-l**: Ordinary one-way ANOVA with Tukey correction. Hollow points indicate patients with overlap syndrome. **a-d, i-l**: Line represents the mean.

Post-HCT kidneys from patients receiving conventional, T cell replete transplants exhibited significantly increased overall and intraepithelial CD3^+^ T cell infiltration compared to normal controls (**Fig. 2b**). In contrast, infiltration in T cell depleted (TCD) recipients did not significantly differ from controls. This selective elevation in the conventional cohort indicates the allograft is the primary source of T cells driving renal inflammation. The severity of extra-renal GVHD (absent to chronic and acute/overlap) showed a positive correlation with T cell infiltration in patient kidneys (**Fig. 2c, d and Supplementary Fig. 3a**), indicating a direct link between extra-renal and renal GVHD.

To verify the origin of kidney-infiltrating T cells, we combined CD3 staining with fluorescence *in situ* hybridization (FISH) for XY chromosomes in four sex-mismatched transplant recipients. Due to the high magnification (100×) required to resolve individual FISH probes, field-of-view limitations restrict bulk tissue quantification. Nevertheless, high-resolution imaging consistently revealed multiple donor-derived XX T cells actively infiltrating the XY recipient tissue (**Fig. 2e**), confirming the allograft as the source of the immune infiltrate.

To verify the polarization and activity of T cells, we conducted multiplexed immunofluorescence analyses on human kidney biopsies utilizing a dedicated T cell activation panel (**Fig. 2f**). Compared to patients without GVHD, kidneys from patients with acute GVHD consistently exhibited a significant enrichment of CD8+ T cells (40.85% vs. 16.42%, p = 0.0181), which were highly activated and polarized towards a cytotoxic Th1 phenotype. Specifically, infiltrating cells in the acute GVHD cohort showed significant co-expression of T-bet (40.70% vs. 14.86%, p = 0.0142), a marked increase in CD8+ T-bet+ double-positive cells (23.64% vs. 4.58%, p = 0.0089), and elevated granzyme B (5.37% vs. 1.14%, p = 0.0162). Conversely, expression of the Th2 master regulator GATA3 was virtually absent in these infiltrates, definitively confirming a Th1-polarized response. We also observed elevated expression of the activation marker CD69 in the acute GVHD cohort (70.56% vs. 48.78%; p = 0.0855). This profile confirms their active role in mediating cytotoxic alloimmune tissue injury, rather than representing a non-specific inflammatory infiltrate. Consistent with this cytotoxic profile, serial section analysis of post-HCT kidney biopsies revealed TUNEL-positive apoptotic cells in regions corresponding to dense CD3+ T-cell infiltration, with adjacent PAS sections confirming tubular injury in the same areas (**Supplementary Fig. 4a**), providing morphological evidence consistent with T-cell-mediated epithelial injury in these regions.

Finally, to assess the localization of T cells we performed multiplexed immunofluorescence (IF) staining of kidney biopsies derived from HCT recipients. In post-HCT and control sections, we labeled T cells (CD3^+^), proximal tubular cells (Aquaporin 1(AQP1)^+^), distal tubular cells (calbindin (Calbindin^+^), collecting duct (Aquaporin-2(AQP2)^+^)), connecting tubule (Aqp1^+^Aqp2^+^), and endothelial cells (CD31^+^) (**Fig. 2g**). To classify kidney tissue sections as stroma or epithelium, we employed a machine learning-assisted pixel classifier algorithm based on the specific marker expression, where cells positive for epithelial markers (AQP1, CALB, AQP2) were classified as epithelium, with additional validation and refinement from a pathologist (see Methods) (**Fig. 2h and Supplementary Fig. 4a**). We then quantified CD3^+^ cells in the epithelium and stroma regions of interest (% positive = amount of CD3^+^ cells/ total cells in epithelium or stroma); we surmise that the cells in the epithelium are potential effector T cells, while those in the stroma non-resident or circulating T cells^52, 53^.

Our analysis revealed that the increased CD3^+^ T cell infiltration in post-HCT patients was equally pronounced within the proximal and distal tubular epithelial compartment (**Fig. 2i**). The marked accumulation of T cells in the epithelium suggests immune-mediated epithelial injury, consistent with reports where T cell localization within epithelial tissues has been implicated in tissue injury in various inflammatory conditions, including skin GVHD and celiac disease^54, 55, 56, 57^. The severity of extra-renal GVHD (absent to chronic, and acute/overlap) showed again a positive correlation with T cell infiltration in epithelium (**Fig. 2k,l and Supplementary Fig. 3b**), indicating a direct link between extra-renal and renal GVHD.

Interestingly, in post-HCT patients, we observed a marked upregulation of AQP1 in the glomeruli with a concomitant reduction in AQP2 expression in the distal nephron (**Supplementary Fig. 4b**). The increase in glomerular AQP1, primarily located in the proximal tubules with some expression in glomerular endothelium^58^, is consistent with findings from renal diseases where AQP1 upregulation occurs in response to injury^59^. Moreover, the reduction in AQP2, particularly in fibrotic areas, aligns with reports of decreased AQP2 expression in renal diseases associated with nephron loss and fibrosis^60^ and could be associated with the clinical manifestations of AKI in post-HCT patients.

### Allo-HCT induces kidney inflammation and injury in experimental GVHD models

Collectively, our findings in patients support a model in which kidney injury in post-HCT patients is not solely attributable to nephrotoxins or hemodynamic stress but is associated with epithelial-directed alloimmune activity proportional to systemic GVHD severity. This observation provided the rationale for mechanistic interrogation in a controlled preclinical system. To further elucidate whether the pathology observed in patients post-HCT is due to underlying kidney alloreactive inflammation and injury, we employed two established mouse models of GVHD (major and minor HLA mismatch) to monitor the disease time course and identify potential therapeutic targets to abrogate this injury.

For the major mismatch model, we performed allo-HCT in mice by administering 900 cGy of fractionated lethal radiation to BALB/c mice followed by retroorbital injection of 5×10^6^ cells from T cell depleted bone marrow from C57BL/6 mice. One million splenic T cells were then added to the bone marrow preparation and injected intravenously into one group (GVHD group), as previously described^61^, while a second group received bone marrow cells without T cell addition (bone marrow only (BMO) group). For the minor mismatch model, C57BL/6 mice received 1100 cGy split-dose radiation followed by a tail vein injection of 15×10^6^ splenocytes from miHA-disparate 129S1 donor mice.

On the day of harvesting, mice were anesthetized with ketamine/xylazine and then subjected to transcardial perfusion with PBS. We then performed kidney histopathologic analysis to determine the degree and nature of kidney injury. We used PAS stain and conducted IHC for CD3, TUNEL, KIM-1 and NGAL. In the major mismatch model with 1M T cells (hereafter denoted as GVHD (1M)), as early as day 7 post-transplantation, the GVHD group had notable tubular injury including tubular necrosis, loss of brush border, and vacuolization which was identified by blinded analysis by a renal pathologist (**Fig. 3a**). In addition, we observed extensive CD3^+^ cell infiltration in both tubules and glomeruli, apoptosis as assessed by TUNEL, as well as increased KIM-1 and NGAL, both sensitive markers of kidney injury (**Fig. 3a)**^62, 63^. We chose to use NGAL and KIM-1 as injury biomarkers because serum urea and creatinine are not as sensitive in mice as in humans^64^. Moreover, creatinine is affected by mouse muscle mass loss (a feature of systemic GVHD)^65, 66^. In the major mismatch model while NGAL positive cells decreased from day 7 to day 14, CD3^+^ cells increased, indicating continuous activation of the immune system (**Fig. 3b**, and **Supplementary Fig. 5**). We observed a significant increase in NGAL and KIM-1 in serum and urine that correlated with the presence of vacuolized and apoptotic cells, seen by morphology and IHC (**Fig. 3c**).

**Fig. 3.**
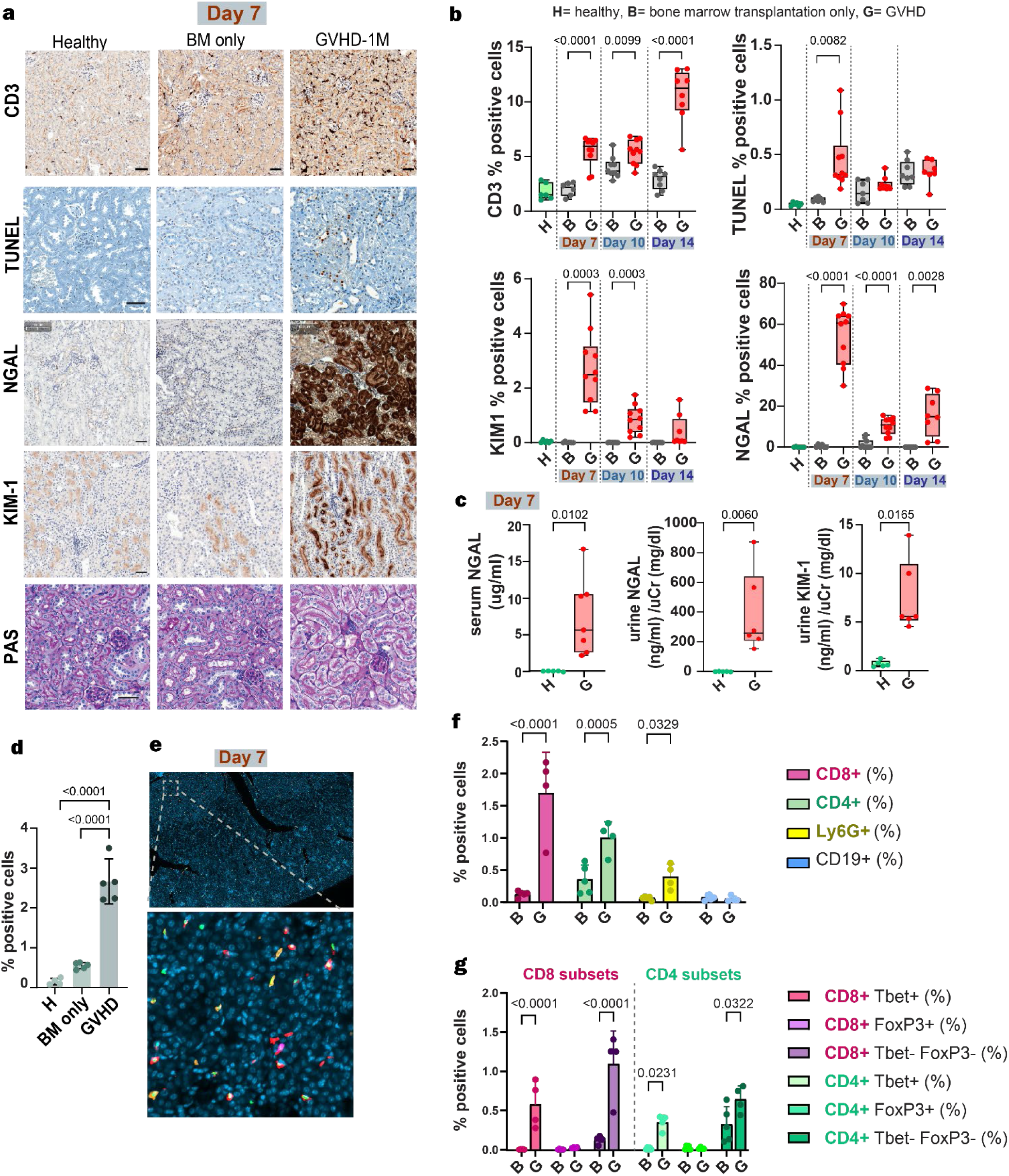
Allo-HCT MHC-disparate murine model of GVHD exhibits pathological and molecular features of AKI. **a**, T cell infiltration (CD3), apoptosis (TUNEL), kidney injury (NGAL, KIM-1) and renal morphology (PAS) on day 7 after bone marrow transplantation. (scale bar: 50um). **b**, Quantification of CD3, TUNEL, KIM-1 and NGAL positive cells over time. n_H_ = 6, n_B_=6-8, n_G_= 8-10, Welch’s t test. **c**, Levels of serum and urine markers of renal damage on day 7 after bone marrow transplantation. n_H_ = 5, n_G_= 6-7, Welch’s t test. **d,** Total number of detected immune cells per condition n_H_ = 4, n_B_=5, n_G_= 5, One-way ANOVA with Tukey correction. **e,** Merged immunofluorescence image, top, (and higher magnification image, bottom) of kidney tissue stained for CD8 (red), Tbet (green), CD4 (orange), FoxP3 (magenta), CD19 (dark blue), Ly6G (yellow) and DAPI (cyan). **f,** Quantification of immune cell subsets identified by multiplex immunofluorescence in each experimental condition. **g**, Quantification of the proportion of T-bet⁺ or FoxP3⁺ cells within CD8⁺ and CD4⁺ T cell populations. n_B_=5, n_G_= 4, Two-way ANOVA with Tukey correction. **b, c**: The box extends from the 25th to 75th percentiles and the whiskers from min to max. Line represents the median. **d,f,g**: Data are presented as mean ± s.d. H = healthy, B = bone marrow transplantation only, G = GVHD.

We repeated the same workflow in the minor HLA mismatch model, evaluating pathology on days 15, 30, and 45 after transplantation (**Supplementary Fig. 6**). Notably, the severity of renal injury in this clinically relevant model was equivalent to that observed in the major mismatch setting. By day 30 post-transplant, mice exhibited markedly elevated levels of the injury biomarkers NGAL and KIM-1, as confirmed by both ELISA and immunohistochemistry. This injury was accompanied by dense CD3^+^ T cell infiltration within the renal parenchyma and concurrent pSTAT3 activation, demonstrating that GVHD-mediated renal pathology is deeply conserved across both major and minor mismatch settings. Since the major mismatch model provides a more rapid, robust, and highly synchronized onset of severe renal pathology, it offers an optimal and reliable therapeutic window for evaluating our targeted nanomedicine interventions. Therefore, we performed the rest of our efficacy analyses using the major mismatch model.

To attain a better understanding of the immune landscape in GVHD kidneys, we characterized the immune cell subpopulations of kidney infiltrates. We performed multiplexed immunofluorescence and computer-aided quantification in whole tissue to determine how immune cell numbers differ from healthy tissue or the tissues of BMO controls (**Fig. 3** and **Supplementary Fig. 7**). Differences were observed in both the total number of infiltrated immune cells, which was approximately tenfold higher in GVHD kidneys as compared to healthy and BM-only controls (**Fig. 3d**) and in the distribution of specific immune subsets. In GVHD kidneys, CD8⁺ T cells were the most abundant immune population, followed by CD4⁺ T helper cells, neutrophils, and B cells (**Fig. 3f**). To further define inflammatory polarization and regulatory balance, we quantified the expression of T-bet and FoxP3 at the single-cell level within CD8⁺ and CD4⁺ cell phenotypes using colocalization analysis (**Fig. 3g**). GVHD kidneys demonstrated a marked increase in T-bet⁺ CD8⁺ effector T cells (p < 0.0001) and T-bet⁺ CD4⁺ Th1-like cells (p = 0.0231), whereas FoxP3⁺ regulatory T cells remained scarce in both compartments. These findings indicate a strong skew toward pro-inflammatory T-bet–driven effector responses with limited regulatory control. The rapid recruitment of CD8^+^ cells to the kidney is consistent with findings in other GVHD target organs, such as the liver, colon, and skin^61^. However, the role of Tbet+ CD4+ cells appears to vary by tissue, with Th1 responses being more prominent in liver and intestinal GVHD, while skin GVHD is often associated with stronger Th2 and Th17 components^67^.

To gain deeper insight into the gene expression profile of GVHD-AKI kidneys and to identify therapeutic targets, we performed bulk sequencing of RNA (RNA seq) isolated from the perfused kidneys 7, 10 and 14 days post-transplantation, from healthy, BMO and GVHD (1M) mice. On day 7, 1542 genes were up-regulated and 1838 genes were down-regulated (log2(fold change)>1) in GVHD mice compared to healthy control (**Fig. 4a**), while 1495 genes were upregulated and 1508 genes were down-regulated in GVHD mice compared to mice receiving bone marrow only.

**Fig. 4.**
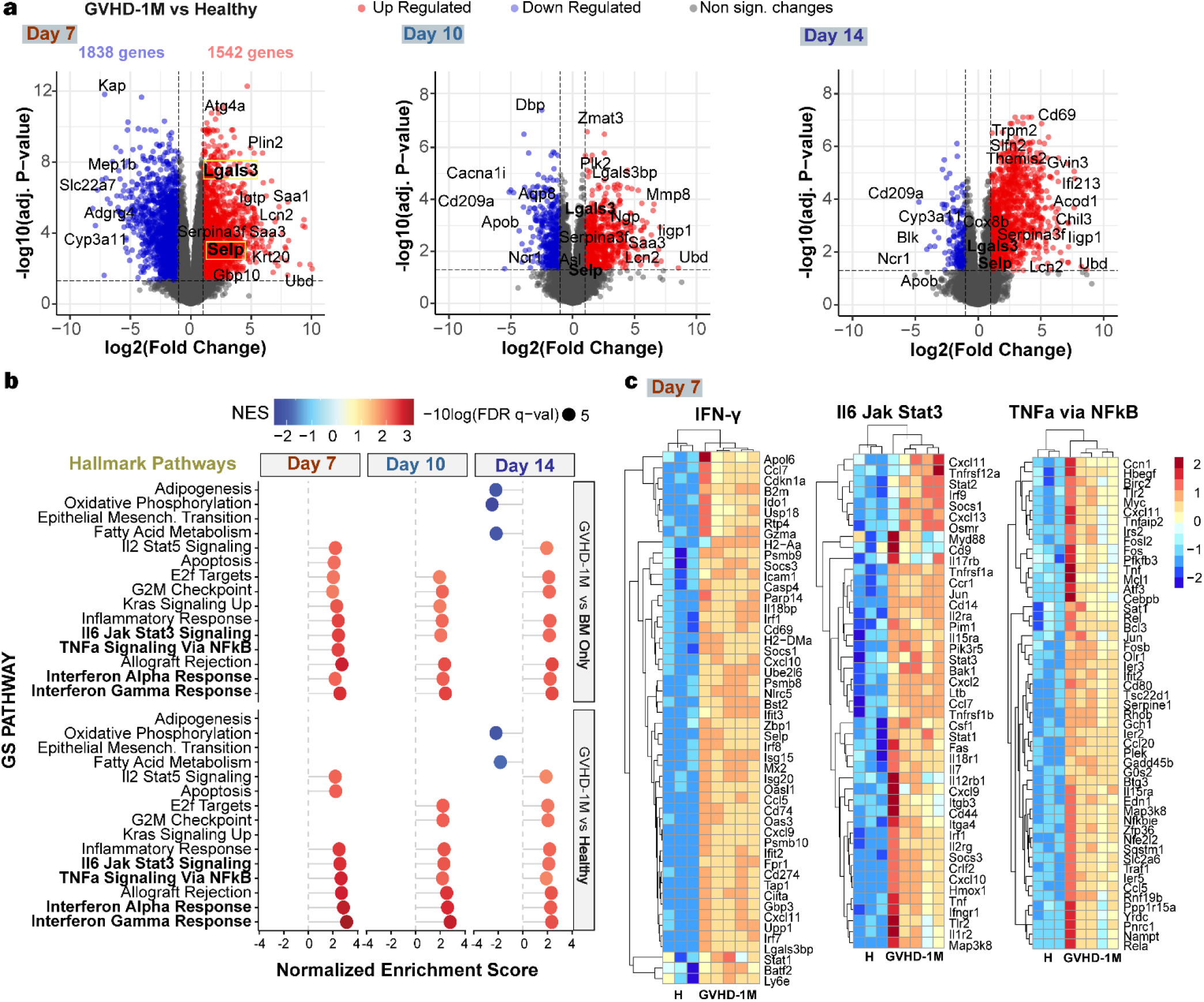
Transcriptome analysis of kidney lysates from GVHD mice in a MHC-disparate model reveals highly dysregulated immune-related pathways. **a,** Volcano plots of genes differentially expressed between GVHD and healthy mice on days 7, 10 and 14 post BMT. **b,** Hallmark pathway analysis of top dysregulated pathways between GVHD, BM only and healthy mice on days 7, 10 and 14 post transplantation. **c,** Leading edge genes contributing to the enrichment of the major pathways chosen for pharmacological inhibition.

We found strong evidence of persistent immune pathway activation, in accordance with data from human kidneys (**Fig. 4a** and **Supplementary Fig. 8)**, as well as significant upregulation (more than 4-fold) of genes associated with the IFN-γ, IFN-α, JAK-STAT3 and NFκB pathways, as indicated by the gene set enrichment analysis (GSEA)^68^ of the aforementioned comparisons against the mouse Hallmark Gene Set database (**Fig. 4b)**.

We also observed a dramatic increase in expression of genes associated with kidney injury (*Ubd* , ubiquitin d, *Lcn2*, NGAL), and pro-fibrotic pathways (*Saa3, Saa1, Chil3*)^69^. Hallmark pathway analysis indicated that the top upregulated pathways included IFN-γ, IFN-α, JAK-STAT3 and NFκB (**Fig. 4b,c)**. JAK-STAT signaling in both innate and adaptive arms of immunity plays a central role in the development and maintenance of GVHD. Specifically, JAK/STAT signaling is active in: (a) tissue damage from disease and conditioning chemoradiotherapy, (b) donor T cell activation, and (c) recruitment and activation of other immune cells^70, 71^. It has also been shown that inflammatory cytokine signaling in tissues can contribute to tissue pathology^72^. NF-κB signaling is also a well-established mediator of the inflammatory response, as it regulates cytokine production, leukocyte recruitment and cell survival^73^, but despite its association with numerous inflammatory diseases, its role in GVHD has only recently been appreciated.

To validate our findings, expression of top candidate genes involved in enriched pathways identified in our RNAseq results were subsequently confirmed by biochemical assays. We performed qPCR on effectors of the dysregulated pathways, including STAT1, STAT3, TNFRSF1A, TLR2, MAP3K8, RELB, SOCS3, NFκB2, TNF, PLIN2, SOCS1 and LIF and we found that the expression of those genes was significantly elevated in the GVHD group compared to control (**Supplementary Fig. 9**). We also confirmed elevation of STAT3, pSTAT3, NFκB and RELB by Western Blot. (**Supplementary Fig. 10**).

### Galectin-3 and P-selectin-targeted nanocarrier localization in GVHD-AKI kidneys

We analyzed the gene expression profiles for evidence of modulation of known drug delivery targets. We found elevated P-selectin (SELP, log_2_fold change = 3.32, adjusted *P*= 2.8E-10), a transmembrane protein normally overexpressed in activated endothelial cells^74, 75^. We also observed substantially enhanced galectin-3 (LGALS3, log_2_fold change = 1.4, adjusted *P* =6E-15), a multifunctional protein with pivotal role in interstitial fibrosis and progression of chronic kidney disease^76^, and galectin-3 binding protein (LGALSBP, log_2_fold change = 3.29, adjusted *P* = 1.7E-55). We confirmed overexpression of both proteins in GVHD murine kidneys by immunofluorescence staining, with P-selectin being expressed by endothelial cells and galectin-3 by tubular epithelial cells (**Fig. 5a**).

**Fig. 5.**
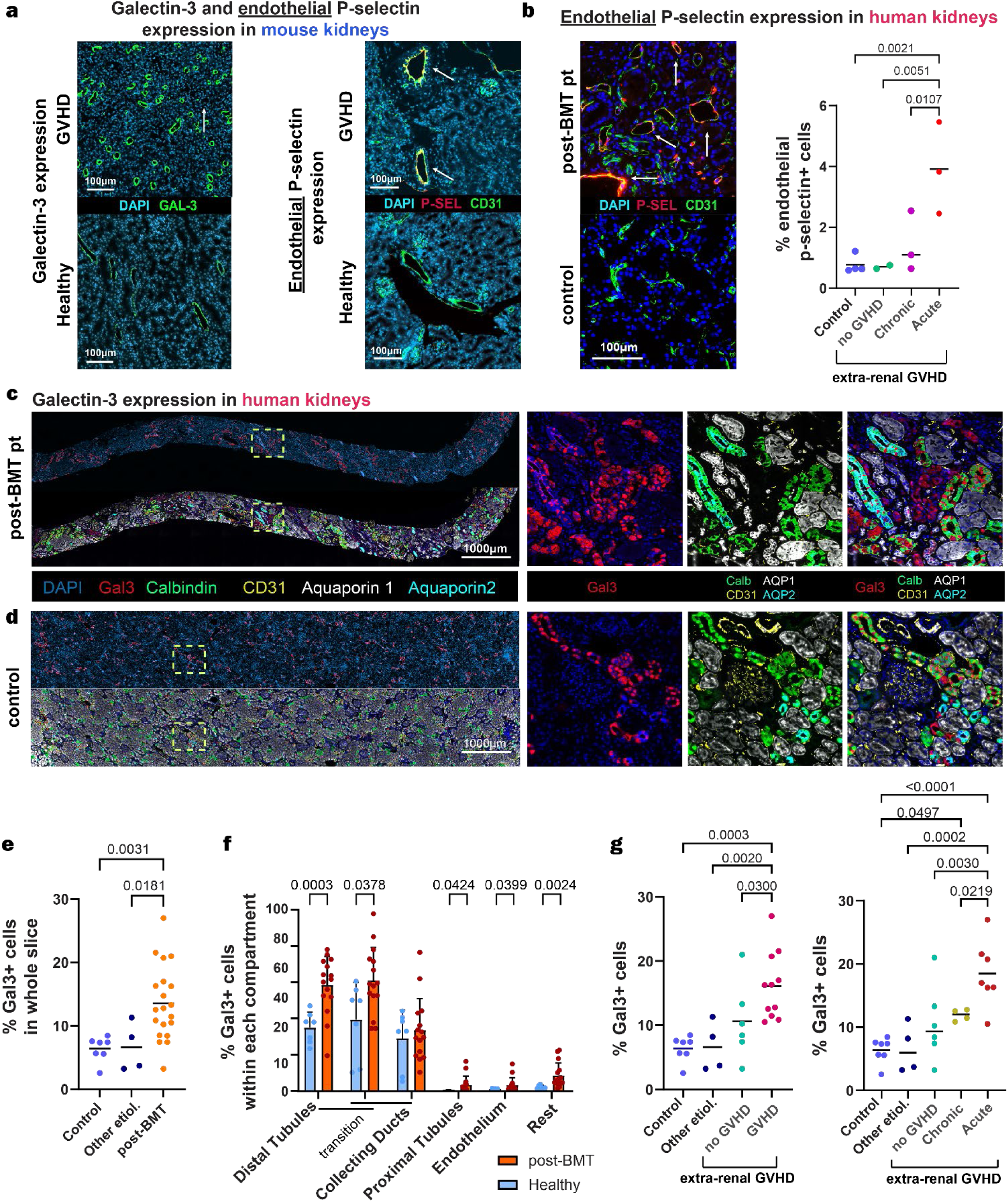
Evaluating nanotherapeutic targets in GVHD-AKI across mouse and human kidneys. **a,** Galectin-3 (left) and endothelial P-selectin (right) expression in healthy vs. GVHD mouse kidney tissues. **b,** Quantification of endothelial P-selectin expression (to exclude expression in blood cells) in human kidney biopsies from post-HCT patients compared to controls, expressed as a percentage of P-selectin^+^CD31^+^ cells in the whole tissue slice. **c, d,** Representative merged immunofluorescence image of galectin-3, Calbindin (CALB), CD31, Aquaporin 1 (AQP1), and Aquaporin 2 (AQP2) markers in post-HCT **(c)** and control **(d)** kidney biopsies, highlighting compartmental localization of galectin-3. **e,** Quantification of galectin-3 expression in human kidney biopsies from post-HCT patients compared to controls, expressed as a percentage of galectin-3^+^ cells in the whole tissue slice **f,** Quantification of galectin-3^+^ cells within specific compartments, showing significant increases in the distal tubules, collecting ducts, and endothelium in post-BMT human kidneys compared to controls. **g,** Quantification of galectin-3+ cells comparing control groups with various GVHD manifestations For **b**: n_post-HCT_=8, n_control_=6. For **e**: n_post-HCT_=7, n_other etiologies_=6, n_control_=6, For **g**: n_post-HCT_=17, n_other etiologies_=4, n_control_=6. **b,e,g**: one-way ANOVA with Tukey correction. Line represents the mean. **f,** unpaired Welch’s t-tests.Data are presented as mean ± s.d.

To evaluate the expression enhancement in other target organs of GVHD, we measured P-selectin and galectin-3 in the liver, small and large intestine, and spleen (**Supplementary Fig. 11)**. We found significant upregulation of galectin-3 in kidney, liver, and small intestine during GVHD, while endothelial P-selectin was significantly increased in kidney and large intestine compared to healthy controls. The spleen showed minimal expression of both proteins with no significant changes during GVHD (**Supplementary Fig. 11**).

To examine the distribution of P-selectin and galectin-3 in human kidneys and its potential as a therapeutic target in both normal and diseased states, we quantified its expression in both post-HCT and control biopsies. P-selectin showed no to low-level expression in a minute fraction of endothelial cells in normal kidneys with significant upregulation in the context of extra-renal acute GVHD (**Fig. 5b**). Regarding galectin-3, multiplexed immunofluorescence was performed to quantify its expression across all nephron compartments. In normal kidneys, galectin-3 exhibited considerable expression in the distal tubules, collecting ducts and collecting tubules, with minimal expression levels observed in the proximal tubules and endothelial cells (**Fig. 5d, f**). In post-HCT patient samples, galectin-3 expression was significantly increased in the distal tubules, collecting tubules, proximal tubules, and peritubular endothelium (AQP1^+^CD31^+^ cells)^58^, indicating a possible role in post-transplant inflammation and tissue remodeling (**Fig. 5c, e, f**). Similar to P-selectin, galectin-3 expression was significantly increased in patients with acute extra-renal GVHD (**Fig. 5g**).

This disease-amplified expression pattern suggested that p-selectin or galectin-3 could serve as spatially enriched anchoring molecules within inflamed renal tissue, enabling preferential accumulation of targeted nanoparticles in the context of injury irrespective of the specific cellular compartment in which they are expressed.

To determine if galectin-3 upregulation extends to other GVHD target organs, we analyzed archival patient skin and gastrointestinal biopsies. Galectin-3 was robustly expressed across GVHD samples (skin, duodenum, and colon), with visibly lower expression observed in the available healthy control tissues (**Supplementary Figs. 12, 13).**

Prompted by the observed elevation of those target proteins, we investigated the potential for targeted delivery of nanomedicines to GVHD kidney tissues. We synthesized lipid nanoparticles (LNPs) comprised of a small molecule drug core and a monolayer of structural phospholipids using microfluidics (**Fig. 6a, b**) (see Methods).

**Fig. 6.**
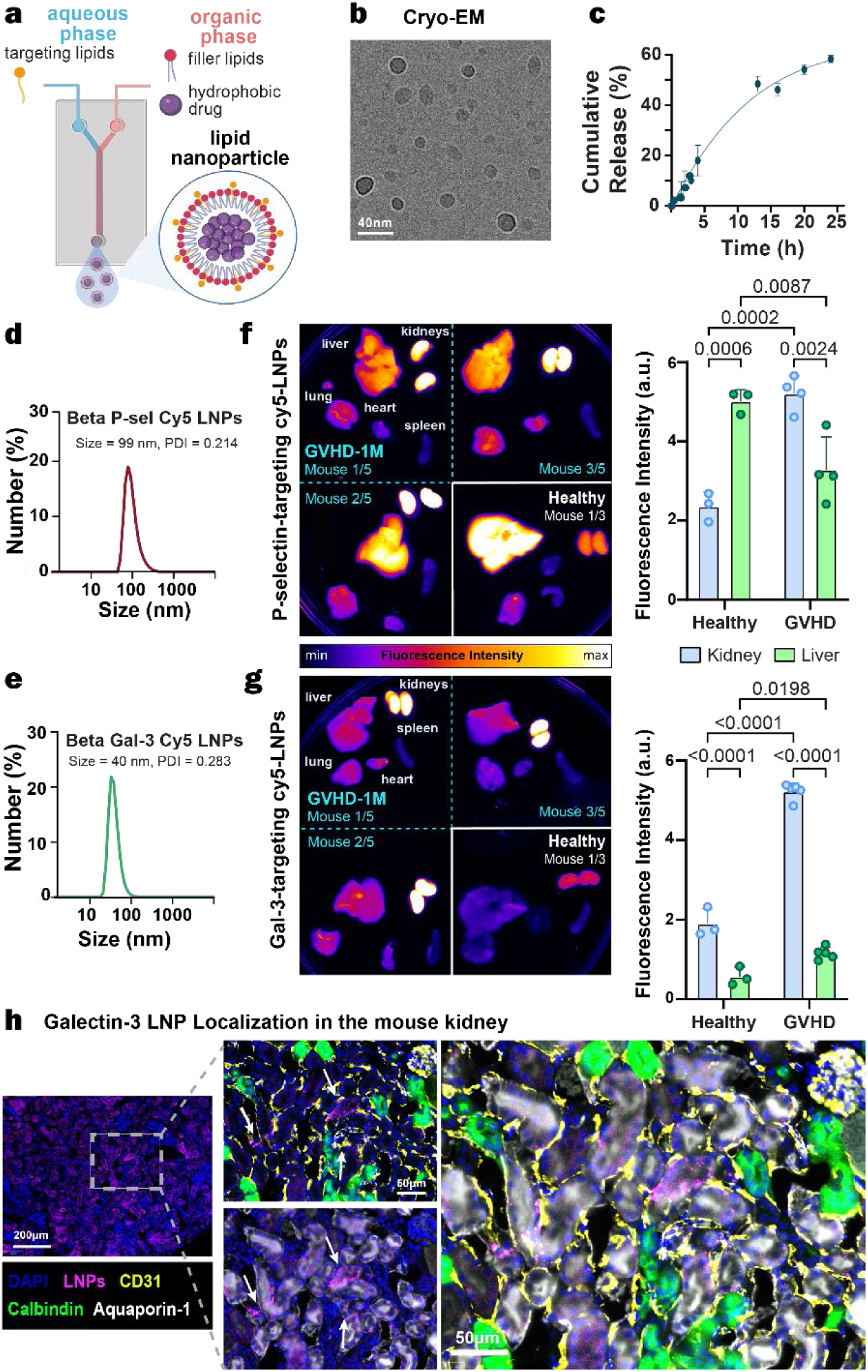
Biodistribution of P-selectin and galectin-3-targeted nanoparticles in GVHD mice. **a,** Schematic of nanoparticle synthesis pipeline **b,** Cryo-EM of Gal-3 targeting betamethasone nanoparticles **c**, Drug release in 50% serum, mean ± s.d **d, e**, Hydrodynamic diameter of Gal-3- and P-sel-targeted LNPs, respectively. **f,** Representative fluorescence image of P-sel-Cy5-LNP biodistribution in the organs of GVHD and healthy mice (left, out of n_H_=3, n_GVHD_=4 per group), and quantification of fluorescence intensity normalized per tissue area in kidney and liver (right). **g,** Representative fluorescence image of Gal-3-Cy5-LNP biodistribution in the organs of GVHD and healthy mice (left, out of n_H_=3, n_GVHD_=5 per group), and quantification of fluorescence intensity normalized per tissue area in kidney and liver (right). **h,** Merged fluorescence image of kidney tissue in mice injected with Gal-3-Cy5-LNPs, showing localization in renal compartments **f,g**: Two way ANOVA with Šidák correction. Data are presented as mean ± s.d.

To achieve preferential accumulation of LNPs at sites of overexpression of P-selectin we incorporated a sulfated glycolipid into the lipid mix (P-sel LNPs). This was substituted with a sphingolipid, to generate LNPs that bind to galectin-3 (Gal-3 LNPs). The hydrodynamic diameter of the particles was drug dependent and ranged from 40-90nm (**Fig. 6b and Supplementary Fig. 14**), while the zeta potential was -40±5mV for all drugs tested. The average drug loading of the particle was 10-13 wt% (weight percentage) with an average release rate of 60% within 24h (**Fig. 6c**). For biodistribution studies, we used Gal-3 LNPs or P-sel LNPs decorated with a cy5-lipid (see Methods) and encapsulating betamethasone dipropionate as the cargo (Beta LNPs) (**Fig. 6d, e).**

We assessed the biodistribution of the targeted LNPs in the context of GVHD-AKI. On day 6 after experimental BMT, Gal-3-Cy5-LNPs or P-sel-Cy5-LNPs were injected by tail IV injection in healthy or GVHD mice and euthanized after 24 hours. Organs were harvested, and fluorescence was measured with a gel reader. Both P-sel LNPs and Gal-3 LNPs accumulated in the kidneys of GVHD mice with higher selectivity than in kidneys of healthy mice. (**Fig. 6f, g).**

Gal-3 LNPs, however, showed a preferential accumulation in kidneys of healthy mice, as compared to the liver (a common site of nanoparticle accumulation). We surmise that these differences may be due to the substantial levels of galectin-3 in kidneys of healthy mice (**Supplementary Fig. 7**) and the smaller hydrodynamic diameter of the Gal-3 LNPs (**Fig. 6e**).

Confocal microscopy of the GVHD kidneys combined with multiplexed immunofluorescence was performed to evaluate LNP distribution across the main nephron compartments, such as proximal tubules (Aquaporin-1^+^), distal tubules (Calbindin^+^), glomeruli (CD31^+^), and endothelial vessels (CD31^+^). Particles were found to localize primarily in the proximal tubules and peritubular capillaries (**Fig. 6h**).

Encouraged by the biodistribution results, we next evaluated whether targeting related pathways with nanoparticle-based therapies could mitigate kidney injury in a preclinical mouse model of GVHD-AKI. We investigated the *in vivo* therapeutic response of nanoparticles designed to bind to the identified delivery targets. Based on the strength of pathway upregulation and the availability of clinically relevant inhibitors, we selected JAK-STAT1 (IFN-γ), JAK-STAT3, and NF-κB as top candidates for pharmacologic inhibition (**Fig. 3b**). However, these therapeutic targets are known to elicit on-target systemic toxicities. Therefore, we synthesized a small library of LNPs, targeted to either galectin-3 or P-selectin, to deliver ruxolitinib (JAK/STAT1 inhibitor), WP1066 (STAT3 inhibitor), Tofacitinib (pan-JAK inhibitor) or TPCA-1 (NF-κB inhibitor) to preliminarily assess their therapeutic efficacy. Ruxolitinib is FDA-approved for treatment of steroid-refractory aGVHD, Tofacitinib is FDA-approved for treatment of arthritis and colitis, while WP1066 and TPCA-1 are still in preclinical development. Mice received daily or every other day IP injections of either the free drug or the drug encapsulated in LNPs, starting from day 2 after transplantation. We tested ruxolitinib (20mg/kg) or WP1066 (20mg/kg) in Gal-3 LNPs, or Tofacitinib (20mg/kg) or TPCA-1 (20mg/kg) in P-sel LNPs (**Fig. 7a)**. The above combinations of encapsulated drugs with targeted lipid carriers were selected based on the stabilities of the drug-loaded LNPs. The stability of all formulations was evaluated over one week in PBS or serum at 4 °C and 37 °C, with Ruxolitinib-Gal-3 LNPs and WP1066 LNPs exhibiting the highest stability under all conditions (**Supplementary Fig. 15**).

**Fig. 7.**
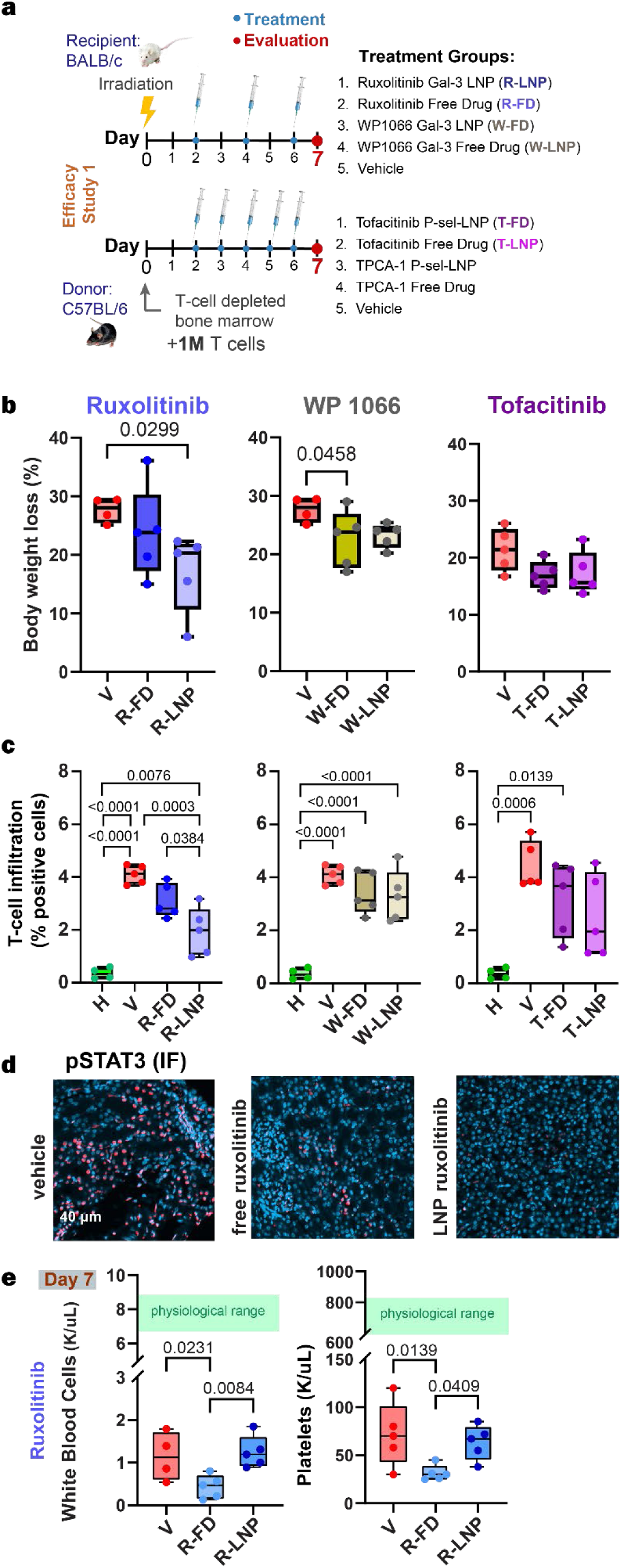
Pharmacological inhibition of JAK/STAT and NF-κB pathways via LNPs on day 7 after BMT. **a,** Schematic of model establishment and treatment regime for the first efficacy study. Gal: Galectin-3 targeting, P-sel: P-selectin targeting. **b**, **c,** Weight loss (**b**) and T cell infiltration (**c**) is reduced more efficiently with LNP-encapsulated Ruxolitinib (left) than WP1066 and Tofacitinib (right). **d**, p-STAT3 (red) levels in mice treated with free or LNP-encapsulated Ruxolitinib. **e**, White blood cell count, and platelet count on day 7 post BMT, show that LNPs can mitigate Ruxolitinib’ s hematotoxicity. R-FD: Ruxolitinib Free Drug, R-LNP: Ruxolitinib LNP, W-FD: WP1066 Free Drug, W-LNP: WP1066 LNP. T-FD: Tofacitinib Free Drug, T-LNP: Tofacitinib LNP. n_H_=3-4, n_GVHD_ = 5 mice per group**. b,d**: Ordinary one-way ANOVA, Tukey correction. The box extends from the 25th to 75th percentiles and the whiskers from min to max. Line represents the median.

Ruxolitinib was the most efficacious in reducing renal T cell infiltration and pSTAT3 out of all the drugs tested, and Gal-3 LNPs improved treatment over the free drug (**Fig. 7** and **Supplementary Fig. 16**). Gal-3 LNP-ruxolitinib also attenuated weight loss compared to other treatment groups (**Fig. 7b**), reflecting reduced overall GVHD burden. Interestingly, we observed that pSTAT3 overexpression was not limited to CD3^+^ cells, but it was also expressed in other cell populations in kidney epithelium, similar to what has been observed with IFNγ in intestinal epithelium of mice with GVHD^72^. Both JAK and STAT3 inhibitors (ruxolitinib, tofacitinib and WP1066) were efficacious in reducing kidney damage biomarkers (NGAL and KIM-1) which were quantified in serum and urine as well as by immunohistochemistry of kidney tissues (**Supplementary Fig. 17**). Representative CD3 and pSTAT3 staining across treatment groups are shown in **Supplementary Fig. 18** and **Supplementary Fig. 19**. PAS staining further confirmed reduced tubular injury, including decreased vacuolization and intratubular cellular debris following treatment (**Supplementary Fig. 20**). The levels of top overexpressed proteins, measured by Western blot (**Supplementary Fig. 21**), were also reduced by inhibitor administration.

Notably, on day 7 post-BMT, mice treated with ruxolitinib-LNPs exhibited a significantly higher recovery in both white blood cell count and platelet count compared to the free drug groups (**Fig. 7d**). This result indicates that the nanoparticles mitigated the known hematologic toxicity of ruxolitinib, which is associated with thrombocytopenia and anemia in patients treated with this drug^34, 36, 37^. These data indicate that spatially restricted JAK inhibition can preserve therapeutic efficacy while reducing systemic exposure of hematopoietic compartments, thereby widening the therapeutic index of ruxolitinib in the context of GVHD.

To evaluate longer-term tolerability, we conducted a 30-day toxicity study in healthy mice receiving intraperitoneal dosing every other day. Serum chemistry and hematological parameters revealed no significant differences between ruxolitinib-loaded galectin-3–targeted lipid nanoparticles, free ruxolitinib, and untreated controls indicating an absence of overt systemic toxicity at the therapeutic dose (**Supplementary Fig. 22**).

Regardless of the delivery approach, administration of the NFκB inhibitor was associated with substantial toxicity in this model. The NFκB inhibitor (TPCA-1) resulted in 100% mortality in all drug-receiving groups (both free drug and encapsulated form) after only two doses in GVHD mice, while it did not cause any weight loss or death in healthy mice at the same dose (20 mg/kg, daily) (**Supplementary Fig. 23**). This result points to on-target toxicity of the drug in GVHD. This deleterious effect is consistent with reports from the use of bortezomib (proteasome and NFκB inhibitor) in GVHD, which paradoxically amplifies GVHD mortality when administered late, due to amplification of inflammatory cytokine generation^77^, but it can have a beneficial effect when administered very early after BMT^78^.

This narrow therapeutic window in NFκB inhibition prompted us to exclude inhibiting this pathway in later studies. We found, thus far, substantial reduction in pSTAT3 expression and T cell infiltration via LNP-encapsulated JAK/STAT inhibitors, especially galectin-3-targeted LNPs encapsulating ruxolitinib which exhibited the greatest reduction in renal inflammation and lower hematotoxicity than the free drug.

### Ruxolitinib LNPs restore kidney impairment in severe GVHD

We next investigated LNP administration in the context of severe GVHD by increasing the dose of donor T cells to 2 million (hereafter denoted as GVHD (2M)) and used this lethal GVHD model to investigate the systemic and kidney-specific efficacy of ruxolitinib-LNPs, since they displayed the highest stability and efficacy out of all candidates. This study also allowed us to introduce more metrics such as glomerular filtration rate and general GVHD scoring. We modified the GVHD-AKI model by injecting 2 million T cells rather than 1M, and we monitored the mice for 14 days instead of 7 days in order to assess the progression of GVHD and kidney injury over an extended period to capture both acute and late-stage effects of ruxolitinib-LNP treatment (**Fig. 8a**).

**Fig. 8.**
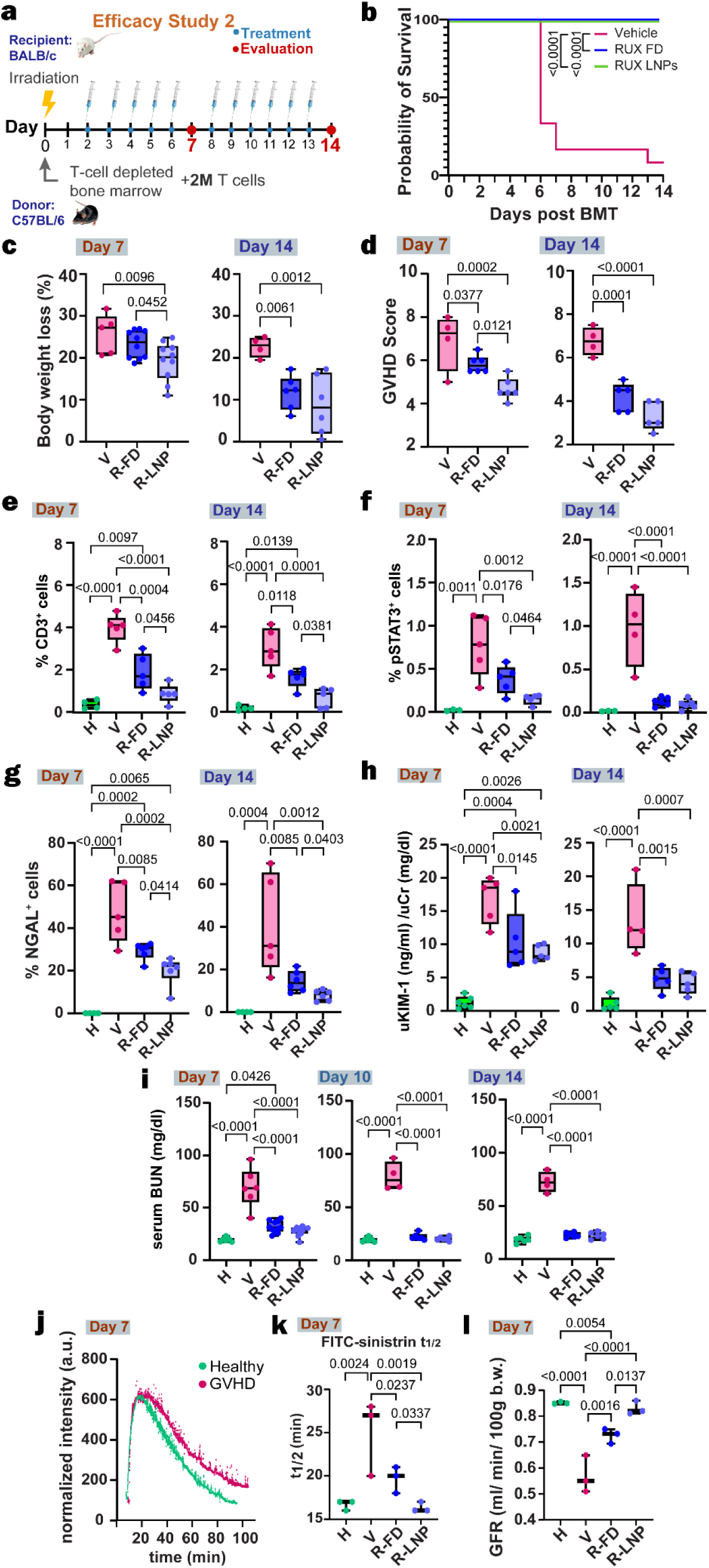
Galectin-3-targeted JAK/STAT inhibition exhibits superior performance to free drug in attenuating severe GVHD (GVHD (2M) and restoring kidney function. **a,** Schematic of severe GVHD (BMT+2M T cells) establishment and treatment regime for the second efficacy study. For days 7 and 14 post-BMT. **b**, Kaplan-Meier survival curve of different groups (comparison with Mantel-Cox log rank test for survival) **c**, Body weight loss per group, mice per group: n_Day7_=5-10, n_Day14_=4-6 **d**, GVHD clinical scoring, mice per group: n_V_ =4, other groups: n=5-6 **e**, T cell infiltration **f**, pStat3 levels (IF quantification) **g**, NGAL levels (IHC quantification). **h**, urine KIM-1 **(e-h)** Mice per group: n_H_=3-6, n_V_=5-6, n_RFD_= 5-6, n_RLNP_ =5-6 **I**, serum BUN levels on days 7, 10 and 14 post-BMT. Mice per group: n_H_=3-6, n_V_=5-6, n_RFD_= 5-11, n_RLNP_ =5-11 **j,** FITC-sinistrin normalized intensity curves for a healthy and a GVHD mouse measured with the Medibeacon transdermal monitor on day 7 post-BMT **k**, Half-time of FITC-sinistrin in the blood on day 7 post-BMT. **l**, Glomerular filtration rate on day 7. **c-I, k-l:** One-way ordinary ANOVA with Tukey correction. **c-i:** The box extends from the 25th to 75th percentiles and the whiskers from min to max. Line represents the median. H: healthy, V: vehicle, R-FD: Ruxolitinib Free Drug, R-LNP: Ruxolitinib LNP.

Mice received daily IP injections of vehicle (DMSO/ Tween 80), free ruxolitinib, or LNP-ruxolitinib and were euthanized 7 or 14 days after transplantation (**Fig. 8a**). Blood was drawn at 7, 10 and 14 days and urine was collected at 7 and 14 days (day 10 was omitted because mice were severely dehydrated). As expected, increasing the amount of T cells during transplantation dramatically increased the severity of GVHD: 65% of mice receiving the vehicle died before day 7 and 90% before day 14. (**Fig. 8b**). Strikingly, administration of LNP-encapsulated ruxolitinib reduced weight loss and improved GVHD clinical scoring (blinded evaluation) to a greater degree than the free drug (**Fig. 8c, d**), indicating improved efficacy against GVHD overall. This observation prompted us to investigate potential concurrent therapeutic effects in liver and gut, traditional targets of GVHD. In mouse livers, while both ruxolitinib-treated groups showed reduced T cell infiltration and apoptosis, there was no significant difference between free drug and LNP groups (**Supplementary Fig. 24**). In the small intestine crypt base region-the primary intestinal site targeted by T cells following transplantation^79^- the LNP formulation showed slightly improved efficacy over the free drug, with greater reductions in pSTAT3 expression, decreased Ki67 expression, and decreased apoptosis and T cell infiltration (**Supplementary Fig. 25**). This differential response between organs may be attributed to the higher levels of galectin-3 overexpression observed in GVHD-affected intestinal tissue compared to liver tissue (**Supplementary Fig. 11**).

Consistent with our first study, LNPs reduced T cell infiltration and STAT3 phosphorylation more than the free drug (**Fig. 8e, Supplementary Fig. 26)**. Kidney injury was assessed by quantifying NGAL by immunohistochemistry and urine KIM-1 by ELISA. NGAL expression significantly increased until day 14 and was reduced more by LNPs as compared to free drug, with the superior performance of LNPs most evident at earlier timepoints (**Fig. 8g**). Urine KIM-1 was elevated in GVHD mice, but the difference between the performance of LNPs and free drug was not significant (**Fig. 8h**), likely due to the small number of samples, as urine collection from these mice was not always possible. The severity of renal injury was reflected in the abnormal elevation of serum BUN, which is not a very sensitive biomarker in mice^80^, and was significantly higher in GVHD mice (**Fig. 8i**). Treatment with ruxolitinib maintained serum BUN close to the normal range (**Fig. 8i**). Importantly, glomerular filtration rate was significantly reduced in GVHD mice and was improved by LNP-ruxolitinib administration to a greater extent than the free drug (**Fig. 8j-k**). Transcutaneous glomerular filtration rate (tGFR) measurements, normalized for animal weight, were performed using a transdermal continuous kidney function monitor (see Methods). Overall, these results demonstrate that severe GVHD has a significant impact in kidney function that can be restored more rapidly and efficiently by the administration of an LNP-encapsulated ruxolitinib as compared to the free drug.

## Discussion

Kidney injury is a common, potentially life-threatening, and inadequately understood complication of BMT, with reported incidence rates ranging up to 70% following clinical transplantation^27, 29^. Because kidney biopsies in those patients are rarely performed due to transplant-associated thrombocytopenia and coagulopathy^81, 82^ the histopathologic basis of this injury remains poorly characterized. Notably, even in the largest heretofore published biopsy and autopsy series to date, no definitive morphological correlates of acute GVHD in the kidney were identified using conventional histopathologic assessment^82^. Renal dysfunction in these patients has been largely attributed to drug nephrotoxicity and thrombotic microangiopathy^22^.

By applying multiplexed immunofluorescence with machine learning-assisted tissue classification to analyze post-HCT kidney biopsies, we identified a donor-derived, Th1-polarized cytotoxic infiltrate that correlated with extra-renal GVHD severity. This finding reframes kidney injury in these patients, suggesting an important component of allo-immunity for the kidney damage.

Our analysis of 20 BKV-negative post-HCT patient biopsies, collected within the first 14 months post-transplant, revealed significant renal T cell infiltration particularly within the epithelial compartment. FISH analysis confirmed that kidney-infiltrating T cells were donor-derived, establishing the allograft as the source of the immune attack on the kidney.

Phenotypically, infiltrating T cells in acute GVHD kidneys were CD8-dominant, activated (CD69^+^), and polarized toward a cytotoxic Tc1 phenotype, with significant enrichment of T-bet and Granzyme B and virtual absence of the Th2 master regulator GATA3. This profile is consistent with the effector phenotype described in other acute GVHD target organs such as the gut, where alloreactive donor T cells differentiate along a Th1/Tc1 axis characterized by IFN-γ production, T-bet expression, and cytotoxic granule-mediated killing of epithelial cells^3, 67, 72, 79.^

Our parallel observations here in the kidney - donor-derived, CD8^+^T-bet^+^Granzyme B^+^ effector cells infiltrating the tubular epithelium - provides compelling evidence that the kidney may be subject to a similar alloimmune program that drives injury in canonical GVHD target organs such as the GI tract^57^. Additionally, the degree of T cell infiltration in post-HCT kidneys was significantly greater than in kidneys from patients with kidney injury due to other etiologies, indicating that the observed infiltrate is not merely a nonspecific consequence of tissue injury but reflects a disease-specific alloimmune process.

The presence of T cells in the proximal and distal tubular renal epithelium, coupled with acute tubular injury and interstitial inflammation, suggests that in patients with acute extra-renal GVHD, kidney injury is mediated at least in part by immune mechanisms, even though the kidneys have not been considered traditional GVHD target organs^3^. While kidney injury post-HCT is multifactorial and drug nephrotoxicity can contribute to it, most patients in this study were not receiving nephrotoxic medications at the time of biopsy, emphasizing that the immune-mediated component of kidney injury should not be overlooked.

These human findings were recapitulated in our preclinical models. In the MHC-mismatch model (C57BL/6→BALB/c), GVHD kidneys exhibited a marked expansion of T-bet^+^CD8^+^ effector cells with scant FoxP3+ regulatory T cells, indicating a strong skew toward a pro-inflammatory effector response with limited regulatory counterbalance. Severe GVHD induced significant impairment in kidney function, highlighted by a significant reduction in glomerular filtration rate and increase in BUN, alongside apoptosis of tubular and glomerular cells. Notably, an independent minor mismatch model (129S1→C57BL/6) produced renal injury of equivalent severity, with comparable T-cell infiltration, tubular injury, and pSTAT3 activation, indicating that GVHD-mediated kidney damage is not attenuated in the more clinically relevant mismatch setting. The concordance between human and murine immune profiles - across two independent preclinical models and patient biopsies - substantially strengthens the case that kidney injury in post-HCT patients with acute GVHD represents *bona fide* immune-mediated organ damage driven by the same Th1/Tc1 effector program that characterizes GVHD in the gut and liver. This alignment with human kidney pathology supports our model as a method for investigating GVHD-associated kidney injury.

Our transcriptome analysis in the kidneys of GVHD mice revealed extensive upregulation of genes related to inflammation and allograft responses, as well as genes related to kidney injury and fibrosis. Among overexpressed pathways in diseased kidneys, we identified JAK-STAT and NF-κB as potential pharmacologic targets for existing small molecule drugs to abrogate kidney injury. JAK/STAT signalling is a key mediator of GVHD pathogenesis and forms the basis for the clinical use of ruxolitinib^37, 71^. However, we anticipated that the systemic administration of small molecule inhibitors of these pathways would not yield a favorable pharmacokinetic profile in the kidneys^43, 44^ and could result in undesirable on-target side effects in other tissues. We identified two potential drug delivery targets overexpressed in the GVHD model: galectin-3 and endothelial P-selectin, both known mediators of inflammation. In post-HCT human biopsies, endothelial P-selectin expression was increased compared to controls, particularly in patients with acute GVHD. Similarly, galectin-3 expression increased significantly in most kidney compartments post-transplant, including proximal tubule and peritubular endothelium, further supporting its role in the inflammatory response associated with GVHD-induced kidney injury. Notably, galectin-3 upregulation was not restricted to the kidney; robust expression was also detected in skin, duodenum, and colon biopsies from GVHD patients, suggesting that galectin-3 represents a damage-associated lectin expressed broadly across GVHD target organs.

Mechanistically, renal epithelial galectin-3 upregulation is driven by stress-activated transcription factors and oxidative stress signalling engaged in parallel with, rather than downstream of, the canonical IFN-γ/JAK/STAT cytotoxic axis. Infiltrating alloreactive Th1 cells release high levels of TNF-α and generate local oxidative stress, which, combined with acute cellular injury, strongly activates NF-κB and related stress-responsive transcription factors. The LGALS3 promoter is highly responsive to these parallel stress-activated pathways, leading to robust galectin-3 expression. From a therapeutic design perspective, this divergent transcriptional regulation is highly advantageous: galectin-3 remains available as a targeting handle even when downstream JAK/STAT signalling is pharmacologically blocked. While previous works have primarily identified galectin-3 as a biomarker or pathogenic mediator in ischemic and toxic acute renal failure^83, 84, 85, 86^, as well as a contributor to progressive renal fibrosis^76, 87, 88^, our approach uniquely repositions this damage-associated lectin as a disease-induced molecular address for precision drug delivery. Because galectin-3 is driven by parallel injury pathways rather than being strictly dependent on JAK/STAT signaling, it serves as a highly stable, orthogonal binding address on the injured renal epithelium. This decoupled regulation ensures that as our targeted lipid nanoparticles deliver therapeutic payloads to silence the destructive JAK/STAT pathway, the local therapeutic effect does not prematurely downregulate or erase the galectin-3 targeting moiety itself, allowing for sustained, localized delivery.

We incorporated certain glycolipids that bind to galectin-3 and P-selectin into phospholipid-based nanoparticles encapsulating small molecule drugs, to enhance their specificity to the affected tissues. The resulting nanoparticles exhibited very small sizes (as low as 15 nm for Gal-3-targeted ruxolitinib-LNPs), high drug encapsulation efficiency, and preferential accumulation in diseased mouse kidneys as compared to other organs and healthy kidneys.

To assess the therapeutic potential of the upregulated targets for both targeted inhibition and drug delivery, we designed nanoparticles to bind to either galectin-3 or P-selectin and loaded them with several different small molecule inhibitors. JAK/STAT inhibitors were most successful in reducing the severity of kidney injury. In a moderate MHC-disparate GVHD mouse model, ruxolitinib encapsulated in galectin-3-targeted LNPs outperformed other inhibitors and free ruxolitinib in reducing inflammation and minimizing drug hematotoxicity.

Ruxolitinib delivery utilized Gal-3-targeted LNPs, which demonstrated greater renal accumulation and superior colloidal stability as compared to the P-selectin-targeted LNPs required for tofacitinib formulation (see more in Supplementary Discussion Note 1). Ruxolitinib directly inhibits the JAK1/JAK2 signalling pathway of multiple cytokines, including the IFN-γ/STAT1 axis, which appeared to be active in our analyses of renal T cell infiltration in GVHD, while WP1066 targets the downstream STAT3 node, a less proximal signal in this effector program. Ultimately, targeted delivery maximizes efficacy when a stable, well-matched platform is paired with a drug addressing the most proximal pathogenic driver, and Gal-3 LNP delivery of ruxolitinib achieves this through both superior renal accumulation and on-target pathway specificity.

This improvement addresses a well-recognized dose-limiting adverse effect of systemic ruxolitinib in GVHD patients - thrombocytopenia and anemia^34, 35, 37^ - and reflects a fundamental advantage of the nanoparticle approach: effective JAK/STAT pathway inhibition in diseased tissue while mitigating systemic exposure of hematopoietic compartments. A 30-day tolerability study in healthy mice further confirmed that galectin-3-targeted ruxolitinib LNPs did not induce overt systemic toxicity at the therapeutic dose, supporting the favorable safety profile of this delivery approach. The increased efficiency led us to further test LNPs in severe, lethal GVHD, over a two-week period. Gal-3-targeted LNPs continued to prove more effective than free ruxolitinib, not only in alleviating kidney injury and restoring GFR but also in reducing overall GVHD scoring and reducing pathology in the liver and intestines. This concurrent multi-organ benefit is consistent with our observation that galectin-3 is upregulated across GVHD target tissues, providing a mechanistic rationale for a platform that could address systemic GVHD through a single targeting moiety.

## Outlook

The clinical relevance of this work is underscored by the convergent findings from both human biopsies and mouse models highlight the role of immune-mediated injury in GVHD-associated kidney damage, which is often overlooked. Both human biopsies and murine data showed increased P-selectin and galectin-3 expression in acute GVHD, supporting their considerations as targets for nanoparticle-based drug delivery. The implications of our work are significant for clinical practice, as GVHD remains a major challenge in allo-HCT. Currently, ruxolitinib is the only approved drug for steroid refractory acute GVHD but its use in the clinic remains limited due to toxicities such as thrombocytopenia and anemia^34, 36, 37^. It is important to note that, in GVHD alone, pancytopenia and, in particular, neutropenia are linked to a higher risk for opportunistic infections that can be fatal^77^. Our data suggest that the systemic hematotoxicity of ruxolitinib may be mitigated using galectin-3-targeted nanoparticle delivery, allowing for the continuation of ruxolitinib treatment beyond what is currently feasible. These nanoparticles can be loaded by many classes of small molecule inhibitors, thereby expanding the applicability of this platform beyond the drugs evaluated in this work. Additionally, the phospholipids used in our nanoparticles are endogenous components of organisms, have a demonstrated excellent safety profile and have been used in FDA-approved formulations^89, 90^.

More broadly, this study illustrates how direct analysis of human disease tissue can guide the rational selection of nanoparticle targets, complementing the formulation- and chemistry-driven strategies that have dominated nanomedicine development. By anchoring target discovery in the pathological features of the diseased organ itself -rather than in pre-existing receptor or ligand libraries- this disease-guided approach may be particularly well-suited to conditions like GVHD, where the affected tissues display distinctive, disease-amplified surface molecules that would be difficult to predict *a priori*. Extending this logic to other inflammatory and alloimmune disorders, where human biopsy material is available but underutilized for delivery design, represents a promising direction for future work.

Regarding limitations of this study, the human cohort, while representing one of the largest collections of renal biopsies within 14 months post-HCT reported to date, remains retrospective and derived from a single center, and larger multi-institutional studies are warranted to establish the prevalence and heterogeneity of GVHD-associated renal pathology. Additionally, while T-cell infiltration was significantly enriched in GVHD patient kidneys, the histologic injury features observed in biopsies (tubular injury, glomerulosclerosis, vascular changes) are multifactorial in origin, limiting our ability to attribute specific structural changes to alloimmune injury alone. Additionally, the absence of an established murine model of chronic GVHD with sustained, well-characterized renal involvement beyond approximately 30 days constrains long-term efficacy assessment; extending GVHD models past this window introduces confounding mortality driven by systemic disease severity rather than renal pathology. Additionally, all efficacy studies here employed an early treatment regimen beginning on day 2 after transplantation. While this design reflects clinically relevant prophylactic strategies in high-risk patients, evaluating galectin-3-targeted LNPs in a delayed, therapeutic dosing regimen would broaden translational applicability.

In conclusion, our findings demonstrate that targeted nanotherapeutics, guided by both early human data and robust preclinical models, can effectively mitigate GVHD-associated kidney injury. The use of galectin-3-targeted nanoparticles provides a promising approach to localizing treatment in inflamed renal compartments, reducing systemic side effects, and improving outcomes in post-transplant patients.

## Methods

### Patient and Control Sample Collection

This study was conducted in accordance with the principles of the Declaration of Helsinki and was approved by the Institutional Review Board (IRB) of Memorial Sloan Kettering Cancer Center (IRB#16-834). Kidney biopsies were obtained from 29 bone marrow transplant (BMT) recipients who underwent biopsy within the first 14 months post-allogeneic hematopoietic cell transplantation (allo-HCT) at MSKCC between 2010 and 2024. Written informed consent was obtained from all patients or their legal guardians prior to tissue collection. Patients with confirmed BK virus nephropathy were excluded from this study to prevent confounding effects on inflammation analysis.

Control kidney biopsies were obtained from formalin-fixed, paraffin-embedded (FFPE) sections of nephrectomy specimens from patients with renal tumors. Only histologically normal kidney tissue, uninvolved by neoplastic or inflammatory processes, was used as controls. The sections were reviewed by a renal pathologist to confirm the absence of pathology. All patient data were de-identified and analyzed in compliance with institutional guidelines and HIPAA regulations.

### GVHD Stratification

Extra-renal GVHD status was annotated based on comprehensive clinical records and consensus assessment by the multidisciplinary bone marrow transplantation (BMT) team at MSK, incorporating physician-documented diagnoses, longitudinal clinical course, and treatment history, both at the time of biopsy and across the post-transplant period (“ever GVHD”).

### Mice bone marrow transplantation and assessment of GVHD

All mice in this study were maintained under protocols approved by the Institutional Animal Care and Use Committee at Weill Cornell Medicine and MSKCC. Eight- to ten-week-old female C57BL/6 and BALB/c recipient mice and 129S1 donor mice were purchased from Jackson Laboratories. In the major mismatch model, mouse allo-HSCT experiments were performed as previously described, with BALB/c mice receiving 900 cGy of fractionated lethal radiation followed by tail vein injection of 10×10^6^ T cell depleted (with anti–Thy-1.2 and Low-Tox-M rabbit complement (CEDARLANE Laboratories) bone marrow from C57BL/6 mice. In the minor mismatch model, C57BL/6 mice received 1100 cGy split dose radiation followed by tail vein injection of 15×10^6^ T cell depleted (with anti–Thy-1.2 and Low-Tox-M rabbit complement (CEDARLANE Laboratories) whole splenocytes from 129S1 mice. Bone marrow only (BMO group) received only bone marrow, while the GVHD group received intravenously splenic T cells. Donor T cells were prepared by harvesting donor splenocytes and enriching T cells by Miltenyi MACS purification of CD3 (routinely >90% purity). Before the beginning of each experiment, we separated each cage of mice from the vendor to be divided as equally as possible among the various treatment groups. This helped to minimize the chances of incorrectly attributing differences observed in renal pathology to treatment effects, when in reality they are due to cage effects. Animals were housed under a 12/12 h light/dark cycle and given access to food and water ad libitum. All groups are monitored up to two weeks after the BM transplant for GVHD and scored according to five clinical parameters (weight, posture, fur, skin, activity).

The numbers of mice analyzed are provided in the main text and/or in the figure legends.

### Staining and Image Analysis

#### Tissue Selection and Processing

- Human Tissues:

`o` **Post-HCT patients:** Formalin fixed, paraffin embedded (FFPE) kidney needle-core biopsies were used for unstained sections.
`o` **Controls:** Unstained FFPE-derived sections were cut from grossly and microscopically normal kidney, routinely sampled in nephrectomy specimen derived from patients with renal tumors. The sections were confirmed to be uninvolved by any neoplastic or overt inflammatory process after review of the respective H&E slides by a pathologist.
- **Murine Tissues:** Tissues were fixed in 4% paraformaldehyde overnight and then transferred to 70% ethanol. Fixed tissues were embedded in paraffin and sections prepared at a thickness of 5 μm.

#### Multiplex Immunofluorescence (human & mouse)

Multiplex immunofluorescence followed by sequential immunohistochemistry (IHC) with AEC chromogen was performed on Leica Bond RX staining processors with paraffin tissue sections in a method recently developed by the MSKCC Pathology and Microscopy Core^91^.

Samples for immunofluorescence were pretreated with EDTA-based epitope retrieval solution for 20 minutes at 95°C. Primary antibodies and their working concentrations or dilutions for mouse tissue are listed in **Supplementary Table 7**. Those for human tissue are listed in **Supplementary Table 8**. Leica Bond HRP Polymer was applied followed by CF® dye tyramide conjugate 594, 543 or 430 (Biotium, B92174, 92172, 96053) or Alexa Fluor tyramide signal amplification reagents 488 and 647 (Life Technologies, B40953, B40958) for detection. After each round of IF staining, a same epitope retrieval was performed for denaturization of primary and secondary antibodies before another primary antibody was applied. After the run was finished, slides were washed in PBS and incubated in 5 μg/ml 4’,6-diamidino-2-phenylindole (Sigma Aldrich, Cat#D9542) in PBS for 5 min, rinsed in PBS, and mounted in Mowiol 4–88.

Immunohistochemical staining of CD3 (human samples) or CD19 (mouse samples) was performed at Molecular Cytology Core Facility of Memorial Sloan Kettering Cancer Center using Leica Bond RX. Samples were pretreated with EDTA-based epitope retrieval solution for 20 minutes at 95°C. Then the primary antibody against CD3 or CD19 was used. Leica Bond HRP Polymer was applied followed by AEC substrate kit (Vector, Cat# SK-4205). AEC chromogen was prepared according to manufacturer’s instruction. After the run was finished, slides were washed in PBS and incubated in 5 μg/ml 4’,6-diamidino-2-phenylindole (Sigma Aldrich, Cat#D9542) in PBS for 5 min, rinsed in PBS, and mounted in Mowiol 4–88.

Slides were scanned using a Panoramic 250 scanner (3DHistech, Budapest, Hungary). A 20× 0.8NA objective was used. The following filters were used to image each fluorophore:

The multiplexed fluorescence was imaged with a pco.edge 4.2 4MP camera, while the AEC slides were imaged with a CIS VCC-FC60FR19CL camera. Regions of interest (ROI), drawn around the whole kidney, from CaseViewer (https://www.3dhistech.com/ v2.4.0.119028) images were exported into Tiled TIFF files, followed by computational alignment and generation of pseudo-fluorescence images from IHC stains done in MATLAB^91^.

| Fluorophore | Excitation (nm) | Dichroic (nm) | Emission (nm) |
| --- | --- | --- | --- |
| DAPI | 387/18 |  | 440/44 |
| CF 430 | 436/20 | 455 | 480/40 |
| Alexa 488 | 482/18 | 506 | 527/30 |
| CF 543 | 530/11 | 550 | 572/28 |
| CF 594 | 586/15 | 600 | 631/36 |
| Alexa 647 | 650/13 | 668 | 700/50 |

#### Immunohistochemistry for CD3 and NGAL (mouse)

Immunohistochemical staining of murine kidney tissues was performed at the Molecular Cytology Core Facility of Weill Cornell Medicine or MSKCC. Kidney tissues were fixed in 4% paraformaldehyde overnight. Fixed tissues were embedded in paraffin and sections prepared at a thickness of 5 μm. CD3, NGAL: IHC was performed using a rabbit polyclonal CD3 antibody (Agilent, Santa Clara, CA) and a Rabbit monoclonal NGAL antibody [EPR21092] (Abcam, Cambridge, MA) on a Leica Bond system (Buffalo Grove, IL) following the manufacturer’s protocol. The sections were pre-treated using heat mediated antigen retrieval with Tris-EDTA buffer (pH = 9, epitope retrieval solution 2) for 20 min or Sodium Citrate buffer (pH = 6, epitope retrieval solution 1) for 30 min. Tissue sections were incubated with CD3 or NGAL antibody (1:100 dilution) for 15 min at room temperature. Target protein was detected using an HRP conjugated compact polymer system and DAB as the chromogen. Each section was counterstained with hematoxylin and mounted with Leica Micromount.

All images were taken with either a bright-field and fluorescence microscope (Zeiss Axio Observer) or digital Panoramic Slide Scanner (3D Histech, Budapest Hungary). MRXS raw images were post-processed using Fiji or QuPath 0.4.3.

#### Galectin Immunofluorescence (mouse)

On the day of harvesting, mice were anesthetized with ketamine/xylazine and then subjected to transcardial perfusion with PBS until colorless fluid was observed coming from the right atrium. Tissues were fixed in 4% paraformaldehyde overnight and then transferred to 70% ethanol. Fixed tissues were embedded in paraffin and sections prepared at a thickness of 5 μm. Immunofluorescence experiments were performed at Molecular Cytology Core Facility of Memorial Sloan Kettering Cancer Center using Leica Bond RX. Samples were pretreated with EDTA-based epitope retrieval solution for 20 minutes at 95°C. Then the primary antibody against galectin 3 (invitrogen, Cat# 14-5301-82, 1ug/mL) was used. Leica Bond HRP Polymer was applied followed by Alexa Fluor tyramide signal amplification reagents 488 (Life Technologies, B40953) for detection. After the run was finished, slides were washed in PBS and incubated in 5 μg/ml 4’,6-diamidino-2-phenylindole (Sigma Aldrich, Cat#D9542) in PBS for 5 min, rinsed in PBS, and mounted in Mowiol 4–88.

#### P-selectin and CD31 Immunofluorescence (mouse)

On the day of harvesting, mice were anesthetized with ketamine/xylazine and then subjected to transcardial perfusion with PBS until colorless fluid was observed coming from the right atrium. Tissues were fixed in 4% paraformaldehyde overnight and then transferred to 70% ethanol. Fixed tissues were embedded in paraffin and sections prepared at a thickness of 5 μm. Immunofluorescence experiments were performed at Molecular Cytology Core Facility of Memorial Sloan Kettering Cancer Center using Leica Bond RX. Samples were pretreated with EDTA-based epitope retrieval solution for 20 minutes at 95°C. Then the primary antibodies against p-selectin (LSBio, Cat#LS-B3578) and CD31 (Abcam, Cat#ab182981) were used sequentially. Leica Bond Polymer anti-rabbit HRP was applied followed by CF® dye tyramide conjugate 594 (Biotium, B92174) or Alexa Fluor tyramide signal amplification reagents 488 (Life Technologies, B40953) for detection. After each round of IF staining, a same epitope retrieval was performed for denaturization of primary and secondary antibodies before another primary antibody was applied. After the run was finished, slides were washed in PBS and incubated in 5 μg/ml 4’,6-diamidino-2-phenylindole (Sigma Aldrich, Cat#D9542) in PBS for 5 min, rinsed in PBS, and mounted in Mowiol 4–88.

### Whole slide image analysis (human and mouse)

#### Machine-learning assisted whole biopsy classification (human)

Image handling was carried out using the open-source digital pathology software QuPath v0.5.1 (qupath.github.io). To account for intra-patient variability while maintaining a high accuracy for the algorithm, we trained a different pixel classifier for each biopsy, using a combination of fluorescence markers (Calb, Aqp1, Aqp2) and input from a renal pathologist. We then quantified CD3^+^ cells in the epithelium and stroma using Qupath’s cell detection watershed algorithm; we considered those in the epithelium as potential effector T cells, while those in the stroma were identified as non-resident or circulating T cells. To reduce file size and computation time during analysis, all channels corresponding to epithelium (Aqp1, Aqp2, Calbindin) from the multiplex imaging were merged into a single channel. Following this preprocessing step, we employed a machine-learning assisted pixel classifier based on a random forest algorithm in QuPath (version 0.5.1) to classify the kidney tissue sections as stroma, epithelium, or background. A total of 150 manual annotations per slide were generated as ground truth, informed by fluorescence markers and reviewed by a renal pathologist who reviewed the corresponding PAS staining to ensure accurate annotation. These annotations were split into 60% for training, 20% for validation, and 20% for testing. The classifier differentiated between stroma, epithelium, and background, although background regions, which were cell-free, were excluded from CD3 quantification. After optimizing model parameters to achieve an F-score greater than 0.8, the classifier was applied to the entire biopsy image, allowing for robust, machine-learning assisted separation of stromal and epithelial regions for subsequent CD3 quantification.

#### Immunofluorescence image analysis (human or mouse)

Cell segmentation was carried out using the open-source digital pathology software QuPath v0.5.1 (qupath.github.io). Nuclear detection was performed on the DAPI channel using an unsupervised watershed algorithm with parameters tuned on a validation set of 10 ROI. After nuclear detection, the cytoplasm around each nucleus was simulated by cell expansion of 5 μm and measurements generated for marker intensity in different compartments (mean, minimum, maximum and standard deviation of intensity in cytoplasm, nucleus, or the whole cell). Positivity was determined by the intensity of each marker in the primary cell compartment where it is usually expressed. In our study, markers were cytoplasmic or membranous. Guided by our pathology core, a single threshold for each marker was selected as a cut-off to determine positivity across the entire dataset. The threshold was identified by its ability to separate positive from negative cells in a set of 10 ROIs from 10 different samples. In order to avoid bias introduced during ROI selection, each ROI included a whole kidney slice, which resulted in detection of 200.000-400.000 cells per kidney, with the associated intensity values for each channel. Since those files were too large to open and manually navigate, a Python script was used to quantify cells which were positive (or double/triple positive) based on the thresholds set above for each cellular compartment.

#### Immunohistochemistry image analysis (human or mouse)

Nuclear detection was performed on the hematoxylin channel and positivity was determined by the intensity of the DAB chromogen, with the thresholding validation performed as described above.

### Immunoblotting (mouse)

Kidney tissue was dissected from mice and homogenized through tissue sonication and a sonicating tip in tissue lysis buffer (5% of 50 mM Tris buffer pH 7.4, 2% SDS, 10% glycerol and 1% protease and phosphatase inhibitor cocktail (ThermoFisher 78446). Bradford reagent (BioRad 5000205) was mixed 1:1 with deionized water diluted samples containing BSA standard or the sample lysate. Absorbance was measured at 595 nm in a Tecan Infinite M1000 Pro plate reader. Loading buffer consisted of 1x Laemmli Buffer/2-Mercaptoethanol (BioRad 1610747/1610710) and deionized water if necessary. Final loading samples were aliquoted and stored at -80°C until use. Frozen samples were immediately heated for 8 minutes at 75°C (phospho-protein targets) or for 6 minutes at 90°C (all other targets). TGX precast gels (BioRad 4568094, 4-20%) were loaded into a Tetra Cell electrophoresis apparatus (BioRad 1658004) filled with chilled 1x Tris/Glycine/SDS buffer (BioRad 1610732). 1.5 μL running ladder (LI-COR 92698000), 10-30 μL sample, or 10-30 μL blank sample buffer were injected into each lane. Electrophoresis was run at 110 Volts, CV, for 80 minutes or until appropriate ladder separation. Blot sandwiches comprised a 0.2 μm PVDF membrane (BioRad 1620174) pre-incubated in methanol followed by transfer buffer (BioRad 10026938) held between buffer wetted-stack paper (BioRad) and the gel. A Trans-Blot Turbo transfer system (BioRad 1704150) was used on the 7-minute Midi-Turbo setting. Blot membranes were cut and immediately rinsed in 1x TBS, then moved into a blocking solution (3% w/v BSA in 1x TBS-T) (BioRad 1706435; Sigma P1379) and agitated at room temperature for 1.5 hours. Primary antibody was diluted into 1 mL of blocking buffer, which was sandwiched with the membrane between two parafilm sheets before sealing. Membrane sandwiches were incubated overnight at 4°C with rotation. Membranes were peeled from the parafilm into 1x TBS-T and washed three times for 10 minutes at room temperature with agitation. HRP conjugated secondary antibody was diluted in enough blocking solution to cover the membrane for 1 hour at room temperature with agitation. After three more 10-minute washes, membranes were rinsed in RT 1x TBS then incubated for 1 minute in HRP substrate (Millipore WBLUR0500). The membranes are imaged on an Amersham ImageQuant 800.

The antibodies used are described in **Supplementary Table 5**.

### Serum and urine biomarker quantification (mouse)

For serum chemistry, blood was collected into tubes containing a serum separator. The tubes were then centrifuged, and the serum was obtained for analysis. Blood urea nitrogen concentration (BUN) and creatinine concentration were measured in a Beckman Coulter AU680 analyzer. NGAL and KIM-1 were measured using Abcam ELISA kits (Mouse Lipocalin-2 ELISA Kit (NGAL) (ab199083) and Mouse KIM-1 ELISA Kit (TIM1) (ab213477). Urine was centrifuged before ELISA measurements according to the kit’s instructions. Urine NGAL/KIM-1 values were normalized to urine creatinine, also measured in a Beckman Coulter AU680 analyzer.

### Glomerular filtration rate measurements (mouse)

Transcutaneous glomerular filtration rate (tGFR) measurements, normalized for animal weight, were performed using a transdermal continuous renal function monitor (MediBeacon GMBH, Manheim, Germany). The mice involved in the experiment were weighed, anesthetized, and had their fur removed from the upper back region in the thoracic area by shaving and epilation cream. The devices were then secured on the hairless area using double-sided adhesive patches provided by MediBeacon and surgical tape (Micropore, 3M Health Care, St Paul, MN). A mixture of normal saline and FITC-sinistrin (Mannheim Pharma & Diagnostics, Mannheim, Germany) was injected intravenously via tail vein. Devices were left on for at least 65 min while mice were placed in individual cages. Measures were taken to ensure minimal stimulus including lower lighting and minimal sound. GFR was calculated from FITC-sinistrin plasma clearance using an established two-compartment model^92, 93^. MB_Studio3 software and MB_Lab2 software (MediBeacon GMBH, Manheim, Germany) was used for importing raw data from the device and GFR quantification

### RNA sequencing (mouse)

#### RNA extraction

On the day of harvesting, mice were anesthetized with ketamine/xylazine and then subjected to transcardial perfusion with PBS until colorless fluid was observed coming from the right atrium. Kidneys were immediately harvested, and flash frozen in liquid nitrogen. 20-30 mg frozen tissue were homogenized in 1mL TRIzol Reagent (ThermoFisher catalog # 15596018) and phase separation was induced with 200 µL chloroform. RNA was extracted from 350 µL of the aqueous phase using the miRNeasy Mini Kit (Qiagen catalog # 217004) on the QIAcube Connect (Qiagen) according to the manufacturer’s protocol. Samples were eluted in 35 µL RNase-free water.

#### Transcriptome sequencing

After RiboGreen quantification and quality control by Agilent BioAnalyzer, 500 ng of total RNA with RIN values of 8.9-9.8 underwent polyA selection and TruSeq library preparation according to instructions provided by Illumina (TruSeq Stranded mRNA LT Kit, catalog # RS-122-2102), with 8 cycles of PCR. Samples were barcoded and run on a NovaSeq 6000 in a PE100 run, using the NovaSeq 6000 S4 Reagent Kit (200 Cycles) (Illumina). An average of 37 million paired reads was generated per sample. Ribosomal reads represented 0.62% of the total reads generated and the percent of mRNA bases averaged 87%.

#### Transcriptome sequencing analysis

Quality control (QC) was performed with FastQC (v0.11.9)^94^. Raw reads were trimmed using Trimmomatic (v0.38)^95^ with default parameters for paired-end reads and the cropping option specific to the TruSeq PE adapters (TruSeq3-PE-2.fa). Trimmed reads were then aligned with STAR aligner (v2.7.0e)^96^ against the mouse genome assembly (GRCm38.p5) and the aligned reads were in turn assigned to genes using featureCounts (v1.6.3)^97^. The resulting count tables were imported to R for further processing, analysis, and visualization. Lowly expressed genes were filtered out using edgeR’s filterByExpr function and normalization factors for library size scaling were calculated using edgeR’s calcNormFactors function. Differential expression analysis was performed using the limma’s voom function and a threshold of FDR≤0.05 was set to define genes that change with significance between the different datasets. The table of all differentially expressed genes and their fold changes was used as a pre-ranked list in GSEA (v4.2.2)^68^ against the mouse gene set hallmark resource (mh.all.v0.3.symbols.gmt) to predict signaling pathways that are enriched in any of the pairwise comparisons. Pathways were defined as enriched if they had a false discovery rate (FDR) value < 0.05.

### Reverse transcription and RT-qPCR

cDNA was generated using the Superscript™ IV VILO™ Master Mix with ezDNase™ Enzyme according to the manufacturer’s protocol (Applied Biosystems). RT-qPCR was performed using Applied Biosystems TaqMan Fast Advanced Master Mix on a ViiA7 PCR machine (Life Technologies). PPIA and RPS18 were chosen as endogenous controls as they are the most stable in inflammatory disease models^98, 99^. Fold changes in expression were calculated using the ΔΔCT method. We used pre-validated, commercially available TaqMan Gene Expression Assays from Thermo Fisher Scientific for the following genes: STAT1, STAT3, TNFRSF1A, TLR2, MAP3K8, RELB, SOCS3, NFκB2, TNF, PLIN2, SOCS1 and LIF.

| Gene Symbol | Assay ID | Dye Label |
| --- | --- | --- |
| Tnfrsf1a | Mm00441883_g1 | FAM-MGB |
| Socs3 | Mm01249143_g1 | FAM-MGB |
| Stat1 | Mm01257286_m1 | FAM-MGB |
| Stat3 | Mm01219775_m1 | FAM-MGB |
| Tnf | Mm00443258_m1 | FAM-MGB |
| Tlr2 | Mm00442346_m1 | FAM-MGB |
| Nfkb2 | Mm00479807_m1 | FAM-MGB |
| Relb | Mm00485664_m1 | FAM-MGB |
| Map3k8 | Mm00432637_m1 | FAM-MGB |

### LNP Synthesis

Chicken egg L-α-phosphatidylglycerol (sodium salt) (PG) was purchased from Avanti Polar Lipids, Inc. (AL, USA) and soy Phosphatidylcholine (PC) was a kind gift from Lipoid GmbH (Germany). Particles were prepared using the NanoAssemblr™ Benchtop (Precision Nanosystems, Canada). The organic phase consisted of a lipid mixture and a small molecule drug in ethyl acetate. PG, PC and ethyl acetate were mixed in a 1:10:100 mass ratio. The small molecule drug of choice was added to the above lipid mixture at 5.6 mM final concentration. The aqueous phase consisted of Galectin-3 or p-selectin binding glycolipids dissolved in water, in a concentration of 2mg/ml and 3 mg/ml respectively. Before the formulation the cartridge was prewashed with water and ethyl acetate at a flow rate of 4ml/min, total volume 2ml, and water: ethyl acetate ratio 1:1. Afterwards the aqueous and organic phases were loaded into separate syringes and pumped through the NanoAssemblr cartridge at a total flow of 8ml/min and volume ratio of 8.5:1.5. For fluorescent LNPs used in biodistribution studies, after the microfluidic mixing, 1,2-dioleoyl-sn-glycero-3-phosphoethanolamine-N-(Cyanine 5) purchased from Avanti Polar Lipids, Inc. (AL, USA) was dissolved in ethanol to a stock concentration of 5mg/ml and added to the LNP mixture at a final concentration of 0.1mg/ml. The resulting mixture was briefly vortexed and placed into a rotor evaporator to remove the organic solvent. For in vivo administration, D(+)-sucrose was added to the mixture to a final concentration of 10% (w/v) and the resulting mixture was filtered through a 0.2 µm filter.

### LNP characterization

#### Size measurements

Size (including number, volume, and intensity mean) as well as PDI were determined by dynamic light-scattering (DLS) measurements acquired with a Malvern Zetasizer Nano ZS. Nanoparticles were prepared for DLS by diluting 1:100 in saline solution. Size and homogeneity of nanoparticles was further characterized by cryogenic electron microscopy (cryo EM) modified from methods already described previously.^100^ Briefly, nanoparticles were imaged using a Titan Krios G2 microscope (ThermoFisher, MA, USA) equipped with a K3 detector (Gatan, CA, USA).

#### Drug concentration and loading

Drug concentration in the LNPs was quantified using high-performance liquid chromatography (HPLC). Nanoparticles were prepared for HPLC analysis by first vortexing the LNP suspension with a 50/50 methanol/saline solution in a 1:4 (LNP:solution) ratio to release the drug from the nanoparticles. This was further diluted in methanol for a final drug dilution of 1:50. Samples were then assessed on an Agilent 1260 Infinity II HPLC system with an InfinityLab Poroshell 120 EC-C18, 4.6 × 75 mm^2^, 2.7 µm analytical LC column. The mobile phase consisted of acetonitrile and/or deionized water, each containing 0.1% trifluoroacetic acid. Chromatographic separation was achieved by gradient elution with acetonitrile (0–95%) at a flow rate of 1 ml min^−1^. For each small molecule drug used, a standard curve was also prepared and used for quantification. For each drug, a single peak was observed at their corresponding retention times and absorbances, as detailed in **Supplementary Table 6**. Prior to HPLC analysis, the UV–vis absorbance spectrum for each drug tested was obtained using a UV/vis/nIR spectrophotometer (Jasco V-670, Tokyo, Japan) to select the appropriate absorbance wavelength for HPLC measurements.

Drug loading to determine the mass ratio of drug to drug-loaded nanoparticles was calculated by filtering the LNPs using 50 kD MWCO dialysis membrane (Spectra/Por® 6 Standard RC tubing, 28 mm width, 1 m length) into dialysate composed of double distilled water at 4 °C. The LNP suspension was then subject to lyophilization for 16 hours to determine the dry weight of LNPs. After resuspending the dried LNPs in methanol, HPLC analysis to quantify the mass of drug was then conducted as previously described.

#### Stability measurements

The stability of drug-loaded Gal-3– and P-sel–targeted LNP formulations was evaluated by dynamic light scattering (DLS). Ruxolitinib-Gal-3, WP1066-Gal-3, TPCA-1-P-sel, and Tofacitinib-P-sel LNPs were diluted in PBS or 25% adult bovine serum (ABS) in PBS and stored at 4 °C, room temperature, or 37 °C under continuous magnetic stirring. At each time point (Days 0–7), aliquots were collected and analyzed for hydrodynamic diameter and polydispersity index (PDI) using a Malvern Zetasizer Nano (Malvern Panalytical). Three technical measurements were recorded for each sample at every time point, and mean values were plotted.

#### Drug release

The drug release kinetics of LNPs was characterized by performing dialysis. LNPs were confined within a pre-wetted dialysis membrane compatible with hydrophobic drugs (Spectra/Por® 6 Standard RC tubing, MWCO 50 kD, 28 mm width, 1 m length) which allowed the release of free drug into the dialysate outside the membrane. The dialysate was comprised of equal parts PBS and adult bovine serum (ABS) (ThermoFisher, MA, USA) to simulate physiological conditions, with 1% DMSO (Alfa Aesar, MA, USA) and 1% Tween-80 (MP Biomedicals, France) added to increase drug solubility and thus ensure sink conditions were satisfied. LNPs within the dialysis membrane were in a solution of the same composition. Moreover, the volume of LNPs to dialysate was 1:100, per the recommendations of the Spectra/Por User Guide. The dialysis system was maintained at a constant 37°C and mixed at 75 rpm by a Corning® 5 x 7 Inch Top PC-420D Stirring Hot Plate. Over a 48-hour period, 150 µL was sampled from within the dialysis bag at each timepoint.

Quantification of the drug remaining within the dialysis bag was conducted by HPLC. LNPs withdrawn from the dialysis bag were diluted 1:50 in methanol to precipitate serum proteins, which were then spun down at 30 000 rcf for 6 min. The supernatant was subsequently filtered through 0.2 µm Whatman syringeless filter devices made of polytetrafluoroethylene (PTFE) (Cytiva, MA, USA). HPLC analysis was then conducted as previously described.

### LNP biodistribution studies

GVHD (1M T cells) was induced as described above. The study groups were: GVHD (1M) – 5mice, Healthy -5 mice, Uninjected control (for background fluorescence normalization)- 1 healthy and 1 GVHD mouse. On day 6 after BMT, Cy5-conjugated LNPs were injected by tail IV into the mice, and mice were sacrificed 24 h later. Liver, kidneys, heart, lung and intestines were harvested and washed briefly with PBS. Tissues were placed on a petri dish (tissues from the same mouse were grouped together), marked with a permanent marker for identification. Tissues were imaged in an Invitrogen iBright FL1500 Imaging System, in fluorescence mode, choosing the Cy5 excitation and detection wavelengths. Various durations of exposure time were tested, until 1s was chosen for all samples as it produced no saturated pixels but enough intensity for quantification. Mouse intestines were highly auto-fluorescent (data from the uninjected control) so they were excluded from subsequent images as they were saturating the receptor signal. All other mouse organs from uninjected mice were always placed in the same imaging plane for autofluorescence normalization. Average fluorescence intensity normalized by the area was quantified by Fiji.

### Efficacy Study 1 (1M T cells, 7 days)

Ruxolitinib, WP1066: GVHD (1M T cells) in mice was induced as described above. Mice were treated with vehicle (V groups), 20mg/kg free drug (FD groups) or drug encapsulated in Gal-3 LNPs (LNP groups), administered intraperitoneally on days 2, 4 and 6 after BMT. Mice were scored on day 7, then euthanized and blood and organs were harvested for analysis. Ruxolitinib was dissolved in 5% DMSO and 2% Tween80 and WP1066 was dissolved in 10% DMSO and 40% PEG300.

Tofacitinib, TPCA-1: GVHD (1M T cells) in mice was induced as described above. Mice were treated with vehicle (V groups), 20mg/kg free drug or drug encapsulated in Gal-3 LNPs, administered intraperitoneally daily on days 2 to 6 after BMT. The last dose of TPCA-1 was not administered because mice died on day 5 (see notes on on-target toxicity of NFκB inhibition above). Mice were scored on day 7, then euthanized and blood and organs were harvested for analysis. Tofacitinib was dissolved in 5% DMSO and TPCA-1 was dissolved in 5% DMSO and 30% PEG300.

### Efficacy Study 2 (2M T cells, 14 days)

GVHD (2M T cells) in mice was induced as described above. Mice were divided in day 7 groups and day 14 groups. Each day group consisted of vehicle (V), free drug (FD) and LNP (R-LNP) groups, that received free vehicle, free Ruxolitinib or Ruxolitinib Gal3- binding LNPs, daily from day 2 after BMT till the day of sacrifice. Day 7 and day 14 groups were sacrificed on the respective days and blood and organs were collected for analysis.

### Long-Term Tolerance and Toxicity Assessment (14 and 30 days)

To assess the long-term systemic toxicity and tolerance of the drug formulations, healthy female BALB/c mice (8-10 weeks old) were randomized into three groups (n=5 per group, treated; n=3, healthy): healthy, Ruxolitinib free drug (R-FD), and Ruxolitinib Galectin-3-targeted lipid nanoparticles (R-LNP). The R-FD and R-LNP groups received Ruxolitinib at a dose of 20 mg/kg, administered by IP injection every other day. The R-LNP formulation and the R-FD groups were prepared as described in the LNP Synthesis section and Efficacy Study. Mice were monitored daily for clinical signs of toxicity and their body weights were measured twice weekly. Mice were euthanized if weight loss exceeded the humane endpoint of 20% of their initial body weight. On days 14 and 30, mice from each respective timepoint were euthanized and kidney, liver, and lung were collected, fixed in paraformaldehyde, and processed for routine H&E staining and histopathological evaluation by a blinded pathologist.

### Reporting Summary

Further information on research design is available in the Nature Research Reporting Summary linked to this article.

## Data Availability

The main data supporting the results in this study are available within the paper and its Supplementary Information. The raw sequencing data have been deposited at the NCBI Gene Expression Omnibus (GEO) under accession number: GSE268861. All data generated or analyzed during the study are available from the corresponding author on reasonable request.

## Acknowledgements

We thank Dr. Doris Ponce for her help with a patient chart evaluation and Ilinca Ariescu for her help with qPCR. We thank the following core facilities: Molecular Cytology and Microscopy at MSKCC (special acknowledgement to Fan Ning, for help with sectioning), CryoEM Facility (Jason De La Cruz), Integrated Genomics Operation Core and the Center for Translational Pathology at Weill Cornell Medicine (TaoTao Zhang and Bing He). Figures 7A and 8A were created with BioRender.com.

## Author contributions

Conceptualization: MP, DAH, EAJ Methodology: MP

Investigation: MP, ANS, KVA, AK, JN, KP, EG, BG, RG, SR

Visualization: MP, ANS, AK Resources: MP, TM, SS

Writing – original draft: MP, ANS

Writing – review and editing: MP, DAH, EAJ, MVDB, AMH

Funding acquisition: DAH, MVDB

## References

1. Blazar BR, Murphy WJ, Abedi M. Advances in graft-versus-host disease biology and therapy. Nat Rev Immunol 12, 443–458 (2012).

2. Zeiser R. Advances in understanding the pathogenesis of graft-versus-host disease. Br J Haematol 187, 563–572 (2019).

3. Ferrara JL, Levine JE, Reddy P, Holler E. Graft-versus-host disease. Lancet 373, 1550–1561 (2009).

4. DeFilipp Z, et al. Nonrelapse mortality among patients diagnosed with chronic GVHD: an updated analysis from the Chronic GVHD Consortium. Blood Advances 5, 4278–4284 (2021).

5. Holtan SG, et al. Disease progression, hospital readmissions, and clinical outcomes for patients with steroid-refractory acute graft-versus-host disease: A multicenter, retrospective study. Bone marrow transplantation 57, 1399–1404 (2022).

6. Kersting S, Dorp SV, Theobald M, Verdonck LF. Acute renal failure after nonmyeloablative stem cell transplantation in adults. Biol Blood Marrow Transplant 14, 125–131 (2008).

7. Grigoryev DN, Liu M, Hassoun HT, Cheadle C, Barnes KC, Rabb H. The local and systemic inflammatory transcriptome after acute kidney injury. J Am Soc Nephrol 19, 547–558 (2008).

8. Hingorani SR, Guthrie K, Batchelder A, Schoch G, Aboulhosn N, Manchion J, McDonald GB. Acute renal failure after myeloablative hematopoietic cell transplant: Incidence and risk factors. Kidney International 67, 272–277 (2005).

9. Hingorani SR, Seidel K, Lindner A, Aneja T, Schoch G, McDonald G. Albuminuria in hematopoietic cell transplantation patients: prevalence, clinical associations, and impact on survival. Biol Blood Marrow Transplant 14, 1365–1372 (2008).

10. Kagoya Y, Kataoka K, Nannya Y, Kurokawa M. Pretransplant predictors and posttransplant sequels of acute kidney injury after allogeneic stem cell transplantation. Biol Blood Marrow Transplant 17, 394–400 (2011).

11. Lopes JA, et al. Acute renal failure following myeloablative autologous and allogeneic hematopoietic cell transplantation. Bone Marrow Transplant 38, 707 (2006).

12. Mae H, et al. Early renal injury after myeloablative cord blood transplantation in adults. Leukemia & Lymphoma 49, 538–542 (2008).

13. Parikh CR, et al. Comparison of ARF after myeloablative and nonmyeloablative hematopoietic cell transplantation. Am J Kidney Dis 45, 502–509 (2005).

14. Parikh CR, Yarlagadda SG, Storer B, Sorror M, Storb R, Sandmaier B. Impact of acute kidney injury on long-term mortality after nonmyeloablative hematopoietic cell transplantation. Biol Blood Marrow Transplant 14, 309–315 (2008).

15. Satwani P, et al. Risk factors associated with kidney injury and the impact of kidney injury on overall survival in pediatric recipients following allogeneic stem cell transplant. Biol Blood Marrow Transplant 17, 1472–1480 (2011).

16. Tokgoz B, et al. Acute renal failure after myeloablative allogeneic hematopoietic stem cell transplantation: incidence, risk factors, and relationship with the quantity of transplanted cells. Ren Fail 32, 547–554 (2010).

17. Weiss AS, Sandmaier BM, Storer B, Storb R, McSweeney PA, Parikh CR. Chronic kidney disease following non-myeloablative hematopoietic cell transplantation. Am J Transplant 6, 89–94 (2006).

18. Shingai N, et al. Early-onset acute kidney injury is a poor prognostic sign for allogeneic SCT recipients. Bone Marrow Transplantation 50, 1557–1562 (2015).

19. Abramson MH, et al. Acute Kidney Injury in the Modern Era of Allogeneic Hematopoietic Stem Cell Transplantation. Clin J Am Soc Nephrol 16, 1318–1327 (2021).

20. Flores FX, et al. Continuous renal replacement therapy (CRRT) after stem cell transplantation. A report from the prospective pediatric CRRT Registry Group. Pediatr Nephrol 23, 625–630 (2008).

21. Gaziev J, et al. Late-Onset Hemorrhagic Cystitis in Children after Hematopoietic Stem Cell Transplantation for Thalassemia and Sickle Cell Anemia: A Prospective Evaluation of Polyoma (BK) Virus Infection and Treatment with Cidofovir. Biology of Blood and Marrow Transplantation 16, 662–671 (2010).

22. Renaghan AD, Jaimes EA, Malyszko J, Perazella MA, Sprangers B, Rosner MH. Acute Kidney Injury and CKD Associated with Hematopoietic Stem Cell Transplantation. Clin J Am Soc Nephrol 15, 289–297 (2020).

23. Piñana JL, et al. Study of Kidney Function Impairment after Reduced-Intensity Conditioning Allogeneic Hematopoietic Stem Cell Transplantation. A Single-Center Experience. Biology of Blood and Marrow Transplantation 15, 21–29 (2009).

24. Liu H, Li YF, Liu BC, Ding JH, Chen BA, Xu WL, Qian J. A multicenter, retrospective study of acute kidney injury in adult patients with nonmyeloablative hematopoietic SCT. Bone Marrow Transplantation 45, 153–158 (2010).

25. Hingorani S, et al. Urinary Elafin and Kidney Injury in Hematopoietic Cell Transplant Recipients. Clinical Journal of the American Society of Nephrology 10, 12–20 (2015).

26. Raina R, et al. Acute kidney injury in pediatric hematopoietic cell transplantation: critical appraisal and consensus. Pediatric Nephrology 37, 1179–1203 (2022).

27. Hingorani S. Renal Complications of Hematopoietic-Cell Transplantation. N Engl J Med 374, 2256–2267 (2016).

28. Choe HK, et al. Kidney Dysfunction Post-Allogeneic Transplant: High Incidence of TMA and Kidney GvHD. Blood 130, 5500 (2017).

29. Lopes J, Jorge S, Neves M. Acute kidney injury in HCT: an update. Bone Marrow Transplantation 51, 755–762 (2016).

30. Schmid PM, et al. Acute Renal Graft-Versus-Host Disease in a Murine Model of Allogeneic Bone Marrow Transplantation. Cell Transplant 26, 1428–1440 (2017).

31. Amin R, et al. The kidney injury caused by the onset of acute graft-versus-host disease is associated with down-regulation of alphaKlotho. Int Immunopharmacol 78, 106042 (2020).

32. Ma Q, Li D, Vasquez HG, You MJ, Afshar-Kharghan V. Kidney Injury in Murine Models of Hematopoietic Stem Cell Transplantation. Biol Blood Marrow Transplant 25, 1920–1924 (2019).

33. Garnett C, Apperley JF, Pavlu J. Treatment and management of graft-versus-host disease: improving response and survival. Ther Adv Hematol 4, 366–378 (2013).

34. Isberner N, et al. Ruxolitinib exposure in patients with acute and chronic graft versus host disease in routine clinical practice-a prospective single-center trial. Cancer Chemother Pharmacol 88, 973–983 (2021).

35. Westin JR, et al. Steroid-refractory acute GVHD: predictors and outcomes. Advances in hematology 2011, 601953 (2011).

36. Zeiser R, et al. Ruxolitinib for Glucocorticoid-Refractory Chronic Graft-versus-Host Disease. N Engl J Med 385, 228–238 (2021).

37. Zeiser R, et al. Ruxolitinib in corticosteroid-refractory graft-versus-host disease after allogeneic stem cell transplantation: a multicenter survey. Leukemia 29, 2062–2068 (2015).

38. Incyte Corporation. JAKAFI (ruxolitinib) tablets, for oral use: full prescribing information. (ed U.S. National Library of Medicine D). Revised June 2025 edn (2025).

39. Ytterberg SR, et al. Cardiovascular and cancer risk with tofacitinib in rheumatoid arthritis. New England Journal of Medicine 386, 316–326 (2022).

40. Patel DA, Crain M, Pusic I, Schroeder MA. Acute Graft-versus-Host Disease: An Update on New Treatment Options. Drugs 83, 893–907 (2023).

41. Pickkers P, Murray PT, Ostermann M. New drugs for acute kidney injury. Intensive Care Med 48, 1796–1798 (2022).

42. Trac N, Ashraf A, Giblin J, Prakash S, Mitragotri S, Chung EJ. Spotlight on Genetic Kidney Diseases: A Call for Drug Delivery and Nanomedicine Solutions. ACS Nano 17, 6165–6177 (2023).

43. Summerlin N, Soo E, Thakur S, Qu Z, Jambhrunkar S, Popat A. Resveratrol nanoformulations: challenges and opportunities. Int J Pharm 479, 282–290 (2015).

44. Cote JM, Murray PT, Rosner MH. New drugs for acute kidney injury. Curr Opin Crit Care 26, 525–535 (2020).

45. Williams RM, Jaimes EA, Heller DA. Nanomedicines for kidney diseases. Kidney Int 90, 740–745 (2016).

46. Huang Y, Wang J, Jiang K, Chung EJ. Improving kidney targeting: The influence of nanoparticle physicochemical properties on kidney interactions. J Control Release 334, 127–137 (2021).

47. Wang J, Masehi-Lano JJ, Chung EJ. Peptide and antibody ligands for renal targeting: nanomedicine strategies for kidney disease. Biomater Sci 5, 1450–1459 (2017).

48. Miyata M, Ichikawa K, Matsuki E, Watanabe M, Peltier D, Toubai T. Recent Advances of Acute Kidney Injury in Hematopoietic Cell Transplantation. Front Immunol 12, 779881 (2021).

49. Filipovich AH, et al. National Institutes of Health consensus development project on criteria for clinical trials in chronic graft-versus-host disease: I. Diagnosis and staging working group report. Biology of blood and marrow transplantation 11, 945–956 (2005).

50. Daniele N, et al. Overview of T-cell depletion in haploidentical stem cell transplantation. Blood Transfusion 10, 264 (2012).

51. Saad A, Lamb L. Ex vivo T-cell depletion in allogeneic hematopoietic stem cell transplant: past, present and future. Bone marrow transplantation 52, 1241–1248 (2017).

52. Mackay LK, et al. The developmental pathway for CD103+ CD8+ tissue-resident memory T cells of skin. Nature immunology 14, 1294–1301 (2013).

53. Takamura S. Divergence of tissue-memory T cells: distribution and function-based classification. Cold Spring Harbor Perspectives in Biology 12, a037762 (2020).

54. Diaz MF, et al. Injury intensifies T cell mediated graft-versus-host disease in a humanized model of traumatic brain injury. Scientific Reports 10, 10729 (2020).

55. Davies SP, et al. Expression of E-cadherin by CD8+ T cells promotes their invasion into biliary epithelial cells. Nature Communications 15, 853 (2024).

56. Jansen SA, Nieuwenhuis EE, Hanash AM, Lindemans CA. Challenges and opportunities targeting mechanisms of epithelial injury and recovery in acute intestinal graft-versus-host disease. Mucosal immunology 15, 605–619 (2022).

57. Washington K, Jagasia M. Pathology of graft-versus-host disease in the gastrointestinal tract. Human pathology 40, 909–917 (2009).

58. Maunsbach AB, Marples D, Chin E, Ning G, Bondy C, Agre P, Nielsen S. Aquaporin-1 water channel expression in human kidney. Journal of the American Society of Nephrology 8, 1–14 (1997).

59. Bedford JJ, Leader JP, Walker RJ. Aquaporin expression in normal human kidney and in renal disease. Journal of the American Society of Nephrology 14, 2581–2587 (2003).

60. Ho KM, et al. Altered expression of aquaporin-2 in human explants with chronic renal allograft dysfunction. BJU international 95, 1104–1108 (2005).

61. Stein-Thoeringer CK, et al. Lactose drives Enterococcus expansion to promote graft-versus-host disease. Science 366, 1143–1149 (2019).

62. Mishra J, Ma Q, Kelly C, Mitsnefes M, Mori K, Barasch J, Devarajan P. Kidney NGAL is a novel early marker of acute injury following transplantation. Pediatric Nephrology 21, 856–863 (2006).

63. Adedeji AO, et al. The Utility of Novel Urinary Biomarkers in Mice for Drug Development Studies. Int J Toxicol 40, 15–25 (2021).

64. Kaucsár T, et al. Urine/plasma neutrophil gelatinase associated lipocalin ratio is a sensitive and specific marker of subclinical acute kidney injury in mice. PloS one 11, e0148043 (2016).

65. Parmaksız G, Noyan A, Dursun H, İnce E, Anarat R, Cengiz N. Role of new biomarkers for predicting renal scarring in vesicoureteral reflux: NGAL, KIM-1, and L-FABP. Pediatric nephrology 31, 97–103 (2016).

66. Schmid PM, et al. Acute renal graft-versus-host disease in a murine model of allogeneic bone marrow transplantation. Cell Transplantation 26, 1428–1440 (2017).

67. Lai HY, Chou TY, Tzeng CH, Lee OKS. Cytokine Profiles in Various Graft-Versus-Host Disease Target Organs Following Hematopoietic Stem Cell Transplantation. Cell Transplantation 21, 2033–2045 (2012).

68. Subramanian A, et al. Gene set enrichment analysis: a knowledge-based approach for interpreting genome-wide expression profiles. Proc Natl Acad Sci U S A 102, 15545–15550 (2005).

69. Saliu TP, et al. Serum Amyloid A3 Promoter-Driven Luciferase Activity Enables Visualization of Diabetic Kidney Disease. Int J Mol Sci 23, 899 (2022).

70. Ferrara J, Antin J. Thomas’ hematopoietic cell transplantation.). Blackwell Malden, MA (2004).

71. Schroeder MA, Choi J, Staser K, DiPersio JF. The Role of Janus Kinase Signaling in Graft-Versus-Host Disease and Graft Versus Leukemia. Biol Blood Marrow Transplant 24, 1125–1134 (2018).

72. Takashima S, et al. T cell–derived interferon-γ programs stem cell death in immune-mediated intestinal damage. Science immunology 4, eaay8556 (2019).

73. Lawrence T. The nuclear factor NF-kappaB pathway in inflammation. Cold Spring Harb Perspect Biol 1, a001651 (2009).

74. Shamay Y, et al. P-selectin is a nanotherapeutic delivery target in the tumor microenvironment. Science Translational Medicine 8, 345ra387–345ra387 (2016).

75. Lorenzon P, Vecile E, Nardon E, Ferrero E, Harlan J, Tedesco F, Dobrina A. Endothelial cell E-and P-selectin and vascular cell adhesion molecule-1 function as signaling receptors. Journal of Cell Biology 142, 1381–1391 (1998).

76. Henderson NC, et al. Galectin-3 expression and secretion links macrophages to the promotion of renal fibrosis. Am J Pathol 172, 288–298 (2008).

77. Sun K, et al. Differential effects of proteasome inhibition by bortezomib on murine acute graft-versus-host disease (GVHD): delayed administration of bortezomib results in increased GVHD-dependent gastrointestinal toxicity. Blood 106, 3293–3299 (2005).

78. MacDonald KPA, et al. Effector and regulatory T-cell function is differentially regulated by RelB within antigen-presenting cells during GVHD. Blood 109, 5049–5057 (2007).

79. Fu Y-Y, et al. T cell recruitment to the intestinal stem cell compartment drives immune-mediated intestinal damage after allogeneic transplantation. Immunity 51, 90–103. e103 (2019).

80. Edelstein CL. Biomarkers in Acute Kidney Injury. Biomarkers of Kidney Disease, 2nd *Edition*, 241–315 (2017).

81. Troxell ML, Pilapil M, Miklos DB, Higgins JP, Kambham N. Renal pathology in hematopoietic cell transplantation recipients. Modern Pathology 21, 396–406 (2008).

82. Girsberger M, Halter JP, Hopfer H, Dickenmann M, Menter T. Kidney pathology after hematologic cell transplantation—a single-center observation study of indication biopsies and autopsies. Biology of Blood and Marrow Transplantation 24, 571–580 (2018).

83. Nishiyama J, et al. Up-regulation of galectin-3 in acute renal failure of the rat. The American journal of pathology 157, 815–823 (2000).

84. Liu L, et al. The KLF4/Galectin-3 cascade is a key determinant of tubular cell death and acute kidney injury. International journal of biological sciences 21, 5802 (2025).

85. Boutin L, Legrand M, Sadoune M, Mebazaa A, Gayat E, Chadjichristos CE, Dépret F. Elevated plasma Galectin-3 is associated with major adverse kidney events and death after ICU admission. Critical Care 26, 13 (2022).

86. Wang F, et al. The potential roles of galectin-3 in AKI and CKD. Frontiers in Physiology 14, 1090724 (2023).

87. O’Seaghdha CM, Hwang S-J, Ho JE, Vasan RS, Levy D, Fox CS. Elevated galectin-3 precedes the development of CKD. Journal of the American Society of Nephrology 24, 1470–1477 (2013).

88. Ou S-M, et al. Identification of galectin-3 as potential biomarkers for renal fibrosis by RNA-sequencing and clinicopathologic findings of kidney biopsy. Frontiers in medicine 8, 748225 (2021).

89. Barenholz YC. Doxil®—The first FDA-approved nano-drug: Lessons learned. Journal of controlled release 160, 117–134 (2012).

90. Van Hoogevest P, Wendel A. The use of natural and synthetic phospholipids as pharmaceutical excipients. European journal of lipid science and technology 116, 1088–1107 (2014).

91. Kang W, et al. Multiplex Spatial Protein Detection by Combining Immunofluorescence with Immunohistochemistry. In: Signal Transduction Immunohistochemistry: Methods and Protocols). Springer (2022).

92. Eisner C, et al. Major contribution of tubular secretion to creatinine clearance in mice. Kidney Int 77, 519–526 (2010).

93. Schreiber A, et al. Transcutaneous measurement of renal function in conscious mice. Am J Physiol Renal Physiol 303, F783–788 (2012).

94. Andrews S. FastQC: a quality control tool for high throughput sequence data.). Babraham Bioinformatics, Babraham Institute, Cambridge, United Kingdom (2010).

95. Bolger AM, Lohse M, Usadel B. Trimmomatic: a flexible trimmer for Illumina sequence data. Bioinformatics 30, 2114–2120 (2014).

96. Dobin A, et al. STAR: ultrafast universal RNA-seq aligner. Bioinformatics 29, 15–21 (2013).

97. Liao Y, Smyth GK, Shi W. featureCounts: an efficient general purpose program for assigning sequence reads to genomic features. Bioinformatics 30, 923–930 (2014).

98. Eissa N, Kermarrec L, Hussein H, Bernstein CN, Ghia JE. Appropriateness of reference genes for normalizing messenger RNA in mouse 2,4-dinitrobenzene sulfonic acid (DNBS)-induced colitis using quantitative real time PCR. Sci Rep 7, 42427 (2017).

99. Muñoz JJ, et al. Ppia is the most stable housekeeping gene for qRT-PCR normalization in kidneys of three Pkd1-deficient mouse models. Scientific Reports 11, 19798 (2021).

100. Roberts S, et al. Sonophore-enhanced nanoemulsions for optoacoustic imaging of cancer. Chemical Science 9, 5646–5657 (2018).

