## Supplementary Materials for "Galectin-3 is a Nanotherapeutic Target in Graft-versus-Host Disease Mediated Kidney Injury"

### Supplementary Materials
#### for

**Galectin-3 is a Nanotherapeutic Target in Graft-versus-Host Disease Mediated Kidney Injury**

Panagiotakopoulou Magdalini *et al.*

**The PDF file includes:**
Supplementary Discussion
Supplementary Figures 1 to 26
Supplementary Tables 1 to 8

#### Supplementary Discussion

##### Note 1: Nanoparticle Trafficking and Target Engagement in GVHD-Mediated Kidney Injury

In GVHD, the renal microvascular endothelium is profoundly activated and structurally altered by Th1 cytokines, particularly IFN-γ and TNF-α. These cytokines induce endothelial activation, junctional remodeling, and cytoskeletal contraction, resulting in increased permeability of peritubular capillaries and enhanced transendothelial transport of cells and macromolecules[1,2, 3]). Inflammatory cytokine–driven endothelial activation is also known to promote caveolae-mediated transcytosis and paracellular leak, mechanisms that permit nanoparticle extravasation into inflamed tissues independent of classical receptor-mediated uptake[4 5, 6].

Once beyond the endothelium, renal epithelial access is facilitated by the unique vascular–epithelial architecture of the kidney. Tubular epithelial cells are separated from the peritubular capillary blood by their basolateral membrane and tubular basement membrane, a very thin interstitial space, and the peritubular capillary basement membrane and endothelium[7]. In immune-mediated kidney injury, including GVHD, the peritubular-interstitial interface becomes inflamed, with prominent T cell infiltration, endothelial activation characterized by upregulation of MHC class II and adhesion molecules, and tubular epithelial injury with apoptosis. This inflammatory milieu is associated with increased microvascular permeability through disruption of endothelial barrier function, which may render this compartment more accessible to circulating factors and potentially to nanomaterials[8, 9, 10].

Within this inflamed interstitial and epithelial environment, galectin-3 provides a distinct, non-endocytic mechanism for tissue localization and retention. Galectin-3 is a multivalent β-galactoside–binding lectin that forms extracellular galectin–glycan lattices by crosslinking glycosphingolipids and glycoproteins on epithelial and stromal surfaces. This so-called G-lect mechanism enables surface retention, clustering, and gradual internalization of glycan-decorated particles without requiring a canonical receptor[11, 12]. Importantly, cellular uptake via this multivalent Gal-3/GL-Lect interaction triggers clathrin-independent endocytosis. Unlike conventional ligand-targeted nanocarriers that are rapidly routed to the lysosome, this pathway facilitates superior endosomal escape and reduces lysosomal degradation of the payload, maximizing the cytosolic delivery and functional efficacy of the therapeutic cargo. In inflamed tissues, galectin-3 is further enriched at basolateral epithelial surfaces and within the extracellular matrix, where it functions as a damage-associated lectin scaffold.

Finally, although P-selectin- and galectin-3-targeted nanoparticles bind to different cellular compartments, they converge functionally on the same inflammatory signaling axis. P-selectin targeting primarily modulates endothelial activation and leukocyte recruitment, whereas galectin-3 targeting localizes drug delivery to epithelial and interstitial compartments where cytokine amplification and tissue injury occur. In GVHD, both compartments exhibit coordinated JAK/STAT activation, providing a mechanistic explanation for the comparable therapeutic efficacy observed despite distinct binding targets.

##### Note 2: P-selectin Targeting and Potential Platelet Interactions

P-selectin is expressed on activated platelets and inflamed endothelial cells, where it mediates adhesive interactions with leukocytes via PSGL-1. While it does not directly induce platelet activation, it plays a key role in stabilizing platelet–leukocyte aggregates and promoting inflammatory cell recruitment. Accordingly, multivalent presentation of P-selectin-binding ligands on nanoparticles could theoretically enhance these interactions.

In our study, however, we did not observe evidence of exacerbated inflammation in vivo. Sulfatide-targeted nanoparticles did not increase renal $CD3^+$ T cell infiltration or worsen histological injury compared to untreated GVHD controls, and repeated dosing was not associated with overt systemic intolerance. The low molar fraction of sulfatide in the formulation may limit extensive multivalent engagement while preserving targeting to activated endothelium. We acknowledge that platelet activation and aggregation were not directly assessed and represent an area for future investigation.

| Pt | T between allo-HCT and Bx (months) | Gender | Age (y) | allo-HCT indication | Preconditioning | GVHD prophylaxis | Historical | | | Around the time of kidney biopsy | | | |
|---|---|---|---|---|---|---|---|---|---|---|---|---|---|
| | | | | | | | Extra-renal GVHD | Extra-renal GVHD location | Extra-renal GVHD classification | Extra-renal GVHD | Extra-renal GVHD classification | Extra-renal GVHD location | Calci-neurin Inhibitor |
| **PT_1** | 4 | M | 61 | AITL | TBI, CTX, fludarabine, No ATG, No KGF | Tacrolimus, Sirolimus, Mini-methotrexate | yes | Skin | Acute | **yes** | **Acute** | skin | yes |
| **PT_2** | 12 | M | 59 | AML | Busulfan, Melphalan, Fludarabine , Rabbit anti-thymocyte globulin, No KGF | none; TCD transplant | yes | Skin | acute | **yes** | **Acute, recurrent** | skin | no |
| **PT_3** | 11 | F | 51 | ALL | TBI, CTX, Fludarabine, Thiotepa,No ATG,No KGF | cyclosporine, mycophenolate mofetil | yes | Skin | Acute, persistent/ recurrent | **yes** | **Acute, persistent/ recurrent** | skin | yes |
| **PT_4** | 9 | M | 46 | CMML | TBI, fludarabine, melphalan | na | yes | Skin, eye | Acute | **yes** | **Acute** | skin, eye | no |
| **PT_5** | 9 | F | 60 | AML | TBI, CTX, Thiotepa, Fludarabine, No ATG, No KGF | Cyclosporine, MMF | yes | UGI, LGI | Acute, recurrent | **yes** | **Acute, persistent/ recurrent** | UGI, LGI | yes |
| **PT_6** | 9 | M | 64 | AML | melphalan + fludarabine | Tacrolimus and prednisone | yes | Skin, UGI | Acute, recurrent | **yes** | **Acute, recurrent** | skin | yes |
| **PT_7** | 14 | M | 75 | ALAL | TBI, Pentostatin, CTX (no ATG, no KGF) | Tacrolimus, mini-methotrexate | yes | Skin | Acute, late onset | **yes** | **Overlap Syndrome** | Skin, mouth , lungs | No |
| **PT_8** | 12 | M | 66 | AML | Melphalan, Fludarabine, No ATG | Tacrolimus, Mini-methotrexate | yes | UGI, skin, oral, eye | Acute, chronic | **yes** | **Chronic** | skin, oral, eye | no |
| **PT_9** | 12 | F | 78 | MDS | Melphalan, Fludarabine, No ATG | Tacrolimus, Mini-methotrexate | yes | UGI, skin | Chronic | **yes** | **Chronic** | UGI, skin | no |
| **PT_10** | 9 | M | 57 | CMML | 1st transplant: TBI, Thiotepa, CTX, eATG, KGF. 2nd transplant: Fludarabine, CTX, rATG, KGF | 2nd transplant: none; TCD transplant | yes | skin | acute, then chronic | **yes** | **Chronic** | skin | no |
| **PT_11** | 12 | F | 66 | AML | TBI, fludarabine, melphalan | na | yes | skin | Chronic | **yes** | **Chronic** | skin | no |

**Supplementary Table 1 |** Patients Demographics and allo-HCT Characteristics. AML (Acute Myeloid Leukemia), ALL (Acute Lymphoblastic Leukemia), ALAL (Acute Leukemia of Ambiguous Lineage), MDS (Myelodysplastic Syndromes), CMML (Chronic Myelomonocytic Leukemia), NHL (Non-Hodgkin Lymphoma), AITL (Angioimmunoblastic T cell Lymphoma), AA (Aplastic Anemia), CHL (Classical Hodgkin Lymphoma), DLBCL (Diffuse Large B-cell Lymphoma), UGI (Upper Gastrointestinal), LGI (Lower Gastrointestinal), TBI (Total Body Irradiation), ATG (Anti-thymocyte Globulin), KGF (Keratinocyte Growth Factor), TCD (T cell Depletion), MMF (Mycophenolate mofetil), eATG (Equine Anti-thymocyte Globulin), rATG (Rabbit Anti-thymocyte Globulin).

| Pt | T between allo-HCT and Bx (months) | Gender | Age (y) | allo-HCT indication | Preconditioning | GVHD prophylaxis | Historical | | | Around the time of kidney biopsy | | | |
|---|---|---|---|---|---|---|---|---|---|---|---|---|---|
| | | | | | | | Extra-renal GVHD | Extra-renal GVHD location | Extra-renal GVHD classification | Extra-renal GVHD | Extra-renal GVHD classification | Extra-renal GVHD location | Calci-neurin Inhibitor |
| **PT_12** | 9 | M | 51 | NHL | TBI, Cyclophosphamide, Fludarabine, No ATG,No KGF | Cyclosporine, Mycophenolate mofetil | yes | GI | Acute | **Indet.** | **Indeterm./ Intermittent/ Mild Chronic** | GI, skin | no |
| **PT_13** | 2 | M | 45 | AML | Busulfan, Melphalan, Fludarabine, Rabbit ATG, No KGF | none; TCD transplant | yes | skin | Acute | **Indet.** | **Indeterm.** | skin | no |
| **PT_14** | 16 | F | 67 | AML | Busulfan, Melphalan, Fludarabine, Rabbit ATG, No KGF | Tacrolimus, Mini-methotrexate | yes | UGI, LGI | Acute | **Indet.** | **Indeterm.** | skin, GI | tapered |
| **PT_15** | 11 | M | 58 | ALL | Clofarabine, Melphalan, Thiotepa, Rabbit ATG, No KGF | none; TCD transplant | no | - | - | **no** | **no** | - | no |
| **PT_16** | 11 | M | 64 | AML/ MDS | TBI, Cyclophosphamide, Fludarabine, Thiotepa, No ATG, No KGF | mycophenolate mofetil, cyclosporine | yes | UGI | Acute, recurrent | **yes** | **no** | - | yes |
| **PT_17** | 2 | M | 59 | AML | Busulfan, Melphalan, Fludarabine, Rabbit ATG, No KGF | none; TCD transplant | no | - | - | **no** | **no** | - | no |
| **PT_18** | 3 | F | 21 | AA | TBI, Cyclophosphamide, Fludarabine, Rabbit ATG | Tacrolimus, Mini-methotrexate | no | - | - | **no** | **no** | - | no |
| **PT_19** | 2 | M | 61 | ALL | Busulfan, Fludarabine, No ATG, No KGF | Sirolimus, Mycophenolate mofetil, Post transplant cyclophosphamide | yes | UGI, LGI | Acute, resolved, later developed late acute/chronic | **no** | **no, resolved** | - | no |
| **PT_20** | 16 | M | 69 | MDS | Busulfan, Fludarabine, No ATG, No KGF | Tacrolimus, Mini-methotrexate | yes | UGI, LGI | acute | **no** | **no, quiescent** | - | yes |

**Supplementary Table 1 (continued) |** Patients Demographics and allo-HCT Characteristics. AML (Acute Myeloid Leukemia), ALL (Acute Lymphoblastic Leukemia), ALAL (Acute Leukemia of Ambiguous Lineage), MDS (Myelodysplastic Syndromes), CMML (Chronic Myelomonocytic Leukemia), NHL (Non-Hodgkin Lymphoma), AITL (Angioimmunoblastic T cell Lymphoma), AA (Aplastic Anemia), FL (Follicular Lymphoma), UGI (Upper Gastrointestinal), LGI (Lower Gastrointestinal), TBI (Total Body Irradiation), ATG (Anti-thymocyte Globulin), KGF (Keratinocyte Growth Factor), TCD (T cell Depletion), MMF (Mycophenolate mofetil), eATG (Equine Anti-thymocyte Globulin), rATG (Rabbit Anti-thymocyte Globulin).

| Pt | Full list of medications |
|---|---|
| **PT_1** | Acetaminophen, acyclovir oral, allopurinol, amlodipine besylate, atovaquone, benzocaine/menthol, bumetanide INJ, cefepime INJ, ciprofloxacin, enoxaparin INJ, entecavir, filgrastim subcut, finasteride, furosemide INJ, hydrocortisone INJ with additives, hydromorphone (C-II) oral and INJ, lorazepam INJ (C-IV), methotrexate INJ, methylprednisolone INJ, metoprolol INJ and succinate ext release 24hr and tartrate immediate release, ondansetron INJ and oral, oxycodone (C-II), pantoprazole oral, phenazopyridine, prednisone, prochlorperazine INJ, rituximab INJ, senna, sirolimus, spironolactone, tacrolimus oral and IVCI, vancomycin oral, voriconazole oral and INJ |
| **PT_2** | Acetaminophen, acyclovir, amlodipine besylate, amphotericin B liposome INJ, azacitidine INJ, azithromycin oral and INJ, aztreonam INJ, benzonatate, brincidofovir, cefepime INJ, cefpodoxime, cefuroxime, cyclophosphamide oral, cytarabine INJ, diphenhydramine oral and INJ, entecavir, furosemide, gabapentin, hemorrhoidal, heparin (porcine) IVCI, hydralazine HCl and INJ, imipenem/cilastatin IV, labetalol, lactulose, levofloxacin oral and INJ, loperamide, lorazepam (C-IV) INJ and oral, methotrexate INJ, methylprednisolone INJ, metronidazole oral and INJ, morphine sulfate INJ (C-II), Nystatin, ondansetron INJ and oral, oxycodone (C-II) oral and extended release, oxycontin, pantoprazole INJ and oral, piperacillin/tazobactam INJ, polyethylene glycol 3350, posaconazole delayed release, prednisone, prochlorperazine oral and INJ, rituximab, senna, sildenafil, sulfamethoxazole/trimethoprim, Valganciclovir, Vancomycin INJ and oral, Voriconazole oral and INJ, Zolpidem, immune globulin |
| **PT_3** | Acetaminophen oral, aciclovir, atovaquone, benzocaine/menthol, benzonatate, cyclosporine (neoral) and IVPB, diphenhydramine, hydrocortisone INJ, hydroxyzine oral and INJ, isavuconazonium, lactulose, loratadine, lorazepam INJ (C-IV), Medroxyprogesterone, Meropenem INJ, Methylprednisolone INJ, midostaurin, Mycophenolate mofetil, Olanzapine, Ondansetron oral and INJ, Palonosetron INJ, Pantoprazole, Piperacillin/Tazobactam INJ, Penicillin V potassium, Polyethylene Glycol 3350, Prednisone, Senna, Simethicone, Vancomycin INJ, Voriconazole oral and INJ, Ursodiol, Zolpidem |
| **PT_5** | Acetaminophen, Acetylcysteine, Acyclovir INJ, Amlodipine besylate, Azithromycin, Benzocaine/Menthol, Budesonide, Carvedilol, Ceftriaxone INJ, Clonidine, Cosyntropin INJ, Cyclosporine (neoral) and IVPB, Diphenhydramine, Diphenoxylate/Atropine (C-V), Fentanyl INJ (C-II), Filgrastim, Subcut, Fluconazole, Furosemide INJ, Ganciclovir INJ, Hydralazine INJ, Hydrocortisone INJ, Insulin Sliding Scale ASPART INJ, Labetalol INJ, Levofloxacin, Loperamide, Lorazepam oral and INJ, Meropenem INJ, Methylprednisolone INJ, Metoclopramide HCl INJ, Metoprolol succinate (ext release 24hr and tartrate immediate release), Metronidazole INJ, Morphine sulfate INJ (C-II), Mycophenolate mofetil oral and INJ, Nystatin Oral, Olanzapine, Omeprazole, Ondansetron, Oxycodone (C-II), Pantoprazole INJ and oral, Pentamidine, Phenazopyridine, Piperacillin/Tazobactam INJ, Prednisone, Prochlorperazine INJ, Senna, Simethicone, Ursodiol, Vancomycin INJ, Voriconazole oral and INJ, Zolpidem, immune globulin |
| **PT_6** | Atorvastatin, acyclovir, amlodipine, gabapentin, tobramycin/dexamethasone ophthalmic, ergocalciferol (vitamin D-2), metoprolol, prednisone, sevelamer, sodium bicarbonate, tacrolimus, enoxaparin, ipratropium/albuterol |
| **PT_7** | Lisinopril, Acyclovir, Multivitamin, Triamcinolone topical, Metronidazole topical |
| **PT_8** | Acetaminophen, aciclovir INJ and oral, allopurinol, amlodipine besylate, ampicillin INJ, aspirin, atenolol, atovaquone, baclofen oral, benzocaine/menthol, benzonatate, CAB hydromorphone IV, CAB morphine sulfate IV, ceftriaxone INJ, chlorpromazine INJ, cytarabine INJ, diphenhydramine INJ, fentanyl IV PCA, fluconazole, furosemide INJ, guaifenesin syrup, haloperidol oral and INJ, hydralazine INJ, hydromorphone INJ (C-II), insulin sliding scale aspart (rapid-acting) INJ, insulin sliding scale regular (short-acting) INJ, irbesartan, labetalol INJ and oral, loperamide, lorazepam (C-IV) oral and INJ, losartan, meperidine INJ (C-II), methotrexate INJ, metoclopramide HCl oral and INJ, metolazone, midazolam INJ (C-IV), mirabegron, morphine sulfate INJ (C-II), Nystatin, olanzapine, omeprazole, ondansetron oral and INJ, oxycodone (C-II), oxybutynin, pantoprazole INJ and oral, piperacillin/tazobactam INJ, ramelteon, senna, simethicone, sucralfate, sulfamethoxazole-trimethoprim, tacrolimus oral and IVCI, tamsulosin, torsemide, ursodiol oral and suspension 50mg/ml, vancomycin INJ, voriconazole oral and INJ |
| **PT_9** | Acetaminophen oral and INJ, acyclovir oral, amiodarone IVCI, amlodipine besylate, anastrozole, apixaban, atovaquone, bisacodyl, budesonide, CAB Fentanyl IV, carvedilol, cefepime INJ, ciprofloxacin INJ, dexamethasone INJ, diltiazem INJ, diphenhydramine INJ, fentanyl IV PCA, fluconazole INJ, furosemide, guaifenesin ER, hydralazine INJ, hydromorphone (C-II), isavuconazonium sulfate, letermovir oral, levofloxacin INJ, lisinopril, loperamide, loratadine, lorazepam INJ (C-IV), methotrexate INJ, metoprolol INJ and tartrate immediate release, metronidazole INJ, olanzapine, ondansetron INJ, palonosetron INJ, pantoprazole oral and INJ, penicillin V potassium oral, posaconazole delayed release, prochlorperazine INJ, ramelteon, senna, simethicone, sucralfate, sulfamethoxazole/trimethoprim, tacrolimus oral and IVCI, valganciclovir, vancomycin INJ and oral, voriconazole oral and INJ |
| **PT_10** | From 2nd graft onwards:, Acetaminophen, Acyclovir, Amlodipine, Amoxicillin, Atovaquone, Azithromycin dihydrate, Benzonatate, Cefepime HCl, Codeine Sulfate, Daptomycin, Fentanyl citrate, Guaifenesin/codeine, Hydrochlorothiazide, Linezolid, Metoprolol Succinate, Sennoside, Sucralfate, Voriconazole |

| | |
|---|---|
| **PT_12** | Acetaminophen, acyclovir, amlodipine besylate, benzocaine/menthol, cyclophosphamide, cyclosporine (neoral), diphenhydramine INJ, fludarabine phosphate, filgrastim, furosemide, guaifenesin syrup, heparin (porcine) INJ, hydralazine HCl, hydroxyzine HCl, insulin aspart sliding scale INJ, levofloxacin INJ, loperamide HCl, lorazepam, methylprednisolone INJ, metoprolol succinate ext release 24hr, mycophenolate mofetil, ondansetron HCl, oxycodone HCl, oxymetazoline HCl, pantoprazole oral, piperacillin/tazobactam, posaconazole, prochlorperazine edisylate, prochlorperazine maleate, senna, simethicone, sildenafil, sulfamethoxazole/trimethoprim, vancomycin HCl, voriconazole, zolpidem tartrate, immune globulin |
| **PT_13** | Acetaminophen, Aciclovir, Amphotericin B Liposome INJ (Ambisome), Atovaquone, Benzocaine/Menthol, Cevimeline (saliva stimulant), Diphenhydramine INJ, Hydromorphone, Letermovir Oral, Lorazepam, Loperamide, Ondansetron oral and INJ, Oxycodone, Pantoprazole, Phenazopyridine, Piperacillin/Tazobactam INJ, Posaconazole Delayed Release, Scopolamine, Simethicone, Sucralfate, Sulfamethoxazole-trimethoprim, Ursodiol, Vancomycin |
| **PT_14** | Tacrolimus, Losartan, Budesonide, Pantoprazole, Acyclovir, Pregabalin, Fioricet, Timolol ophthalmic, Cholecalciferol, Calcium supplement, Cannabidiol, Omalizumab, Prednisone, Turmeric, Curcumin, Letermovir, Sulfamethoxazole-trimethoprim, Methotrexate, Diphenhydramine, Acetaminophen, Midostaurin, Cytarabine, Melphalan, Fludarabine |
| **PT_15** | Acetaminophen, Acyclovir, Amlodipine besylate, Chlorpromazine HCl, Clofarabine, Cytarabine INJ, Dasatinib Oral, Diphenhydramine oral and INJ, Labetalol, Lidocaine, Lorazepam, Loperamide, Lomotil, Melphalan, Octreotide Acetate, Omeprazole, Ondansetron HCl, Oxycodone, Posaconazole, Probenecid, Prochlorperazine Edisylate, Ranitidine HCl, Scopolamine, Thiotepa, immune globulin |
| **PT_16** | Acetaminophen, Acyclovir oral, Amlodipine Besylate, Budesonide 3mg oral capsule extended release, Carvedilol, Cyclosporine (neoral) oral and IVPB, Diphenhydramine INJ, Famotidine INJ, Fluconazole INJ, Furosemide INJ, Hydralazine INJ, Hydrocortisone INJ, Insulin Sliding Scale Aspart (rapid acting) INJ, Isavuconazonium, Labetalol, Lorazepam (C-IV) oral and INJ, Loperamide, Melatonin, Methylprednisolone INJ, Metoprolol, Mycophenolate Mofetil, Olanzapine, Ondansetron INJ, Omeprazole, Oxycodone, Pantoprazole INJ, Palonosetron INJ, Pentamidine, Prednisone, Prochlorperazine INJ, Ribavirin, Sulfamethoxazole/trimethoprim, Tamsulosin, Vancomycin INJ, Voriconazole INJ and oral, immune globulin |
| **PT_17** | Acetaminophen, Acetylcysteine, Acyclovir oral and INJ, Atovaquone, Baclofen, Benzonatate, Bisacodyl, Brincidofovir, CAB Fentanyl IV, Cefepime INJ, Chlorpromazine INJ, Diphenhydramine INJ, Enoxaparin INJ, Entecavir, Fentanyl INJ (C-II), Furosemide INJ, Guaifenesin syrup, Haloperidol INJ, Heparin IVCl, Hydralazine INJ, Hydroxyzine INJ, Lactulose, Lidocaine, Linezolid INJ, Loperamide, Lorazepam INJ, Metronidazole, Morphine sulfate INJ (C-II), Nystatin, Olanzapine, Oxycodone (C-II), Pantoprazole IV and oral, Piperacillin/Tazobactam INJ, Polyethylene Glycol 3350, Polyethylene Glycol Electrolyte, Senna, Simethicone, Sucralfate, Ursodiol, Valganciclovir, Vancomycin INJ and oral, Voriconazole oral and INJ, Zolpidem |
| **PT_18** | Acetaminophen, Aciclovir INF and oral, Amlodipine Besylate, Bebtelovimab, Ceftriaxone INJ, Ciprofloxacin, Diphenhydramine, Elthrombopag, Fluconazole, Furosemide INJ, Hydrocortisone INJ, letermovir INJ and oral, Levofloxacin oral, Lorazepam INJ (C-IV), Medroxyprogesterone oral, Methotrexate INJ, Olanzapine, Ondansetron INJ, Penicillin V Potassium, Pentamidine INJ, Phenazopyridine, Piperacillin/Tazobactam INJ, Rituximab INJ, Simethicone, Sulfamethoxazole/Trimethoprim, Tacrolimus (oral and IVCl), Ursodiol, Estradiol transdermal |
| **PT_19** | Acetaminophen, acyclovir, aprepitant, baclofen, budesonide, CAB hydromorphone IV, candesartan, cefepime INJ, cidofovir inj, ciprofloxacin INJ, cyclophosphamide INJ, diltiazem immediate release, diphenhydramine INJ, diphenoxylate/atropine (C-V) oral, furosemide, hydromorphone INJ (C-II), hydroxyzine HCl, labetalol, Letermovir INJ and oral, loperamide, lorazepam (C-IV), mesna, methylprednisolone INJ, metoprolol succinate ext release 24hr and INJ, morphine sulfate INJ and immediate release, mycophenolate mofetil oral and INJ, ondansetron, oxycodone (C-II), oxycontin (oxycodone extended release), palonosetron INJ, pantoprazole INJ and oral, pentamidine, phenazopyridine, piperacillin/tazobactam INJ, posaconazole delayed release, prednisone, pregabalin (C-V), prochlorperazine oral and INJ, ramelteon, simethicone, sirolimus, sucralfate, sulfamethoxazole/trimethoprim, ursodiol, verapamil INJ and IVCl, immune globulin |
| **PT_20** | Ergocalciferol, Opium tincture, Losartan, Diphenoxylate-atropine, Acyclovir, Niacinamide, Pantoprazole, Sulfamethoxazole-trimethoprim, Cyanocobalamin, Budesonide, Tacrolimus, Psyllium, Multivitamin, Calcium carbonate |

**Supplementary Table 2 |** Full list of medications received by post-HCT patients

| Pt | Extra-renal GVHD at the time of biopsy | Tubular injury (TI) | Tubular Atrophy (TA) | TMA | Glomeruli | Arterio- & Arteriolosclerosis | Inflammation | Other |
|---|---|---|---|---|---|---|---|---|
| **PT_1** | **Acute** | Diffuse acute TI with focal necrosis, severe | Focal TA with mild interstitial fibrosis | - | Minimal change nephrotic syndrome with massive proteinuria, global glom/sclerosis | Mild to moderate | Areas of active interstitial inflammation, mild to moderate | - |
| **PT_2** | **Acute, recurrent** | Diffuse, acute \TI with focal necrosis and reactive changes, severe | Focal TA with mild interstitial fibrosis | - | Mild focal glom. mesangial proliferation with polyclonal IgM deposits, global glom/sclerosis | Moderate | Active and chronic interstitial inflammation, mild to moderate | - |
| **PT_3** | **Acute, persistent/ recurrent** | Acute TI with early atrophy, moderate | | - | Ischemic glomerular capillary wrinkling with global glom/sclerosis | Moderate to severe | Mild interstitial inflammation | - |
| **PT_4** | **Acute** | | TA with interstitial fibrosis | - | Global glom/sclerosis | Mild with focal segmental arteriolar hyalinosis | Diffuse active and subacute interstitial inflammation, mild to moderate | - |
| **PT_5** | **Acute, persistent/ recurrent** | Diffuse acute TI, moderately severe | | - | Minimal glomerular changes | Mild to moderate | Mild, patchy interstitial inflammation | - |
| **PT_6** | **Acute, recurrent** | Mild-to-moderate diffuse acute TI | Focal TA with mild interstitial fibrosis | - | Diffuse podocytopathy suggestive of minimal change disease; focal partial ischemic collapse with global glomerulosclerosis | Moderately severe with luminal narrowing | Fairly diffuse active interstitial nephritis, moderate; frequent venulitis; focal tubulitis | Suspected renal GVHD post-tacrolimus taper |
| **PT_7** | **Overlap Syndrome** | Almost total tubular simplification, focally severe changes in medulla | Diffuse chronic ischemic damage, advanced/severe; mild to moderate, focally severe hypocellular interstitial fibrosis | - | Diffuse membranous glom/nephritis (Stage II-III, PLA2R-); ischemia/congestion, collapsing & segmental sclerosing | Severe with obliterative changes | Patchy moderate chronic inflammation; periglomerular & perivenular distribution | - |
| **PT_8** | **Chronic** | Lipid vacuolization | | - | Immune complex-secondary membranous-type glomerulonephritis, suggestive of glomerular GVHD, Global glom/sclerosis | Mild to moderate | Mild to moderate accumulation of circulating inflammatory cells | - |
| **PT_9** | **Chronic** | Foci of TI and atrophy with minimal interstitial fibrosis | - | Chronic endothelial injury | Global glom/sclerosis, diffuse podocytopathy consistent with minimal change nephrotic syndrome (clinical). | Moderate | Several capillaries marginated by neutrophils as well as mononuclear inflammatory cells | - |

**Supplementary Table 3 |** Summary of biopsy report findings for all patients (continued to next page)

| Pt | Extra-renal GVHD at the time of biopsy | Tubular injury (TI) | Tubular Atrophy (TA) | TMA | Glomeruli | Arterio- & Arteriolosclerosis | Inflammation | Other |
|---|---|---|---|---|---|---|---|---|
| **PT_10** | **Chronic** | | TA and interstitial fibrosis | Chronic glom. TMA changes, mild | Global ischemic collapse and global glom/sclerosis | Moderately severe with arteriolar hyalinosis | Mild chronic interstitial inflammation | - |
| **PT_11** | **Chronic** | Diffuse, acute TI, severe with focal necrosis | | Diffuse, acute and focal chronic TMA | Partial and global glom/sclerosis | Mid to moderate | Sparse infiltrates | - |
| **PT_12** | **Indeterm./ Intermittent/ Mild Chronic** | Diffuse acute and subacute TI with focal necrosis. | Foci of TA with minimal interstitial fibrosis | - | Mild glomerulomegaly with moderate podocytic injury, minimal change nephrotic syndrome, global ischemic collapse and global glom/sclerosis | Mild | Mild interstitial inflammatory infiltrate in the epithelium and peritubular capillaries | - |
| **PT_13** | **Indeterm.** | Focal evidence of TI | | - | Minimal glomerular changes | - | - | - |
| **PT_14** | **Indeterm.** | Prominent protein resorption droplets | Interstitial fibrosis and tubular atrophy, mild | - | Membranous glomerulonephritis, PLA2R-negative, no mesangial/subendothelial deposits, Congo red negative (no amyloid) Membranous | Moderate | mild | - |
| **PT_15** | **no** | | Diffuse TA and simplification, moderate. | Diffuse, chronic | Focal ischemic glomerular collapse and global glom/sclerosis | Severe with arteriolar intimal hyalinosis | Mild inflammatory infiltrate | Chronic endothelial injury |
| **PT_16** | **no** | Diffuse, acute TI and interstitial edema mild to moderate. | Focal TA with mild interstitial fibrosis | - | Global glom/sclerosis, glomerular endothelial changes suggestive of endotheliosis, consistent with minimal change nephrotic syndrome | Moderate with circumferential arteriolar intimal and focal medial hyalinosis | Sparse chronic lymphocytic infiltrate | - |
| **PT_17** | **no** | Diffuse acute TI with focal necrosis, severe | | - | Minimal glomerular changes | - | Mild interstitial edema | - |
| **PT_18** | **no** | Diffuse tubular simplification and acute TI | | Acute TMA primarily involving glomeruli | Acute TMA primarily involving glomeruli, Global ischemic collapse | Focal arteriolar intimal and medial hyalinosis | Mild | |
| **PT_19** | **no, resolved** | Patchy tubular simplification and tubular epithelial injury, mild. | Rare foci of TA. | - | Mild glomerulomegaly with minimal changes, global glom/sclerosis | Moderately severe with luminal narrowing | - | - |
| **PT_20** | **no, quiescent** | | Interstitial fibrosis and tubular atrophy, severe. | - | membranous glomerulonephritis | Moderate to severe | - | - |

**Supplementary Table 3 (continued) |** Summary of biopsy report findings for all patients

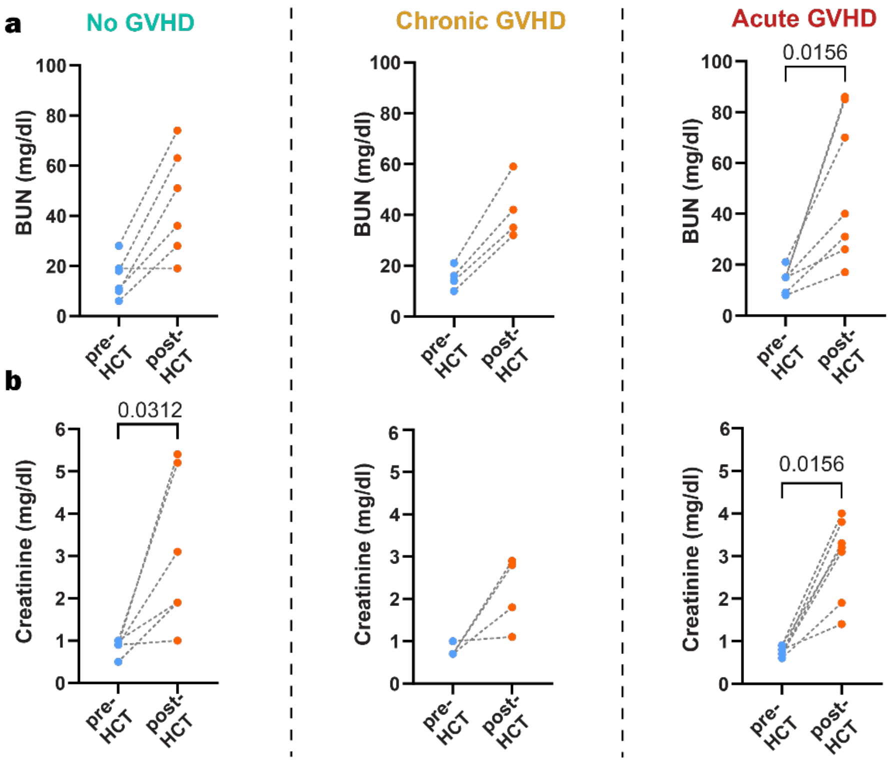


**Supplementary Fig. 1 | Renal function before and after HCT stratified by extra-renal GVHD status.** a, Blood urea nitrogen (BUN) levels measured before hematopoietic cell transplantation (pre-HCT) and after transplantation (post-HCT) in patients stratified by: no GVHD, chronic GVHD, or acute GVHD.
b, Corresponding serum creatinine measurements in the same patients.

| Code | Diagnosis |
|---|---|
| O1 | Acute kidney injury on chronic kidney disease (AKI on CKD), history of cutaneous squamous cell carcinoma |
| O2 | Metastatic ovarian germ cell tumor |
| O3 | Chronic kidney disease (CKD) |
| O4 | Gastric cancer |
| O5 | B-cell lymphoma with lymphomatous infiltration of the kidneys |
| O6 | Renal cell carcinoma |

**Supplementary Table 4 |** Clinical diagnoses of patients included in the “Other etiologies” cohort. These are patients who did not undergo HCT and presented with kidney injury of unrelated etiologies.

**Examples of Patient Glomerulosclerosis**

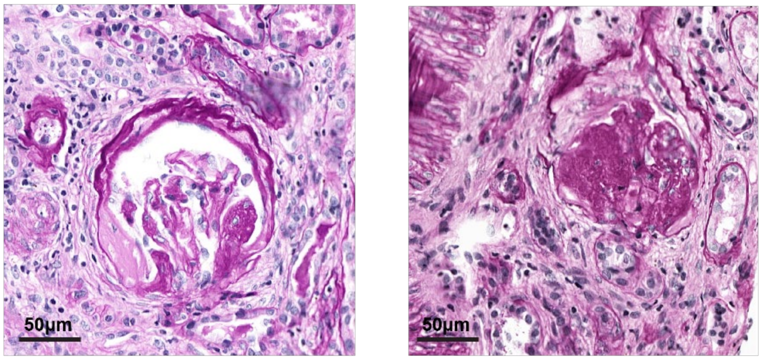


**Supplementary Fig. 2 | a** Representative PAS staining of tubular injury and glomerulosclerosis in post- HCT kidneys

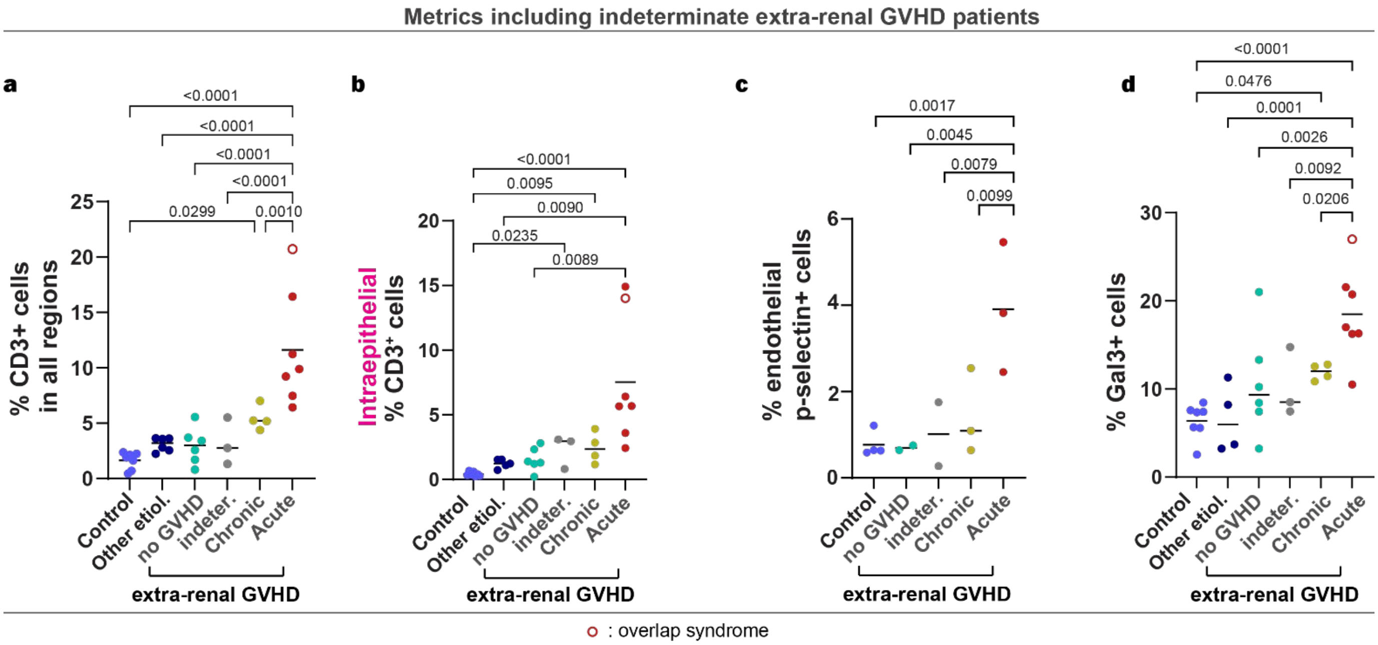


**Supplementary Fig. 3 | T cell infiltration and Galectin-3 expression in post-BMT patients, including the indeterminate group . a-b**, Percentage of CD3⁺ cells in all regions **(a)** and in epithelium **(b)** comparing control groups against post-HCT patients, with various extra-renal GVHD manifestations **c,** Quantification of galectin-3⁺ cells comparing control groups with various GVHD manifestations. **a-c** $n_{control}$=6, $n_{other\ etiologies}$ = 4-6, $n_{post\text{-}HCT}$=20, Ordinary one-way ANOVA with Tukey correction. Hollow points indicate patients with overlap syndrome. Line represents the mean.

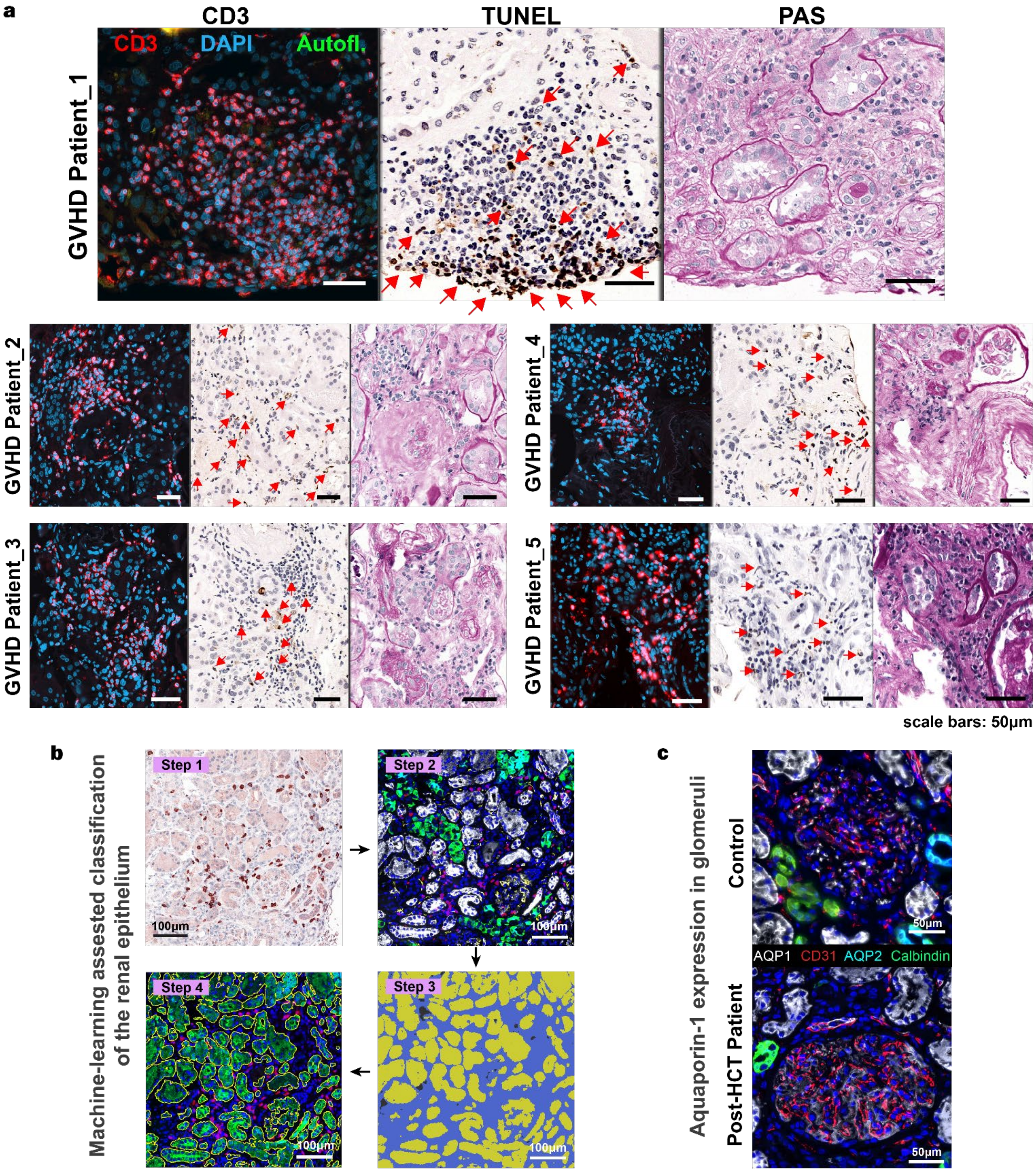


**Supplementary Fig. 4 | a,** Representative serial sections from post-HCT kidney biopsies showing regions of dense CD3+ T-cell infiltration (left), apoptosis assessed by TUNEL staining (center), and corresponding renal morphology by PAS (right). Apoptotic cells were observed in regions with prominent T-cell infiltration on adjacent serial sections. Scale bar, 50 µm. **b,** Workflow for T cell localization analysis (following the direction of the arrows): 1) IHC of $CD3^{+}$ cells, 2) Immunofluorescence markers for kidney compartments: calbindin (distal tubules), CD31 (endothelium), Aquaporin 1 (proximal tubules), and Aquaporin 2 (collecting ducts), illustrating $CD3^{+}$ localization (red, pseudo-color based on IHC), 3) Machine-learning predicted tissue identity (yellow= epithelium, blue=stroma, black= background) , 4) Epithelium annotations (yellow outlines) based on the ML-predictions **c,** Increased expression of Aquaporin-1 in glomeruli of Post-HCT patients

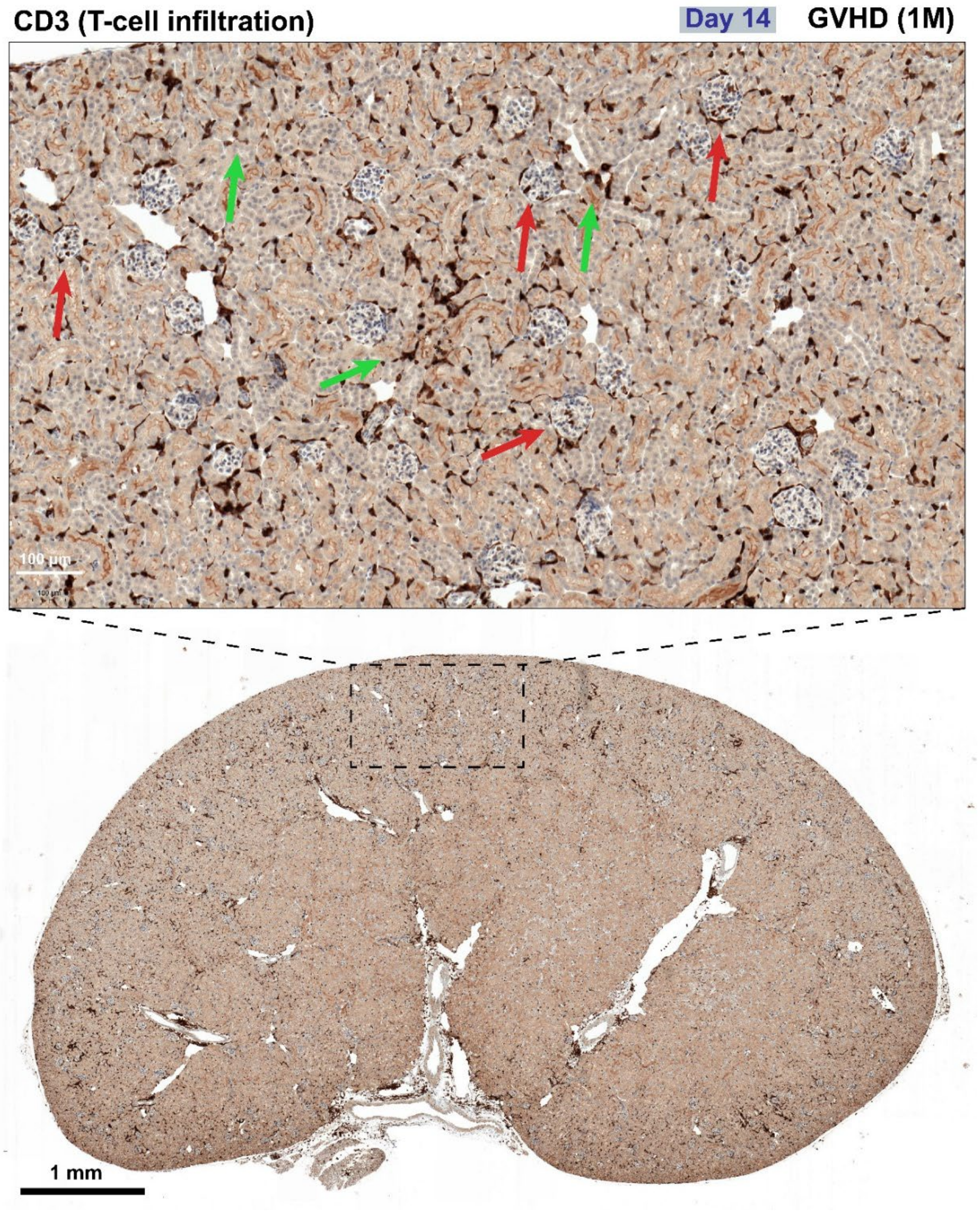


**Supplementary Fig. 5:** T cell infiltration (CD3) on day 14 post transplantation in the GVHD (1M) mouse model. High magnification of the cortex (top) and whole kidney slide (bottom). Red arrows indicate T cells infiltrating glomeruli, while green arrows indicate T cells infiltrating the tubules.

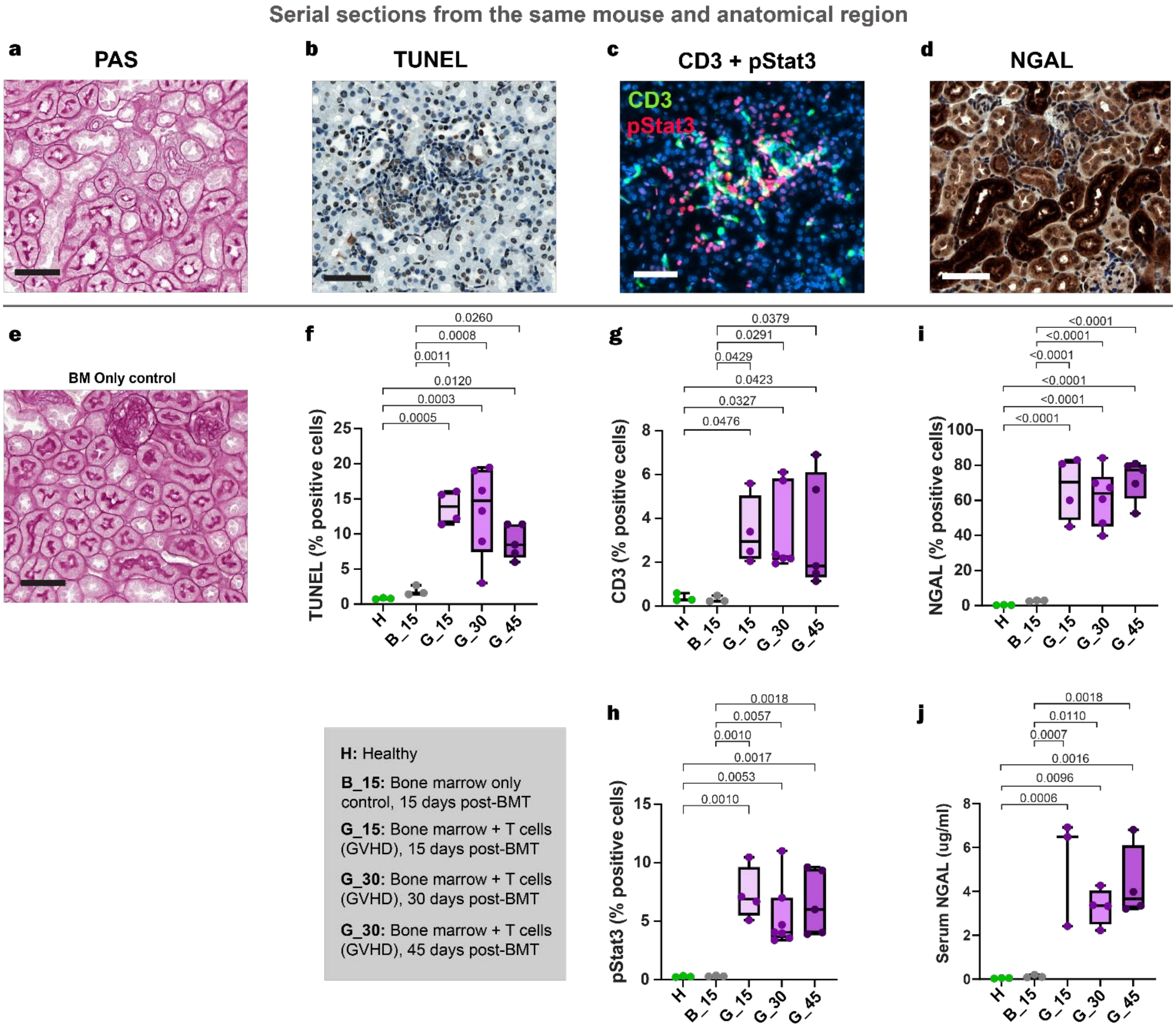


**Supplementary Fig. 6 | Minor GVHD murine model exhibits renal injury and inflammatory activation**. **a-d**, Representative images of renal morphology (PAS), apoptosis (TUNEL), tubular injury (NGAL), T cell infiltration and pSTAT3 signaling (CD3, pSTAT3), and renal morphology (PAS) in a GVHD mouse. Images are derived from serial sections of the same mouse and anatomical region. **e,** Representative PAS-stained kidney section from a bone marrow transplantation–only (BMO) control mouse. **a-e** Scale bar, 50 µm. **f-i**, Quantification of TUNEL$^+$ cells (f), CD3$^+$ cells (g), pSTAT3$^+$ cells (h), NGAL$^+$ cells (i). **j**, Serum NGAL levels measured by ELISA. The box extends from the 25th to 75th percentiles and the whiskers from min to max. The line represents the median. Statistical significance was determined using the tests indicated in the panels. H = healthy mice; $B_{15}$ = bone marrow transplantation only (day 15); $G_{15}$, $G_{30}$, $G_{45}$ = minor GVHD mice analyzed at days 15, 30, and 45 after transplantation.

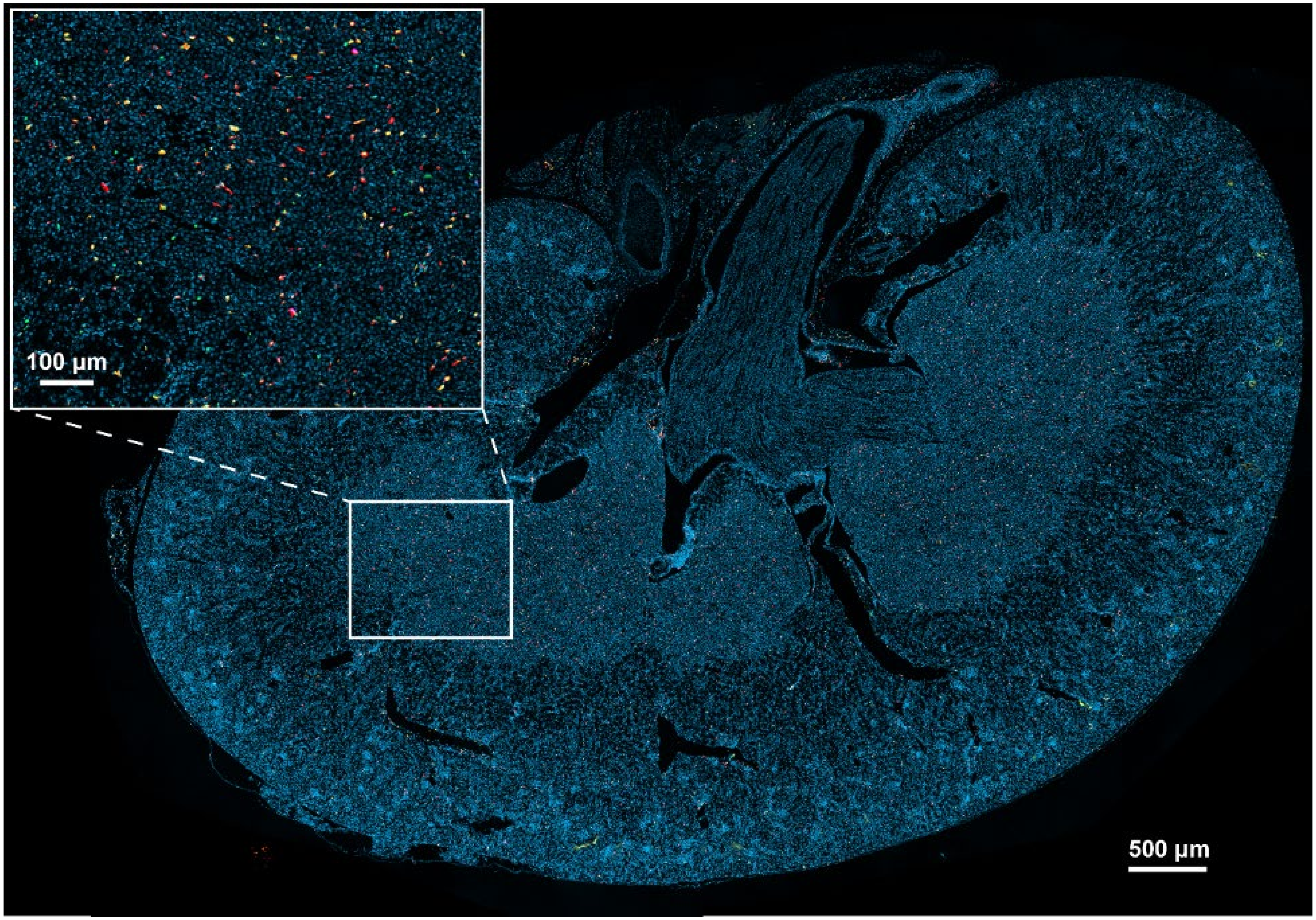


**Supplementary Fig. 7 | Whole kidney multiplex immunofluorescence:** Merged immunofluorescence image and higher magnification image (snippet) of kidney tissue stained for CD8 (red), Tbet (green), CD4 (orange), FoxP3 (magenta), CD19 (dark blue), Ly6G (yellow) and DAPI (cyan). Whole-tissue image analysis with a customized algorithm allows for extraction of quantitative and spatial information on the immune landscape of the kidney.

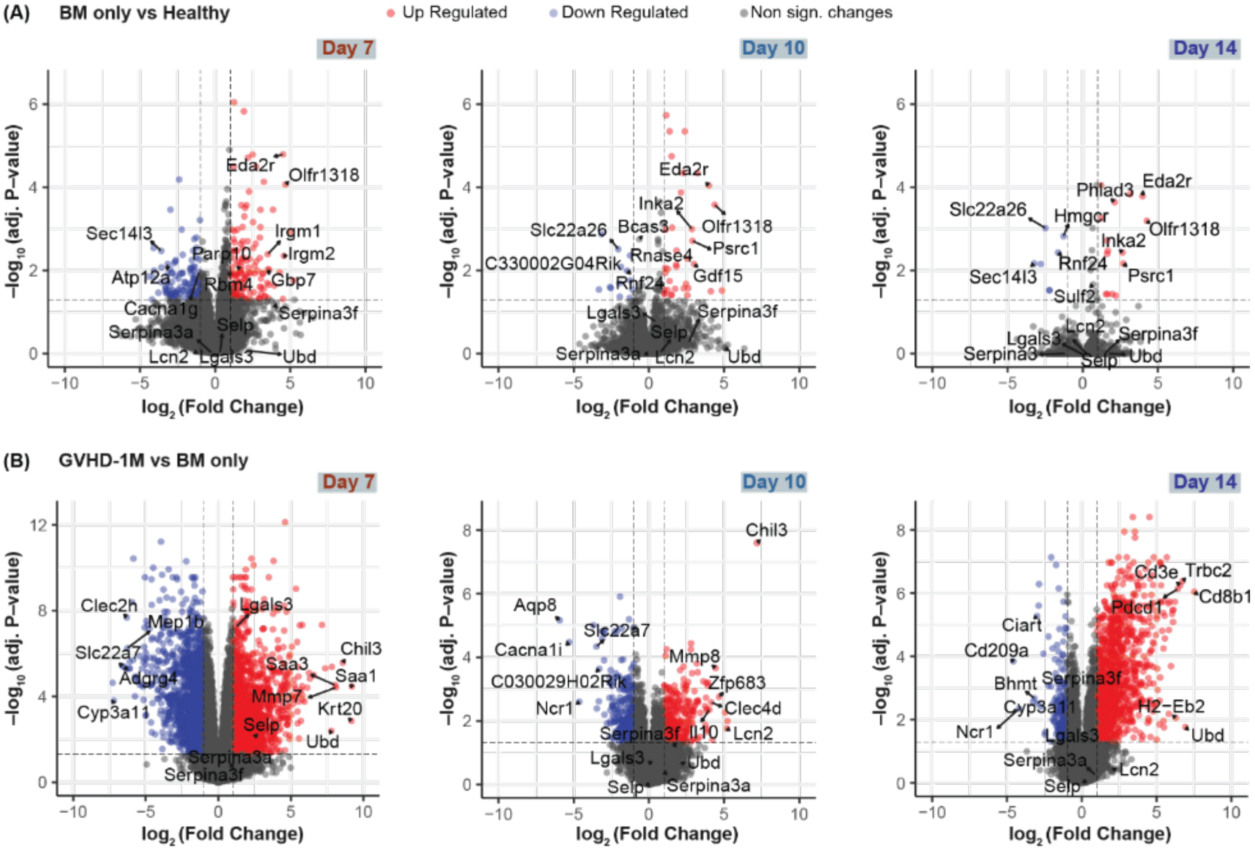


**Supplementary Fig. 8 | Transcriptome analysis of kidney lysates from BM only mice and GVHD (1M) mice.** Volcano plots of genes differentially expressed between (**a**) BM only mice and healthy mice and (**b**) GVHD mice and BM only mice on days 7, 10, and 14 post BMT.

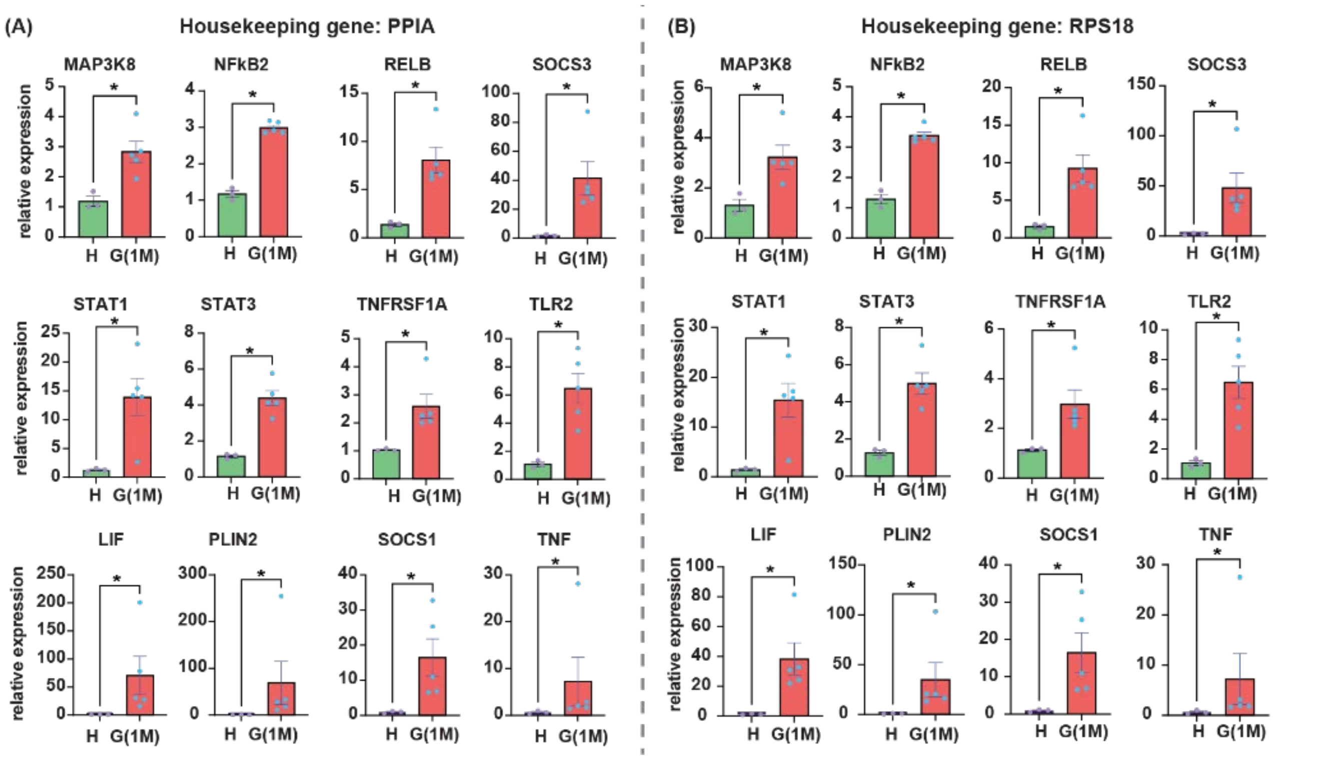


**Supplementary Fig. 9 | qPCR of top genes in upregulated pathways in kidney lysates from GVHD (1M) vs healthy mice. (a)** Housekeeping gene was PPIA **(b)** Housekeeping gene was RPS18 **(***p<0.05,**p<0.010,Mann-WhitneyUtest. Data represent mean ± s.d.)

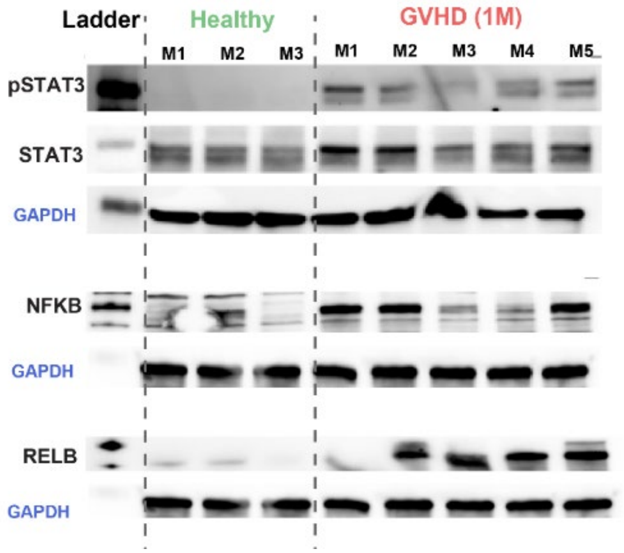


**Supplementary Fig. 10 | Protein expression in kidney lysates by Western Blot in healthy and GVHD (1M) mice.** Phospho-Stat3, NFκB and RelB are overexpressed in mice with GVHD. $n_{healthy}$= 3, $n_{GVHD}$=5.

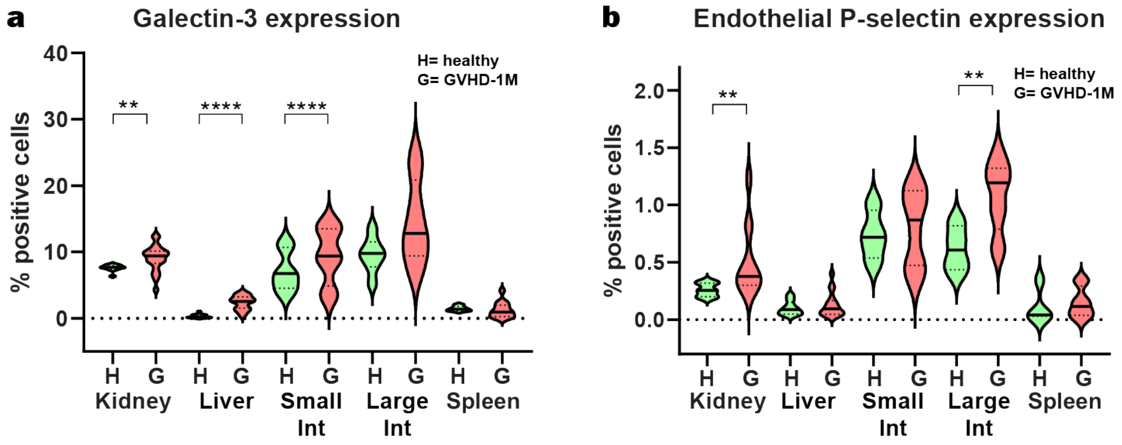


**Supplementary Fig. 11 | Murine Galectin-3 (A) and endothelial P-selectin (B) expression in all tissues in healthy and GVHD (1M) mice quantified by immunofluorescence.** P-selectin was quantified only when it co-localizes with CD31, to exclude expression in blood cells. H = healthy. **(***p<0.05,**p<0.01, Mann-Whitney U test.)

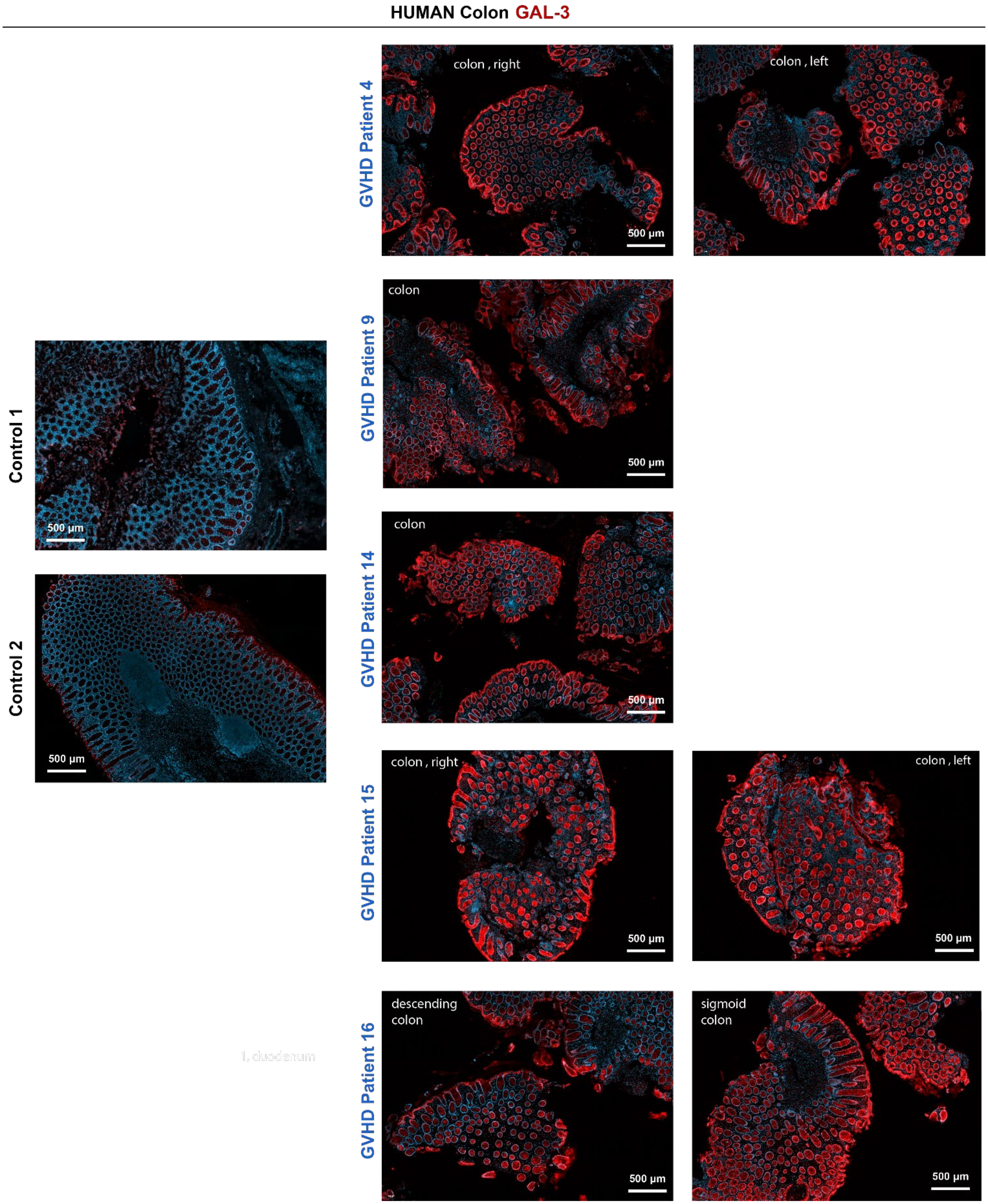


**Supplementary Fig. 12 | Galectin-3 expression in human GVHD target organs.** Representative immunofluorescence images of galectin-3 (Gal-3) in human colon tissue. Left, commercially obtained healthy control colon; right, colon biopsies from patients with GVHD within the kidney biopsy cohort. Images demonstrate increased Gal-3 staining in GVHD tissue compared to control. Scale bar: 500 µm.

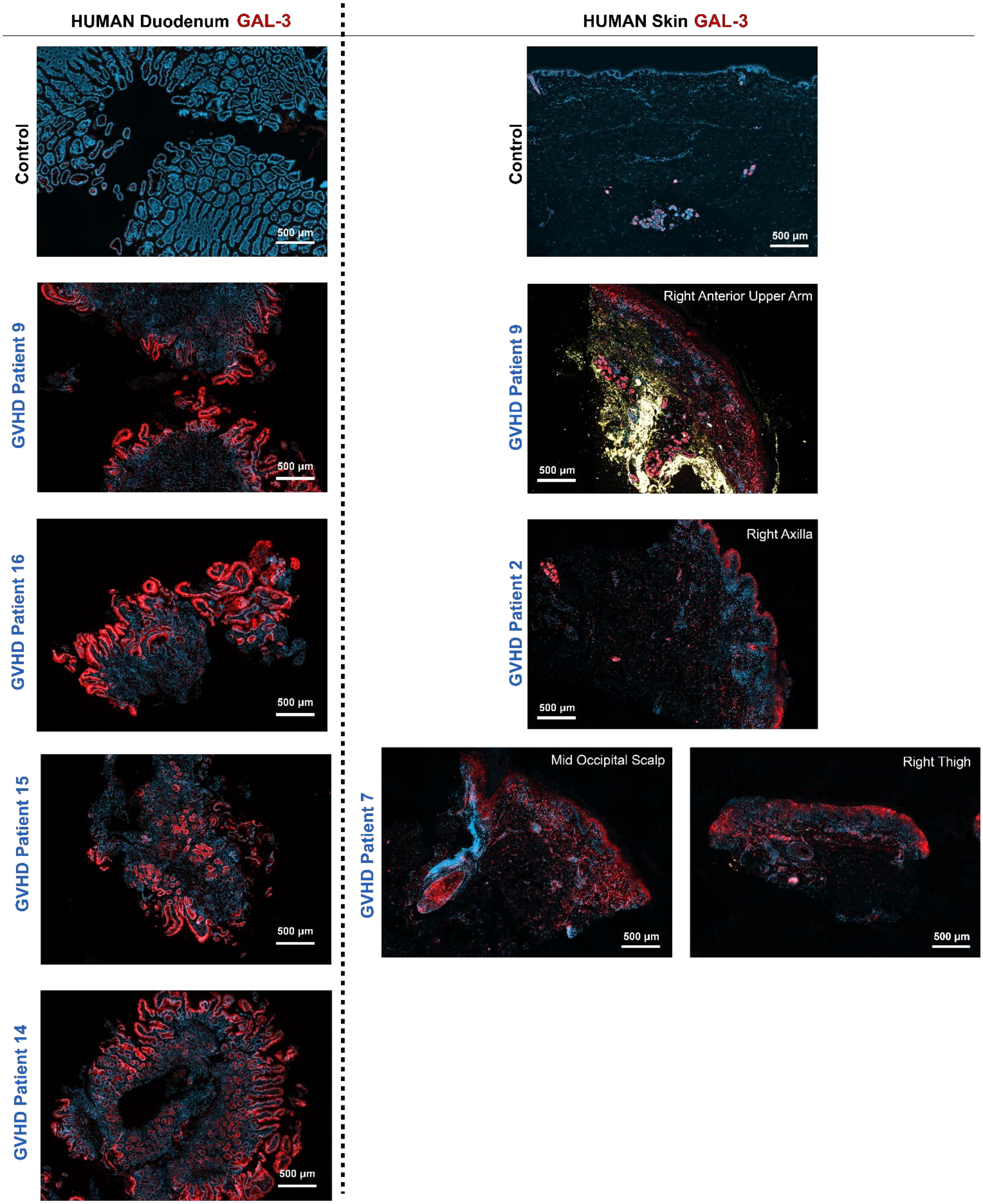


**Supplementary Fig. 13 | Galectin-3 expression in human GVHD target organs.** Representative immunofluorescence images of galectin-3 (Gal-3) in duodenum; left and skin; right. Top row: commercially obtained healthy controls, below, biopsies from patients with GVHD within the kidney biopsy cohort. Images demonstrate increased Gal-3 staining in GVHD tissue compared to control. Scale bar: 500 µm.

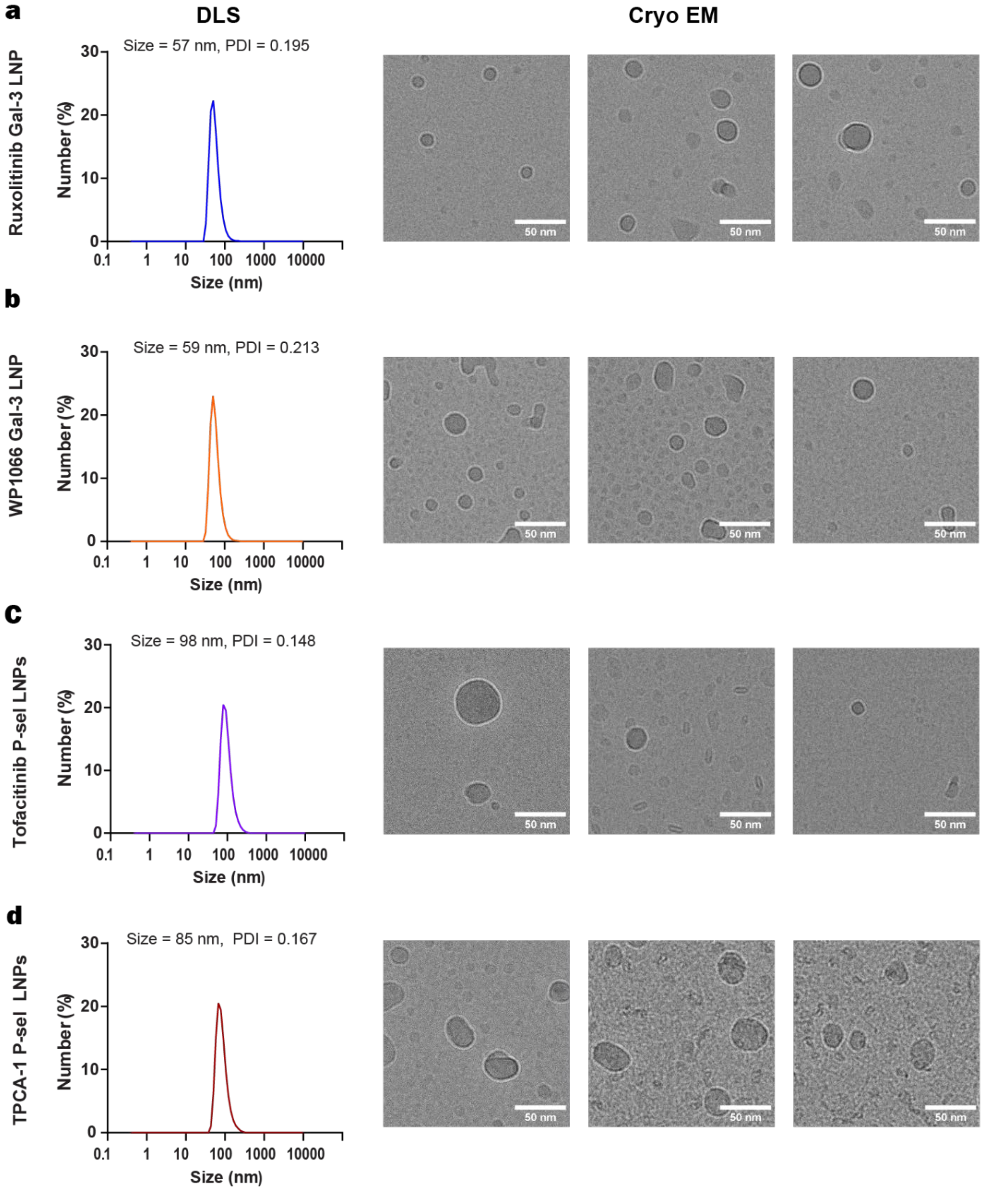


**Supplementary Fig. 14 | LNP size and structure for different drugs:** Hydrodynamic diameter measurements **(left)** and Cryo EM pictures (**right**) of Ruxolitinib Gal-3 LNPs, WP1066 Gal-3 LNPs, Tofacitinib P-sel LNPs, TPCA-1 P-sel LNPs. (Scale bar: 50nm)

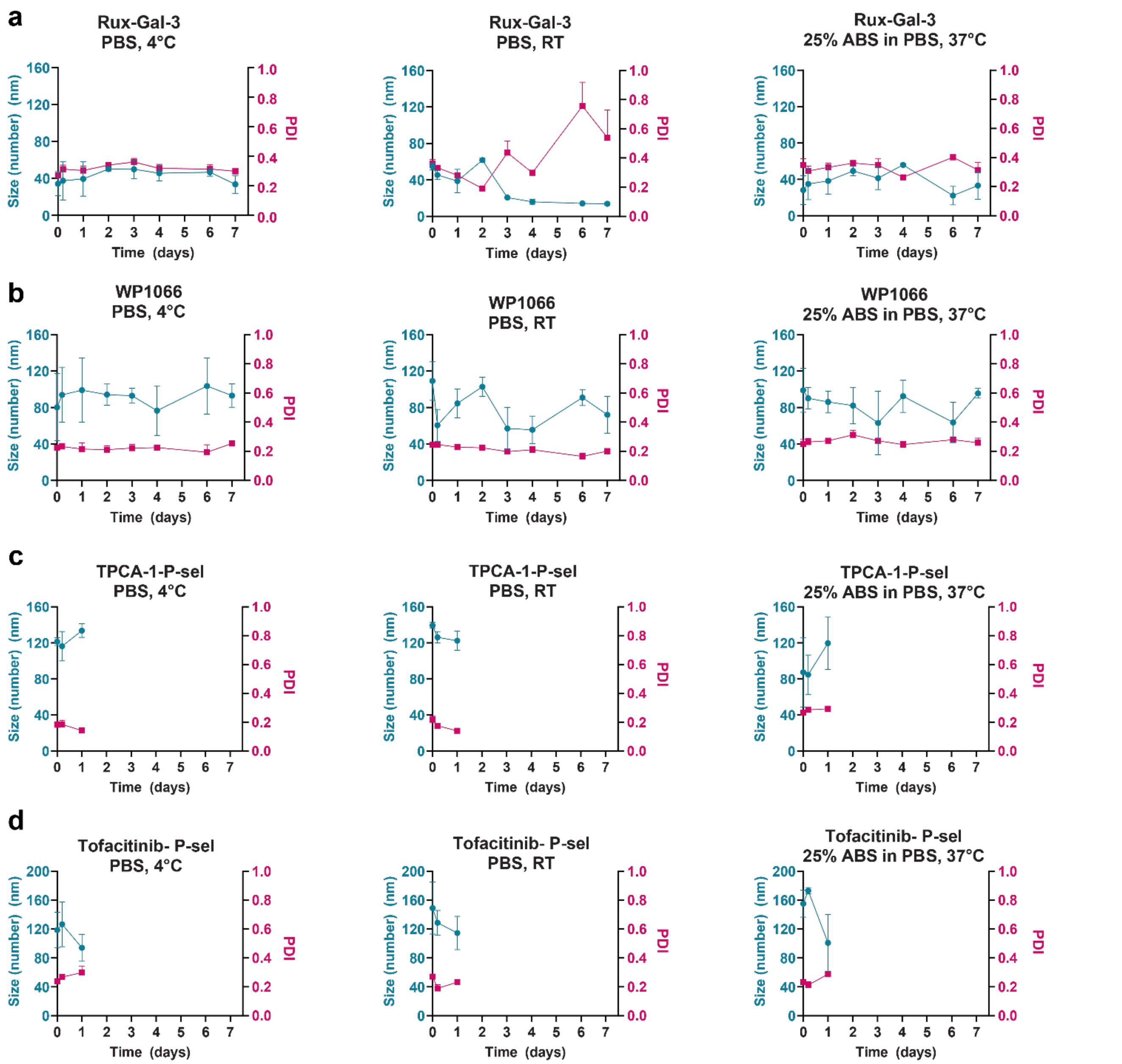


**Supplementary Fig. 15 | Time-dependent stability of Gal-3-targeted LNP drug formulations under storage and blood-mimicking conditions.** Hydrodynamic diameter (blue) and polydispersity index (PDI, pink) of Ruxolitinib-Gal-3 (**a**), WP1066 (**b**), TPCA-1 (**c**), and Tofacitinib LNPs (**d**), were quantified by DLS over 7 days under PBS at 4 °C, PBS at room temperature, and 25% adult bovine serum (ABS) in PBS at 37 °C. Ruxolitinib-Gal-3 and WP1066 LNPs maintained the most stable size and lowest PDI values across conditions, while TPCA-1 and Tofacitinib LNPs showed reduced stability particularly at RT and in serum.

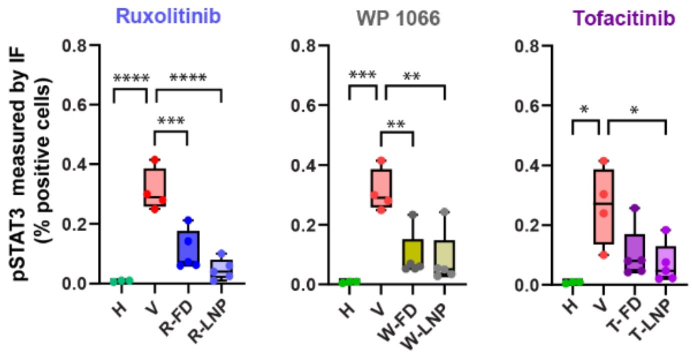


**Supplementary Fig. 16 | The effect of ruxolitinib, WP1066 or tofacitinib treatment (free drug or LNP) on phospho-STAT3 levels in GVHD (1M), measured by immunofluorescence**

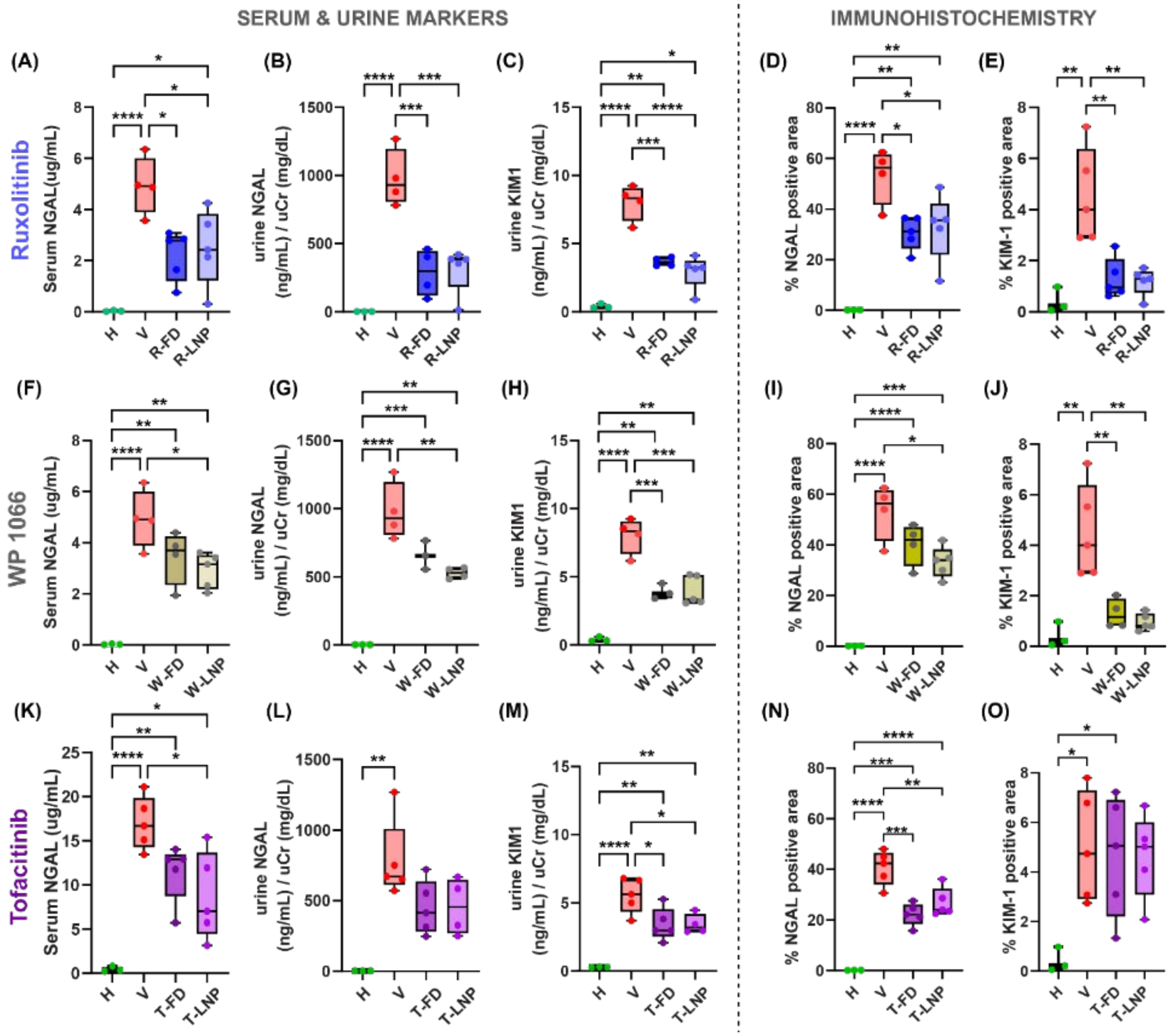


**Supplementary Fig. 17 | The effect of ruxolitinib, WP1066 or tofacitinib treatment (free drug or LNP) on kidney injury markers in GVHD (1M).** The effects of Ruxolitinib (**A-E**), WP1066 (**F-J**) or tofacitinib (**K-O**) on serum NGAL, urine NGAL, urine KIM1, and histological levels of NGAL and KIM1. Serum and urine markers were measured by Elisa, while histological levels were measured by IHC. R-FD: Ruxolitinib Free Drug, R-LNP: Ruxolitinib LNP, W-FD: WP1066 Free Drug, W-LNP: WP1066 LNP, T-FD: Tofacitinib Free Drug, T-LNP: Tofacitinib LNP. n=3-5 mice per group. **(***p<0.05, **p<0.01,***p<0.001, ****p<0.0001, Ordinary one-way ANOVA.)

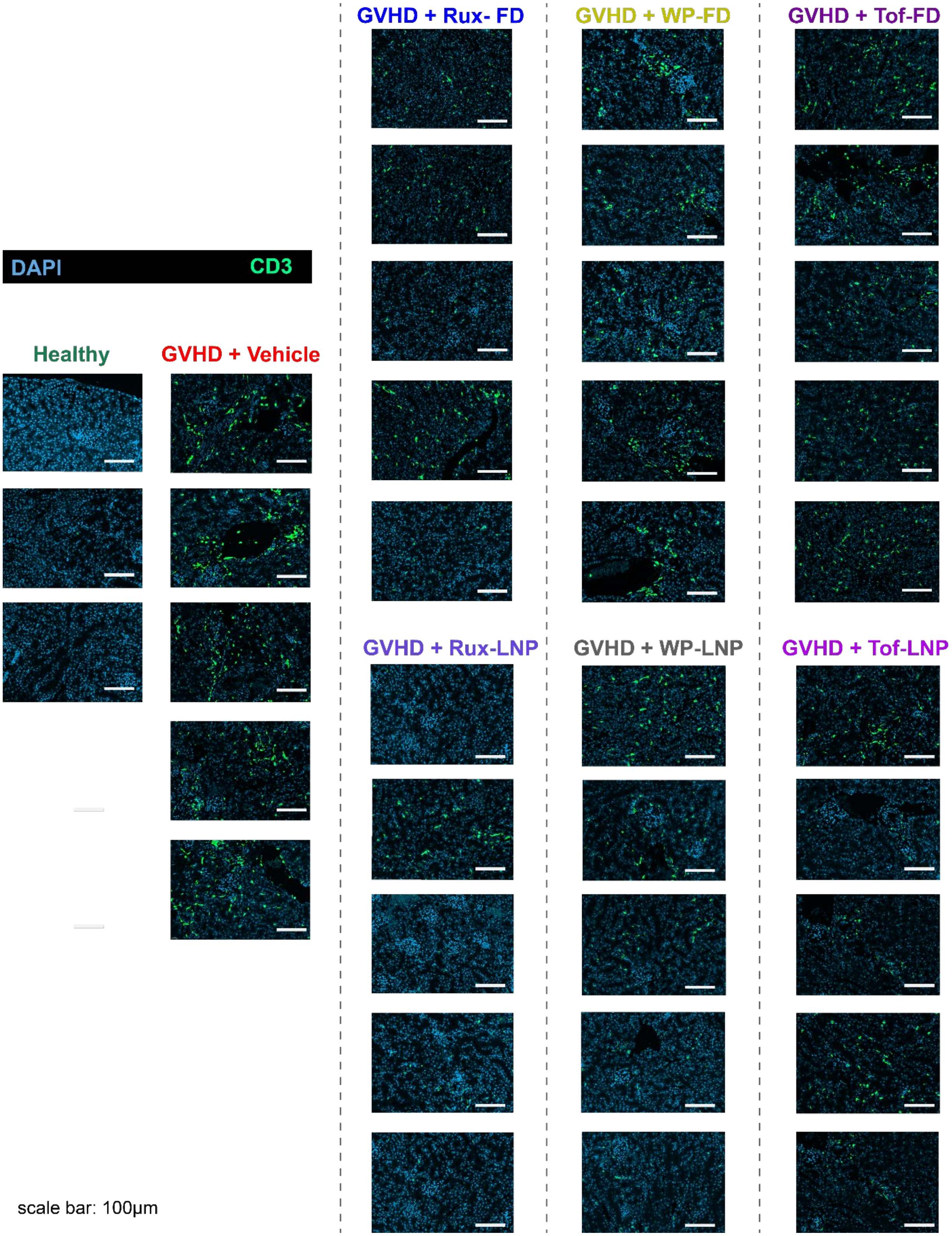


**Supplementary Fig. 18 | CD3 staining across treatment groups in the major mismatch GVHD (1M) model.** Representative immunofluorescence images of renal tissue showing CD3⁺ T cell infiltration in healthy controls and GVHD mice treated with vehicle (V), ruxolitinib (Rux), WP-1066 (WP), or tofacitinib (Tof), administered as free drug (FD) or lipid nanoparticle (LNP) formulations. Scale bar, 100 μm

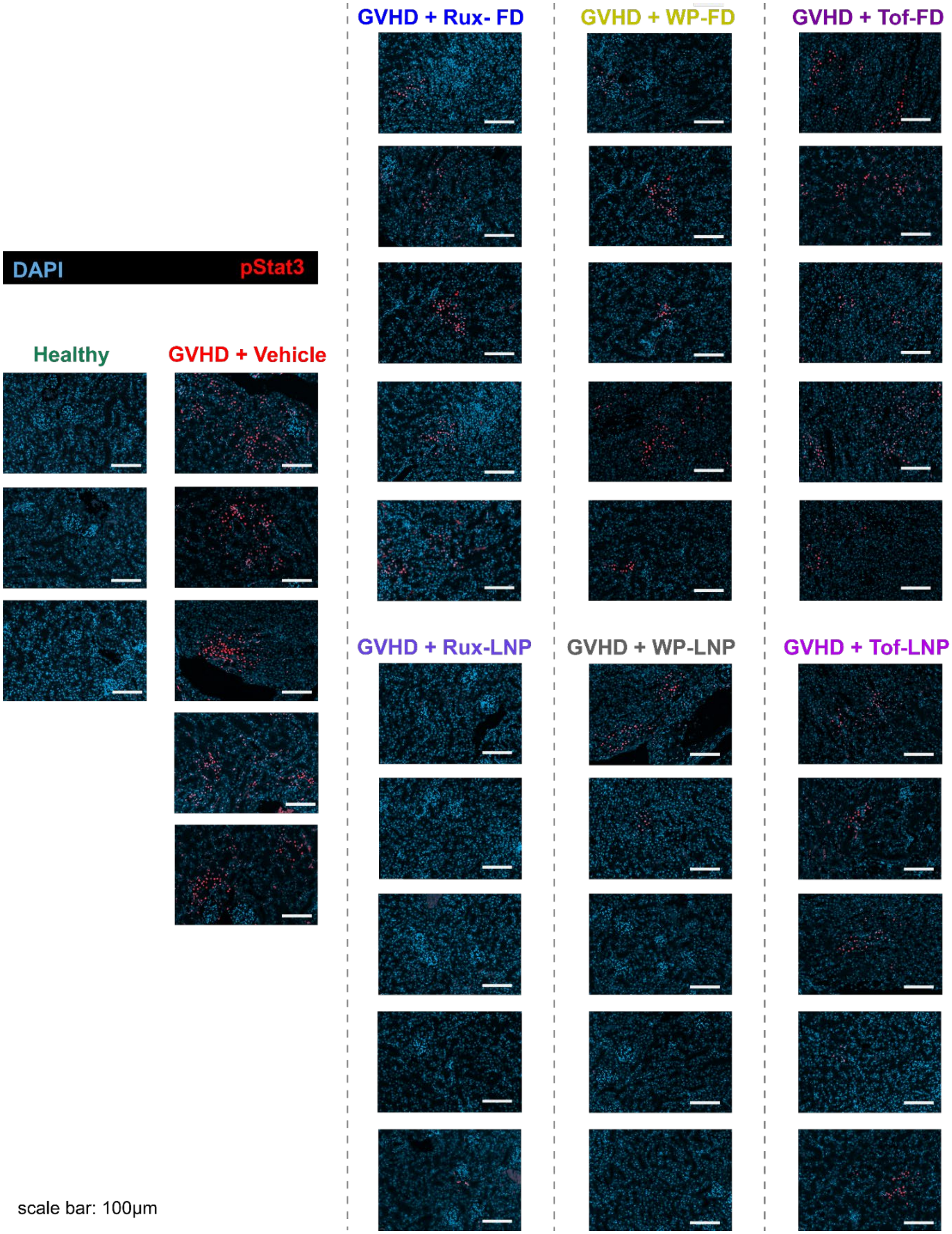


**Supplementary Fig. 19 | pSTAT3 staining across treatment groups in the major mismatch GVHD (1M) model**. Representative immunofluorescence images of renal tissue showing phosphorylated STAT3 (pSTAT3) in healthy controls and GVHD mice treated with vehicle (V), ruxolitinib (Rux), WP-1066 (WP), or tofacitinib (Tof), administered as free drug (FD) or lipid nanoparticle (LNP) formulations. .Scale bar, 100 µm

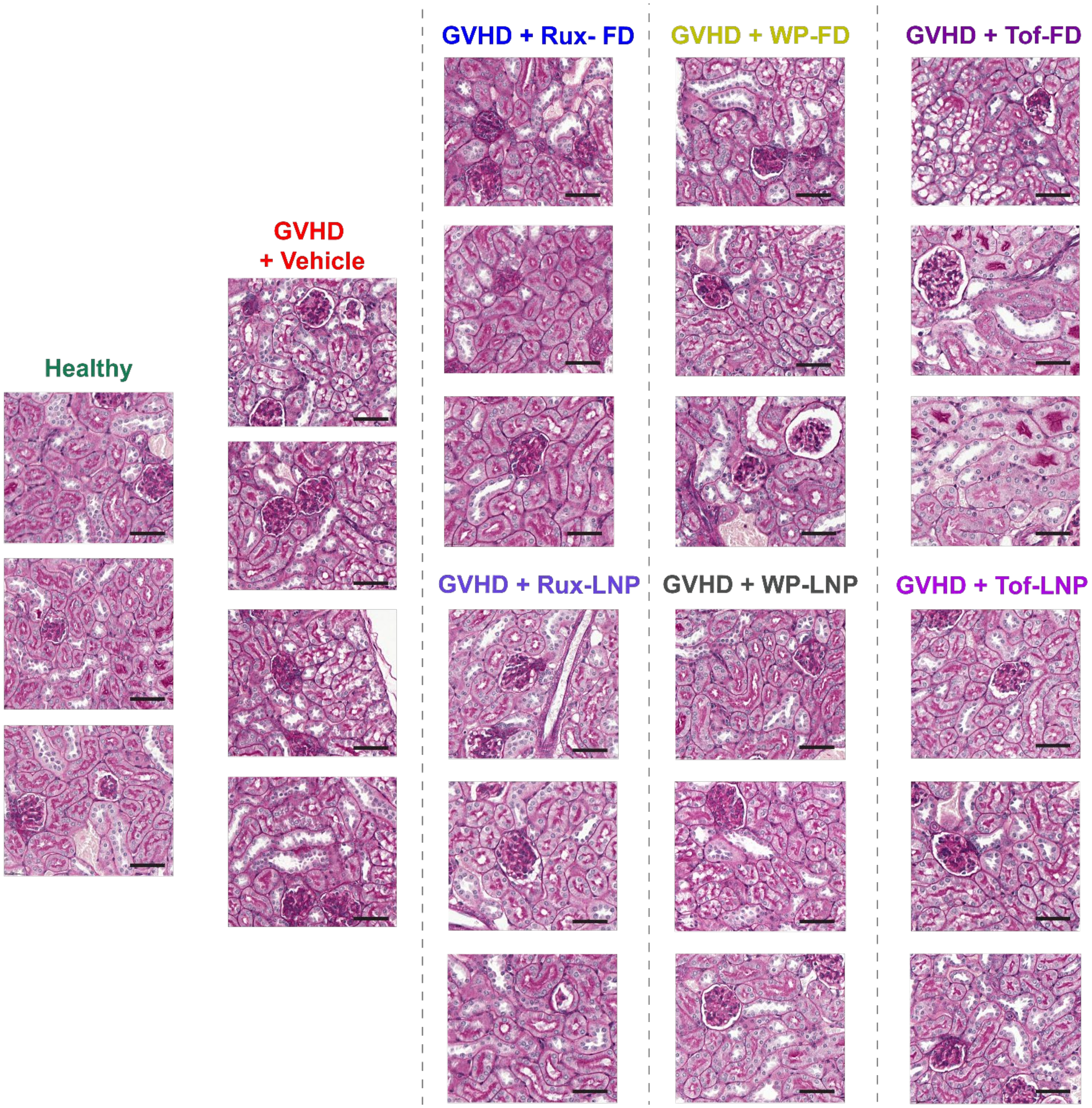


**Supplementary Fig. 20 | PAS staining of renal tissue across treatment groups in the major mismatch GVHD (1M) model.** Representative periodic acid–Schiff (PAS)–stained kidney sections from healthy controls and GVHD mice treated with vehicle (V), ruxolitinib (Rux), WP-1066 (WP), or tofacitinib (Tof), administered as free drug (FD) or lipid nanoparticle (LNP) formulations. Images illustrate differences in tubular injury, including epithelial damage, tubular dilation, loss of brush border, as well as vacuolization and intratubular cellular debris across treatment conditions. Scale bar, 50 μm.

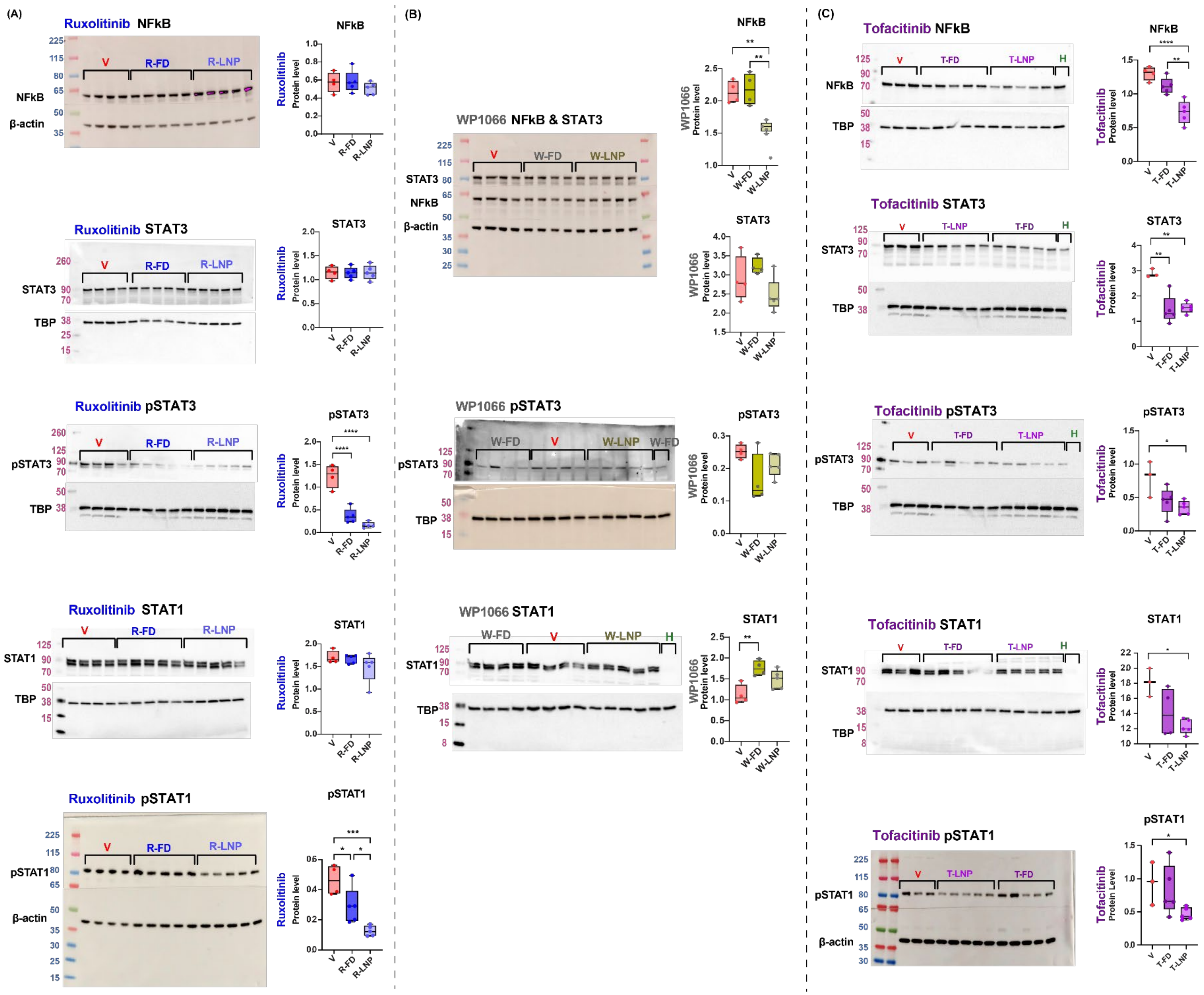
(A)
Ruxolitinib NFkB
NFkB
β-actin
V
R-FD
R-LNP
Ruxolitinib STAT3
STAT3
TBP
Ruxolitinib pSTAT3
pSTAT3
Ruxolitinib STAT1
STAT1
Ruxolitinib pSTAT1
pSTAT1
Ruxolitinib Protein level
(B)
WP1066 NFkB & STAT3
W-FD
W-LNP
WP1066 pSTAT3
WP1066 STAT1
H
WP1066 Protein level
(C)
Tofacitinib NFkB
T-FD
T-LNP
Tofacitinib STAT3
Tofacitinib pSTAT3
Tofacitinib STAT1
Tofacitinib pSTAT1
Tofacitinib Protein level
Tofacitinib Protein Level

**Supplementary Fig. 21 | The effect of free or LNP-encapsulated Ruxolitinib, WP1066, and Tofacitinib on the levels of top overexpressed proteins in GVHD (1M), measured by western blot**. Representative western blot bands and quantifications for the effect of (A) Ruxolitinib, (B) WP1066, and (C) Tofacitinib on STAT3, pSTAT3, NF-κB, STAT1, and pSTAT1 levels. R-FD: Ruxolitinib Free Drug, R-LNP: Ruxolitinib LNP, W-FD: WP1066 Free Drug, W-LNP: WP1066 LNP, T-FD: Tofacitinib Free Drug, T-LNP: Tofacitinib LNP. n=3-5 mice per group. (*p<0.05, **p<0.01,***p<0.001, ****p<0.0001, Ordinary one-way ANOVA.)

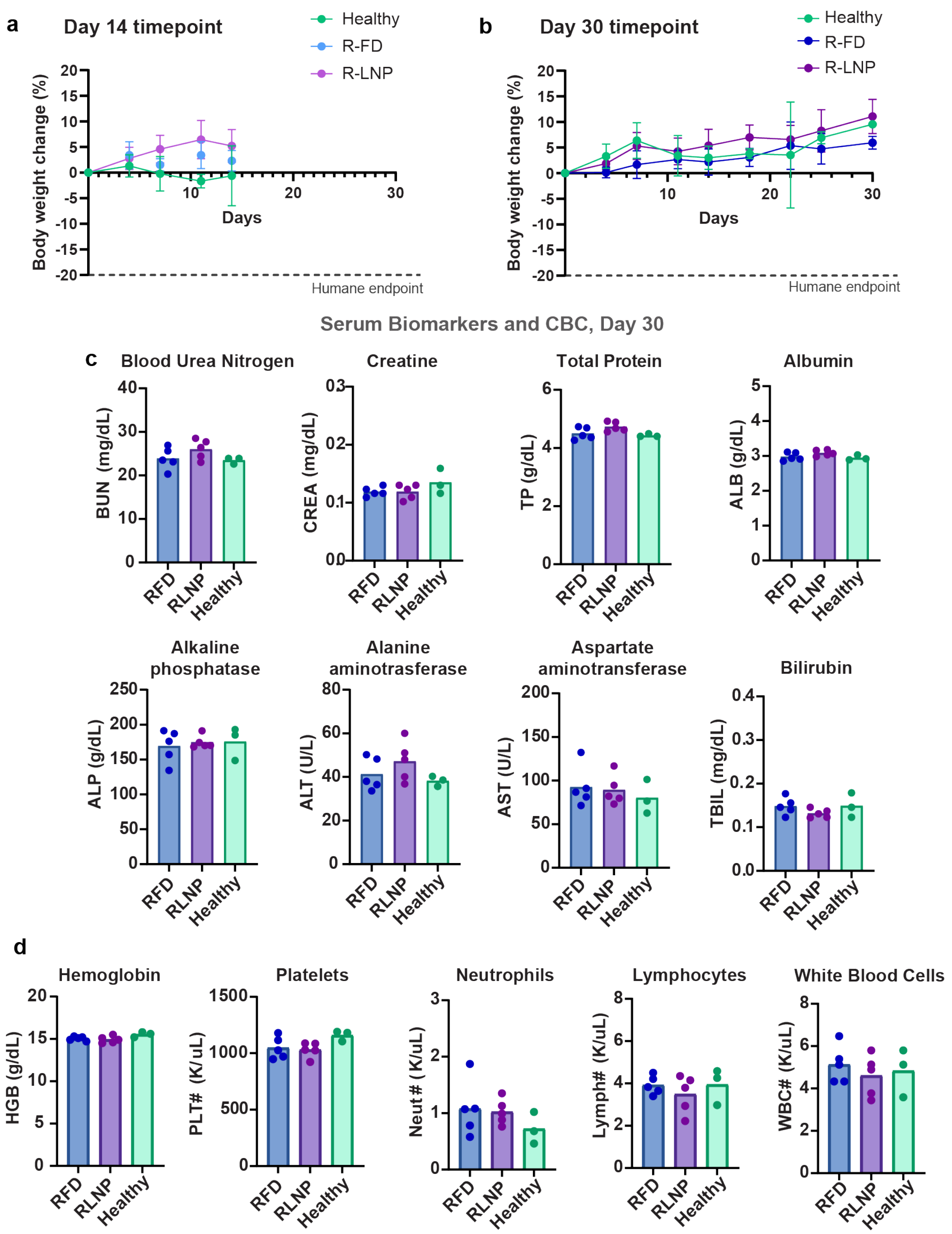


**Supplementary Fig. 22 | Long-term systemic toxicity assessment of free ruxolitinib and ruxolitinib-loaded lipid nanoparticles**. Percentage change in body weight of healthy BALB/c mice over (a) 14 or (b) 30 days following intraperitoneal injections of free ruxolitinib (R-FD), galectin-3–targeted ruxolitinib lipid nanoparticles (R-LNP) every other day, or untreated healthy controls (H). The dashed line indicates the humane endpoint (20% weight loss). (c) Serum chemistry analysis after 30 days of treatment, including blood urea nitrogen (BUN), creatinine, total protein, albumin, alkaline phosphatase (ALP), alanine aminotransferase (ALT), aspartate aminotransferase (AST), and total bilirubin, remained within physiological ranges across all groups. (d) Hematological parameters at day 30, including hemoglobin, platelet counts, neutrophils, lymphocytes, and total white blood cells, showed no evidence of hematologic toxicity associated with R-LNP treatment. Data are presented as mean ± SD (**a,b**) or individual animals with mean ± SD (**c,d**); $n_{Healthy}$=3, $n_{RFD}$=5, $n_{LNP}$=5 in all panels. Statistical analysis was performed using one-way ANOVA, with no statistically significant differences detected between groups.

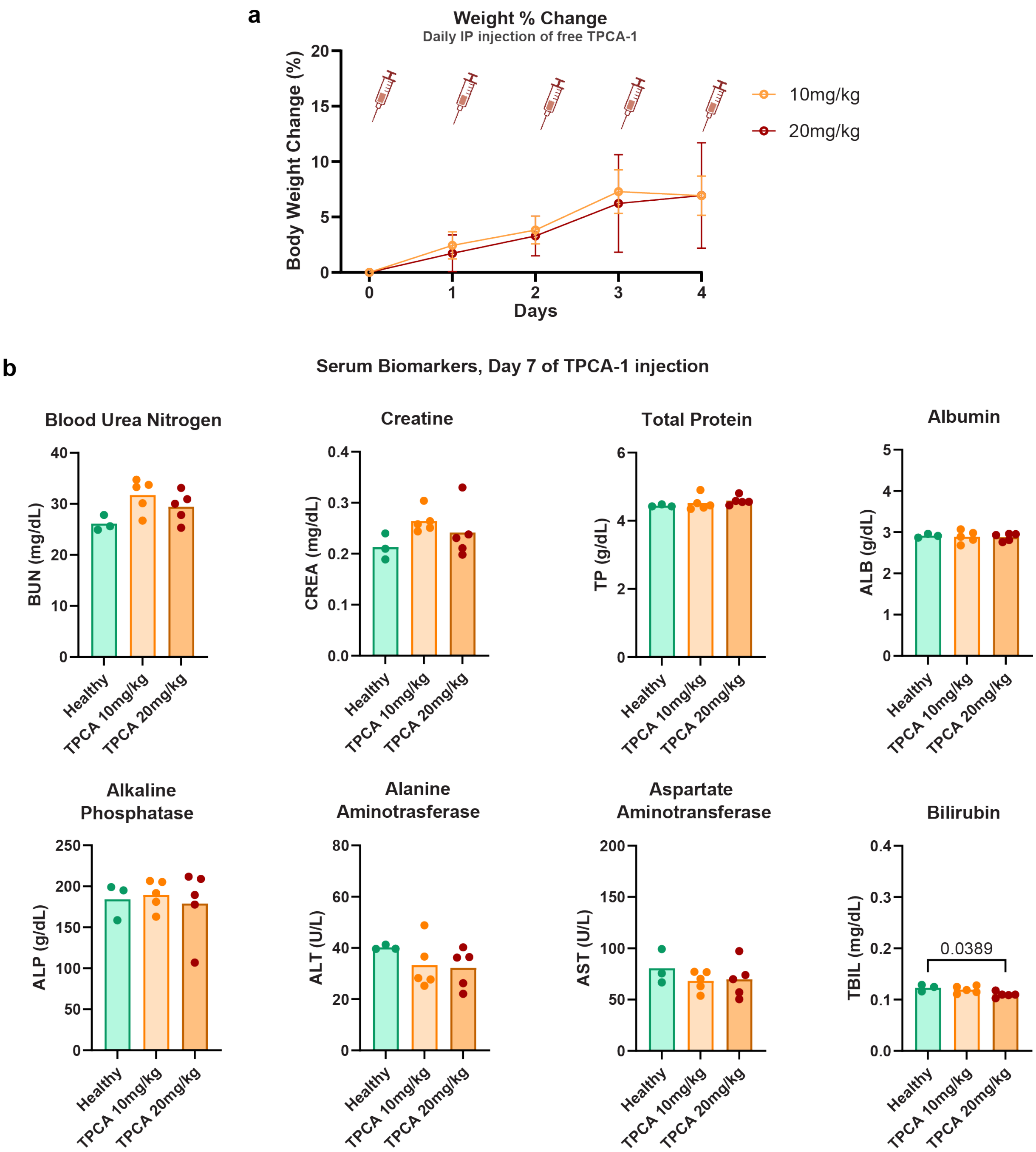


**Supplementary Fig. 23 | Systemic toxicity assessment of TPCA-1 in healthy mice. a**, Body weight change over time in healthy BALB/c mice following daily intraperitoneal administration of TPCA-1 at 10 mg/kg or 20 mg/kg. No weight loss was observed across treatment groups over the dosing period. **b,** Serum chemistry analysis on day 7, including blood urea nitrogen (BUN), creatinine (CREA), total protein (TP), albumin (ALB), alkaline phosphatase (ALP), alanine aminotransferase (ALT), aspartate aminotransferase (AST), and total bilirubin (TBIL), demonstrating no evidence of renal or hepatic toxicity across doses. Data are presented as mean ± s.d. with individual data points shown. n = 3 (healthy), n = 5 (10 mg/kg), n = 5 (20 mg/kg). Statistical comparisons were performed using one-way ANOVA where applicable.

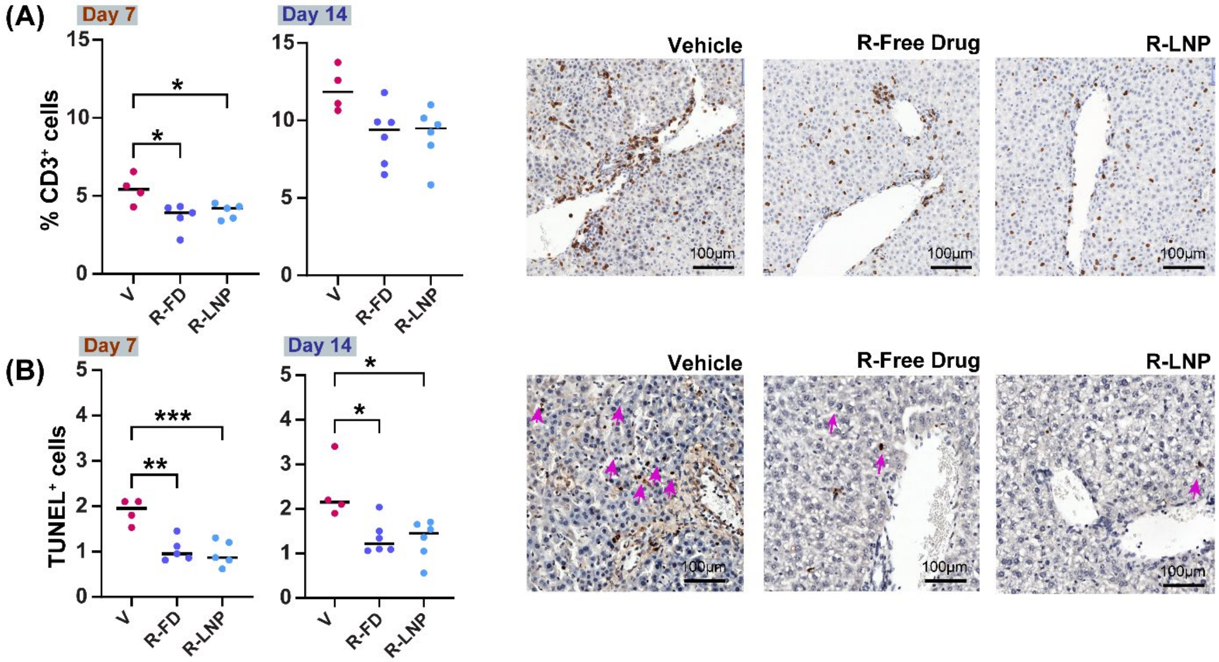


**Supplementary Fig. 24 | Effect of targeted therapy on injury and inflammation in livers of mice in severe GVHD. a**, CD3 positive cells in all groups **b,** TUNEL positive cells V= Vehicle, RFD= Ruxolitinib Free Drug, R-LNP= Ruxolitinib Gal-3 Targeted LNPs (*p<0.05, **p<0.01,***p<0.001, ****p<0.0001, Ordinary one-way ANOVA.)

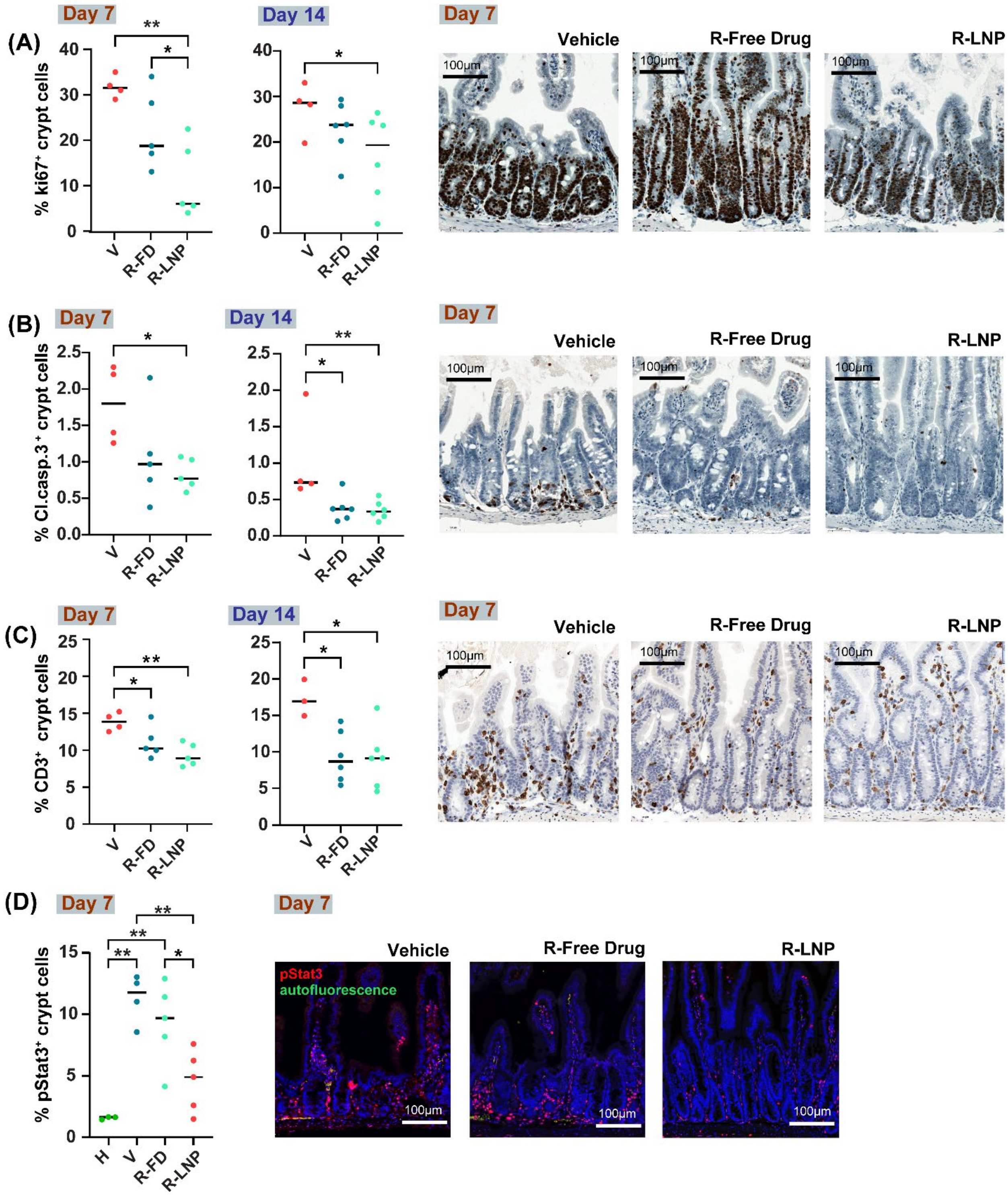


**Supplementary Fig. 25 | Effect of targeted therapy on injury and inflammation in small intestines of mice in severe GVHD a,** Ki67 positive cells in small intestine crypts **b,** Cleaved caspase 3 positive cells in small intestine crypts **c,** CD3 positive cells on the endothelium of small intestine crypts **d**, pSTAT3 positive cells in small intestine crypts **).** V= Vehicle, RFD= Ruxolitinib Free Drug, R-LNP= Ruxolitinib Gal-3 Targeted LNPs (*p<0.05, **p<0.01,***p<0.001, ****p<0.0001, Ordinary one-way ANOVA.)

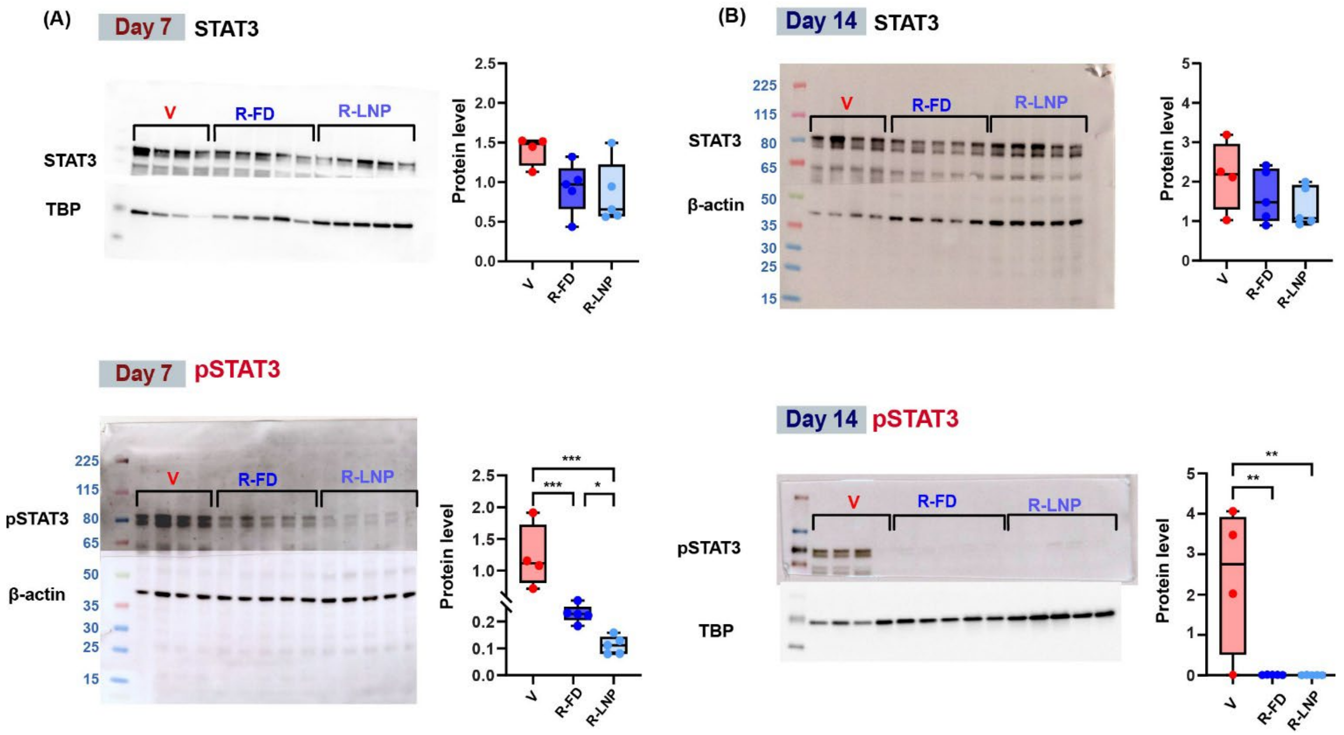


**Supplementary Fig. 26 | The effect of free or LNP-encapsulated Ruxolitinib on STAT3 and p-STAT3 levels in severe GVHD (2M), as measured by western blot on Day 7 (A) or day 14 post BMT (B).** (*p<0.05, **p<0.01,***p<0.001, ****p<0.0001, Ordinary one-way ANOVA.)

| Protein Name | Primary Antibodies | Code | Company | Dilution |
|---|---|---|---|---|
| **STAT3** | STAT3 (79D7) Rabbit mAb | 4904T | Cell Signaling Technology | 1:1000 |
| **pSTAT3** | Phospho STAT3 (Tyr705) (D3A7) XP® Rabbit mAb | 9145S | Cell Signaling Technology | 1:1000 |
| **STAT1** | STAT1 (D1K9Y) Rabbit mAb | 14994T | Cell Signaling Technology | 1:1000 |
| **pSTAT1** | Phospho STAT1 (Ser727) (D3B7) Rabbit mAb | 8826S | Cell Signaling Technology | 1:1000 |
| **pNF-κB** | Anti-NF-κB p65 Rabbit Monoclonal Antibody | 8242S | Cell Signaling Technology | 1:1000 |
| **SOCS2** | SOCS2 Antibody | 2779T | Cell Signaling Technology | 1:1000 |
| **RELB** | RelB (D7D7W) Rabbit mAb | 10544 | Cell Signaling Technology | 1:1000 |
| **GAPDH** | Gapdh (14C10) Rabbit mAb | 2118S | Cell Signaling Technology | 1:1000 |

**Supplementary Table 5: Antibodies used for Western Blot (WB)**

| Drug | Absorbance (nm) | Reference (nm) | Retention time (min) |
|---|---|---|---|
| Ruxolitinib | 230 | 550 | 3.02 |
| WP1066 | 312 | 360 | 4.3 |
| Tofacitinib | 224 | 550 | 2.467 |
| TPCA-1 | 312 | 550 | 3.315 |

**Supplementary Table 6: Absorbance and retention time of drugs in HPLC**

| Antigen | Product | Antibody type | Source | Dilution/ concentration |
|---|---|---|---|---|
| **Tbet** | T-bet/TBX21 (E4I2K) | Rabbit monoclonal | Cell Signaling, 97135 | 0.35 µg/mL |
| **Ly6G** | Ly-6G (E6Z1) | Rabbit monoclonal | Cell Signaling, 87048 | 0.1 µg/mL |
| **CD4** | Anti-CD4 [EPR19514] | Rabbit monoclonal | Abcam, 183685 | 0.5 µg/mL |
| **CD8** | Anti-CD8 alpha [EPR20305] | Rabbit monoclonal | Abcam, 209775 | 0.1 µg/mL |
| **B220** | BD Pharmingen™ Biotin CD45R/B220 | Rat monoclonal | BD Bioscience, 553086 | 0.5 µg/mL |

**Supplementary Table 7: Mouse multiplex primary antibodies in sequential order**

| Antigen | Product | Antibody type | Source | Dilution/concentration |
|---|---|---|---|---|
| **Aquaporin2** | Anti-Aquaporin 2 [EPR21080] | Rabbit monoclonal | Abcam, 199975 | 1:4000 dil |
| **galectin-3** | Invitrogen Galectin 3 (eBioM3/38 (M3/38)) | Rat monoclonal | ThermoFisher Scientific, 14-5301-82 | 1:16000 dil/0.03 µg/mL |
| **Calbindin** | Invitrogen Calbindin D28K (JF05-01) | Rabbit monoclonal | ThermoFisher Scientific, MA5-41211 | 1:128000 dil/ 0.008 µg/mL |
| **CD31** | Anti-CD31 [EPR17259] | Rabbit monoclonal | Abcam, 182981 | 0.08 µg/mL |
| **CD3** | Ventana anti-CD3 (2GV6) | Rabbit monoclonal | Roche, 790-4341 | 1:5 dil |
| **Aquaporin1** | Aquaporin 1 (JM10-98) | Rabbit monoclonal | ThermoFisher Scientific, MA5-32593 | 1:800 dil |
| **p-selectin** | Anti-CD62P antibody | Rabbit Monoclonal | Abcam AB255822 | 0.6µg/ml |
| **CD8** | Anti-CD8 | Rabbit Monoclonal | Ventana 790-4460 | 1/40 dil |
| **Tbet** | T-bet/TBX21 (D6N8B) | Rabbit Monoclonal | CST13232 | 1:100 dil / 0.27 µg/ml |
| **CD69** | Anti-CD69 [EPR21814] | Rabbit Monoclonal | Abcam, ab323396 | 1:500 dil |
| **GATA3** | Anti-GATA3 antibody [EPR16651] - ChIP Grade | Rabbit Monoclonal | Abcam, ab199428 | 2.25 µg/ml |
| **GZB** | Granzyme B Rabbit Polyclonal Antibody | Rabbit Polyclonal | Roche/ Ventana, 760-4283 | 1:4 dil |

**Supplementary Table 8: Human multiplex primary antibodies in sequential order and p-selectin**

#### REFERENCES


1. Clark PR, Kim RK, Pober JS, Kluger MS. Tumor necrosis factor disrupts claudin-5 endothelial tight junction barriers in two distinct NF-kappaB-dependent phases. *PLoS One* **10**, e0120075 (2015).
2. Mehta D, Ravindran K, Kuebler WM. Novel regulators of endothelial barrier function. *Am J Physiol Lung Cell Mol Physiol* **307**, L924-935 (2014).
3. Higo S*, et al.* Acute graft-versus-host disease of the kidney in allogeneic rat bone marrow transplantation. *PLoS One* **9**, e115399 (2014).
4. Wettschureck N, Strilic B, Offermanns S. Passing the vascular barrier: endothelial signaling processes controlling extravasation. *Physiological reviews* **99**, 1467-1525 (2019).
5. Nong J, Glassman PM, Muzykantov VR. Targeting vascular inflammation through emerging methods and drug carriers. *Advanced drug delivery reviews* **184**, 114180 (2022).
6. Simmons S, Erfinanda L, Bartz C, Kuebler WM. Novel mechanisms regulating endothelial barrier function in the pulmonary microcirculation. *The Journal of physiology* **597**, 997-1021 (2019).
7. Scott RP, Quaggin SE. The cell biology of renal filtration. *Journal of cell biology* **209**, 199-210 (2015).
8. Claesson-Welsh L, Dejana E, McDonald DM. Permeability of the endothelial barrier: identifying and reconciling controversies. *Trends in molecular medicine* **27**, 314-331 (2021).
9. Papadimitriou JC, Drachenberg CB, Kleiner D, Choudhri N, Haririan A, Cebotaru V. Tubular epithelial and peritubular capillary endothelial injury in COVID-19 AKI. *Kidney International Reports* **6**, 518-525 (2021).
10. Wilson NA*, et al.* An in vitro model of antibody-mediated injury to glomerular endothelial cells: Upregulation of MHC class II and adhesion molecules. *Transplant immunology* **58**, 101261 (2020).
11. Lakshminarayan R*, et al.* Galectin-3 drives glycosphingolipid-dependent biogenesis of clathrin-independent carriers. *Nature cell biology* **16**, 592-603 (2014).
12. Ivashenka A*, et al.* Glycolipid-dependent and lectin-driven transcytosis in mouse enterocytes. *Communications biology* **4**, 173 (2021).
